# ELOVL6 as a Therapeutic Target: Disrupting c-MYC-Driven Lipid Metabolism to Enhance Chemotherapy in Pancreatic Cancer

**DOI:** 10.1101/2024.11.11.622928

**Authors:** Ana García García, María Ferrer Aporta, Germán Vallejo Palma, Antonio Giráldez Trujillo, Raquel Castillo-González, David Calzón Lozano, Alberto Mora Perdiguero, Raúl Muñoz Velasco, Miguel Colina Castro, Elena de Simone Benito, Raul Torres-Ruiz, Sandra Rodriguez-Perales, Jonas Dehairs, Johannes V Swinnen, Juan Carlos Garcia-Cañaveras, Agustín Lahoz, Sandra Montalvo Quirós, Carlos del Pozo-Rojas, Clara Luque Rioja, Francisco Monroy, Diego Herráez-Aguilar, Marina Alonso Riaño, José Luis Rodríguez Peralto, Víctor Javier Sánchez-Arévalo Lobo

## Abstract

Pancreatic ductal adenocarcinoma (PDAC) is a devastating disease, marked by a survival rate of only 12%. Consequently, the exploration of novel therapeutic strategies becomes a critical clinical imperative. Among the genetic alterations contributing to PDAC, c-MYC overexpression arises due to upstream mutations, amplifications, and copy number alterations. c-MYC serves as a key regulator in the tumor’s metabolic reprogramming, playing a pivotal role in proliferation, migration, and metastasis. This study delves into the investigation of the role of the elongase ELOVL6 in c-MYC-induced cell transformation and its potential as a therapeutic target in PDAC. Here, we demonstrate that c-MYC regulates lipid elongation to promote cell transformation, offering a new avenue for therapeutic intervention. Initially, we show the direct regulation of ELOVLs expression by c-MYC in various PDAC mouse models and cell lines, elucidating its upregulation during transformation and tumor progression. Genetic or chemical inhibition of ELOVL6 results in decreased proliferation and migration, accompanied by alterations in fatty acid elongation. These changes in fatty acid composition led to modifications in membrane rigidity, permeability, and thickness, which collectively affect micropinocytosis and macropinocytosis. Importantly, we observe an increase in Abraxane uptake and a synergistic effect when combined with ELOVL6 interference *in vitro*. *In vivo* validation demonstrates that ELOVL6 inhibition significantly reduces tumor growth and enhances the response to Abraxane, thereby increasing overall survival. Altogether, these results position ELOVL6 as a promising therapeutic target in the treatment of PDAC.

## 1. INTRODUCTION

Pancreatic ductal adenocarcinoma (PDAC) is a highly lethal disease, with a survival rate of only 12%^1^. The impact of PDAC is on the rise, projected to become the third leading cause of cancer death by 2025 and the second by 2030 in the Western world^2,3^. Late-stage diagnosis and chemoresistance due to poor vascularization contribute to this poor prognosis^4,5^. Therefore, there is an urgent need to deepen our understanding of the molecular mechanisms underlying PDAC to develop improved therapeutic options.

The c-MYC oncogene, a transcription factor implicated in a third of all human diseases, plays a pivotal role in cancer, influencing processes such as gene transcription, protein translation, cell cycle and metabolism^6^. Notably, c-MYC involvement is crucial in PDAC^7^, where it is overexpressed in 42% of cases, primarily due to KRAS activation, copy number gains and high copy amplification^8^. Despite its significance, targeting c-MYC directly is challenging, given its involvement in numerous cellular processes; hence, identifying downstream effectors becomes imperative.

Lipids assume a vital role in cellular function, contributing to energy balance and maintaining cell integrity through membrane formation. However, during cell transformation, metabolic changes are necessary to meet the altered energy demands^9,10^. In transformed cells, *de novo* lipogenesis is activated, liberating them from external lipid dependence^11,12^. Furthermore, glycolysis and glutaminolysis are enriched in tumor cells, both processes regulated by c-MYC, which virtually controls genes in glycolysis and many in glutaminolysis^13,14^.

In PDAC, MYC collaborates with KRAS and HIF1A in metabolic reprogramming, directing glutamine carbons to the Krebs cycle^15–17^. Acetyl-CoA, crucial for lipid synthesis, is generated through ATP citrate lyase (ACLY) and subsequently transformed by acetyl-CoA carboxylase (ACCA) to malonyl-CoA, a step enhanced in PDAC^18,19^. Fatty acid synthase (FASN) then utilizes these components, including NADPH, to generate fatty acids, with MYC inducing the expression of enzymes involved^20^.

Detailed lipidomic analyses have implicated ELOVL6 as a significant contributor to the elongation process in lung SCC, with its inhibition markedly impairing tumor growth and cell proliferation in lung SCC and hepatic cancer^21^. The inhibition of ELOVL6 by various microRNAs has been shown to curtail cell proliferation in glioblastoma, underscoring its potential as a therapeutic target^22^. This study sheds light on the mechanism by which c-MYC regulates lipid metabolism in pancreatic ductal adenocarcinoma (PDAC), with a particular focus on the expression of fatty acid elongases (ELOVLs). It identifies ELOVL6 as a promising target for therapeutic intervention in pancreatic cancer. Silencing or inhibiting ELOVL6 in PDAC cells impedes tumor growth *in vitro* and *in vivo*, altering membrane composition and thickness, thus enhancing permeability to therapeutic agents. These findings underscore the therapeutic potential of ELOVL6 inhibition, either alone or in combination with Abraxane, in treating PDAC.

## 2. RESULTS

### 2.1. c-Myc directly regulates the expression of different elongases

In a c-Myc-dependent pancreatic cancer mouse model (*Ela1-Myc* mice), we elucidated c-Myc’s pivotal role in maintaining the acinar phenotype and driving transformation^23^. This transgenic model, where c-Myc is under the control of the Elastase (*Ela1*) promoter, enables c-Myc overexpression in the acinar compartment.

Leveraging RNA-seq data from 8-week-old Ela1-Myc pancreata, we performed Gene Set Enrichment Analysis (GSEA) on the Hallmark gene sets, which reinforced the significant enrichment of pathways related to proliferation and oncogenic signaling (**Supplementary File 1**). Additionally, we conducted a comprehensive transcriptomic analysis, revealing upregulated genes associated with E2F targets, cell cycle, DNA replication, and Myc targets, consistent with our expectations (**Supplementary Fig. 1a**). GSEA analyses further affirmed the enrichment of “Myc targets” and “cell cycle” pathways (**Supplementary Fig. 1b and 1c**).

To investigate c-Myc’s impact on lipid metabolism, we curated an analysis of c-Myc target genes identified by *Gouw et al*.^20^. Notably, *Ela1-Myc* mice exhibited overexpression of *Fasn, Scd1, Acly, Acaca*, and various *Elovls* compared to their wild-type counterparts **(Fig. 1a and 1b)**. Our analysis of ChIP-seq data from the *Ela1-Myc* model demonstrated c-Myc occupancy in the promoters of *Elovl1* and *Elovl6* **(Supplementary Fig. 2a)**. We confirmed the direct regulation of *Elovls* by c-Myc using ChIP-seq data available from the KPC mouse model (*Pdx1-cre; LSL-Kras^G12D/+^; LSL-p53^R172H/+^*), in which oncogenic *Kras* and mutant *p53* are expressed in pancreatic progenitor cells^24^ **(Supplementary Fig. 2b)**. Furthermore, by utilizing the peaks situated in the transcription start site (TSS) of the KPC mouse model, we performed a functional enrichment analysis and identified gene ontology processes related with lipid metabolism and long chain fatty acid metabolism among others (**Supplementary Fig. 2c)**. Additionally, we validated this regulation in an independent model, employing RNA-seq and ChIP-seq data from mouse embryonic fibroblasts (MEFs) infected with the c-Myc-ER inducible system^25^. This analysis revealed the upregulation of all elongases upon c-Myc induction, with c-Myc binding to the promoters of different elongases **(Supplementary Fig. 3a, 3c and 3c)**. Collectively these results support for the first time the direct regulation of the different elongases by the c-Myc oncogene in a context dependent manner.

**Fig. 1.**
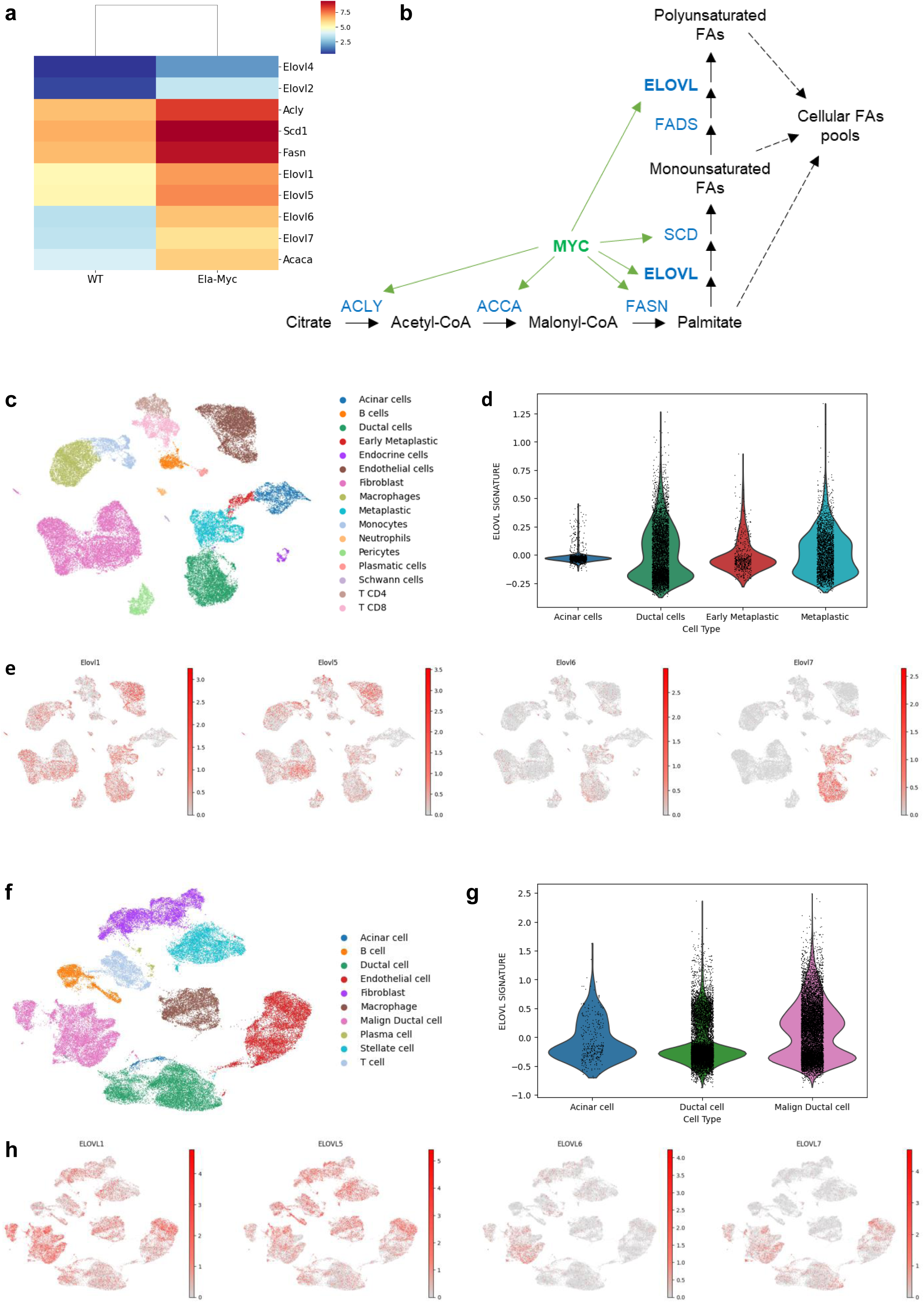
*ELOVLs* are over-expressed in pancreatic ductal adenocarcinoma. **a** Heatmap showing overexpressed FAs synthesis genes in *Ela1-Myc* mouse model; RNA-seq data; scale bar shows the mean (by gene for each condition) of previously scaled count data. **b** Overview of MYC-regulated genes in the FAs synthesis metabolic pathway based on the RNA-seq data shown in (**1a**). **c** UMAP of scRNA-seq data from PDAC tissue of PRT mouse model revealing the cell type composition present in the dataset. **d** Violin plot representing *Elovls* general expression in PDAC relevant cells; *Elovls* signature is comprised of murine *Elovl1*, *Elovl5*, *Elovl6* and *Elovl7*; the score represents average expression of this set of genes in acinar, ductal, early, and late metaplastic cells against a genome-wide reference. **e** UMAPs representing the expression per cell of *Elovl1*, *Elovl5*, *Elovl6* and *Elovl7*; UMAP topology and cell types match the ones shown in (**1c**); high expression level is marked as red, and low expression level as gray; scale bars show normalized (10^4^ counts per cell) and log-transformed counts. **f** UMAP of scRNA-seq data from 24 human PDAC samples revealing the cell type composition present in the dataset. **g** Violin plot representing *ELOVLs* general expression on PDAC relevant cells; *ELOVLs* signature is comprised of human *ELOVL1*, *ELOVL5*, *ELOVL6* and *ELOVL7*; the score represents average expression of this set of genes in acinar, ductal, early and late metaplastic cells against a genome-wide reference. **h** UMAPs representing the expression per cell of *ELOVL1*, *ELOVL5*, *ELOVL6* and *ELOVL7*; UMAP topology and cell types match the ones shown in (**1f**); high expression level is marked as red, and lox expression level as gray; scale bars show normalized (10^4^ counts per cell) and log-transformed counts.

### 2.2. *ELOVLs* are over-expressed during tumor progression

To study the role of Elovls during acinar-to-ductal metaplasia (ADM) and transformation, we analyzed scRNA-seq data from a *Ptf1a-CreER, LSL-Kras-G12D, LSL-tdTomato* (PRT) mouse model conducted by *Schlesinger et al.*^26^. The presence of the *LSL-tdTomato* reporter allowed lineage tracing of acinar cells, revealing metaplastic acinar cells expressing *Krt19* but not *Cpa1* at late time points **(Supplementary Fig. 4a)**. The analysis of this dataset revealed the overexpression of *Elovl1, Elovl5, Elovl6*, and *Elovl7* and this signature was enriched in the tumor cells **(Fig. 1c and 1d, and Supplementary Fig. 4b)**. These findings strongly support the involvement of Elovls in tumor progression from acinar to full metaplastic cells. The expression patterns of *Elovl1* and *Elovl5* were characterized by a diffuse distribution across various cell types, suggesting their widespread presence. In contrast, *Elovl6* and *Elovl7* exhibited a more distinct and specific expression localized within the tumor compartment. This observation highlights the differential roles of these elongase enzymes in various cell types and underscores the potential significance of *Elovl6* and *Elovl7* in the context of the tumor microenvironment **(Fig. 1e)**. To validate these results, we decided to explore dataset available from the *Eµ-Myc* mouse model, which allow to study the progression of the disease from pre-tumoral to tumoral stages^25^. We investigated Myc genomic distribution and *Elovls* expression during B-cell lymphomagenesis *in vivo*. We analyzed the transcriptomic profile coupled with ChIP-seq in B cells from young non-transgenic (control, C) and *Eµ-Myc* transgenic littermates (pre-tumor, P), and in lymphomas arising in adult *Eµ-Myc* animals (tumor, T). Consistent with progressive increases in Myc mRNA, we observed the increase in *Elovl1, Elovl5* and *Elovl6* **(Supplementary Fig. 5a)**. We analyzed c-Myc genomic distribution, and we found it in the promoter of *Elovl1, Elovl4, Elovl5, Elovl6* and *Elovl7* **(Supplementary Fig. 5b)**. So, we can conclude that c-Myc induces the expression of several *Elovls* by directly recruiting them to their promoters, and this induction is heightened during cellular transformation and tumor progression.

### 2.3. ELOLV6 is the only elongase restricted to the tumor compartment in human PDAC

We chose to investigate the expression of *ELOVLs* in the PDAC TCGA cohort using the GEPIA browser. Notably, *ELOVL1, ELOVL5*, and *ELOVL6* exhibited significant overexpression in pancreatic ductal adenocarcinoma (PDAC) compared to normal tissue, with *ELOVL6* being the most prominently overexpressed **(Supplementary Fig. 6a, 6c, and 6c)**. This pattern persisted across various tumoral stages **(Supplementary Fig. 6d)**, making PDAC one of the few tumors in which *ELOVL6* is overexpressed **(Supplementary Fig. 6e)**.

Given these findings, we sought to validate our results using a distinct cohort analyzed through scRNA-seq consisting of 24 PDAC tumor samples and 11 control pancreases without any treatment^27^. Our observations confirmed the upregulation of *ELOVL1, ELOVL5, ELOVL6* and *ELOVL7*, thus corroborating the results obtained in the TCGA cohort **(Supplementary Fig. 4c)**. Importantly, this signature was found to be overexpressed in tumoral cells compared to acinar and ductal cells **(Fig. 1f and 1g)**. Utilizing UMAP, we observed that *ELOVL6* was the only enzyme restricted to the tumor compartment **(Fig. 1h)**.

We can conclude that PDAC revealed significant overexpression of *ELOVL1, ELOVL5*, and notably, *ELOVL6*, with a specific localization to the tumor compartment, emphasizing its potential relevance in PDAC.

### 2.4. c-MYC directly regulates *ELOVL6* expression in PDAC cell lines

Based on the specific expression pattern of *ELOVL6* we decided to describe its role in pancreatic cancer. Therefore, we performed RT-qPCR and western blot in a panel of five different PDAC cell lines (**Supplementary Fig. 7a**) and selected three of them, based on their *c-MYC* expression levels. We observed that T3M4 expressed high levels of *c-MYC*, Patu 8988T exhibited intermediate levels, and Panc1 showed low levels of *c-MYC* **(Fig. 2a**, **Fig. 2b and Supplementary Fig. 7b)**. Interestingly, upon analyzing the expression levels of *ELOVL6* in these cell lines, we observed a parallel pattern to *c-MYC* expression **(Fig. 2c and 2d)**. To assess the correlation between both genes, we chose to downregulate *c-MYC* in T3M4 (the PDAC line expressing the highest *c-MYC* levels) using shRNA (shMYC). We observed that a reduction in *c-MYC* levels was associated with a downregulation in the expression of all *ELOVL* family members **(Fig. 2e)**, resulting in a subsequent decrease in ELOVL6 protein levels **(Fig. 2f)**. Conversely, upon upregulating c-MYC in Panc1 (the PDAC line expressing the lowest c-MYC levels), we noted an upregulation all the ELOVLs **(Fig. 2g)**, resulting in a subsequent elevation of ELOVL6 protein levels **(Fig. 2h)**. Finally, we explored the recruitment of c-MYC to the *ELOVL6* promoter through chromatin immunoprecipitation and RT-qPCR (ChIP-qPCR), affirmatively confirming the binding of c-MYC to this specific region **(Fig. 2i and 2j)**.

**Fig. 2.**
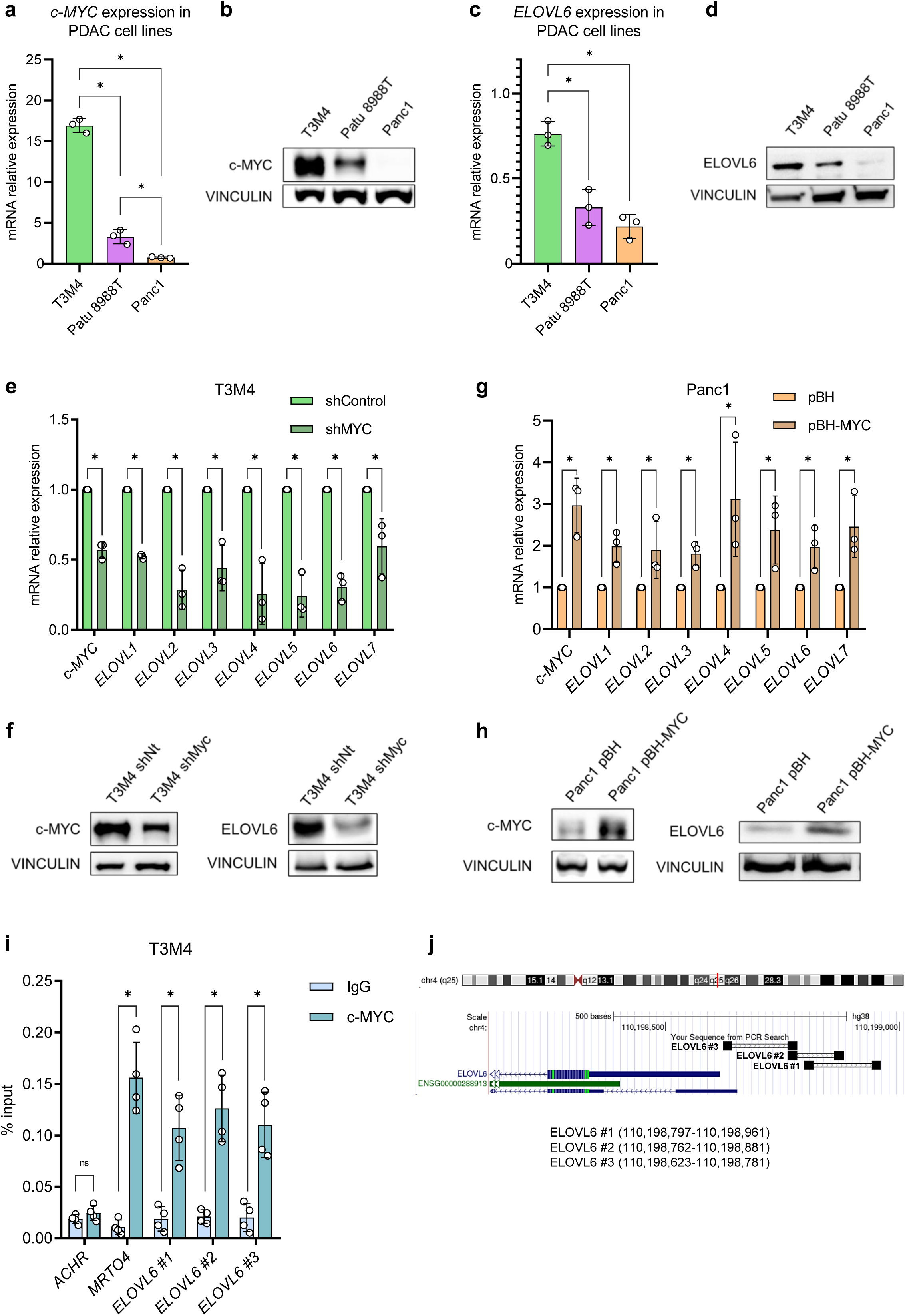
c-MYC directly regulates *ELOVL6* expression in PDAC cell lines. **a** mRNA expression of *c-MYC* in PDAC cell lines; gene expression is normalized to *HPRT*; n = 3 per cell line. **b** Western blot of c-MYC in PDAC cell lines. **c** mRNA expression of *ELOVL6* in PDAC cell lines; gene expression is normalized to *HPRT*; n = 3 per cell line. **d** Western blot of ELOVL6 in PDAC cell lines. **e** mRNA expression of *c-MYC* and *ELOVLs* in T3M4 (shControl and shMYC); gene expression is normalized to *HPRT* and shControl; n = 3 per genotype. **f** Western blot of c-MYC and ELOVL6 in T3M4 (shControl and shMYC). **g** mRNA expression of *c-MYC* and *ELOVLs* in Panc1 (pBH and pBH-MYC); gene expression is normalized to *HPRT* and pBH; n = 3 per genotype. **h** Western blot of c-MYC and ELOVL6 in Panc1 (pBH and pBH-MYC). **i** Chromatin immunoprecipitation of c-MYC on the *ELOVL6* promoter; n = 4. **j** Primers’ hybridization scheme on ELOVL6 promoter. All data are presented as mean ± SD; ns: not statistically significant, *p ≤ 0.05; (a, c, e, g, i) Mann-Whitney test. Source data are provided as a Source Data file.

These results collectively unveil a robust correlation between c-MYC and ELOVLs, particularly emphasizing the unique expression of *ELOVL6* in PDAC tumors. Additionally, our results conclusively demonstrate c-MYC’s direct regulatory influence on ELOVL6.

### 2.5. ELOVL6 interference decreases cell proliferation and migration *in vitro*

To establish the essential role of ELOVL6 in cell proliferation within human PDAC tumors, we interfered ELOVL6 using two different approaches in T3M4 and Patu 8988T cell lines. Firstly, we downregulated ELOVL6 levels using two distinct shRNAs, denoted as shELOVL6 #1 and shELOVL6 #2 **(Fig. 3a)**, leading to a concurrent decrease in ELOVL6 protein levels **(Fig. 3b)**. Additionally, we employed the chemical inhibitor ELOVL6-IN-2. Subsequent analysis of cell proliferation and colony growth revealed a consistent reduction in cell proliferation across the tested cell lines following ELOVL6 interference, irrespective of the method used **(Fig. 3c and 3d)**. To determine the specific effects of ELOVL6-IN-2 on PDAC cells, we designed an ELOVL6 knockout (KO) that resulted in the complete ablation of ELOVL6 **(Supplementary Fig. 8a)**. As expected, the proliferation and formation of colonies by T3M4 ELOVL6 KO cells was reduced, and we did not observe an additive effect when treating these cells with ELOVL6-IN-2 **(Supplementary Fig. 8b)**. This diminished proliferation corresponded with a noticeable accumulation of cells in the G1 phase of the cell cycle **(Fig. 3e and 3f)**, with no changes in the number of apoptotic cells upon ELOVL6 silencing or inhibition **(Supplementary Fig. 9a and 9b)**.

**Fig. 3.**
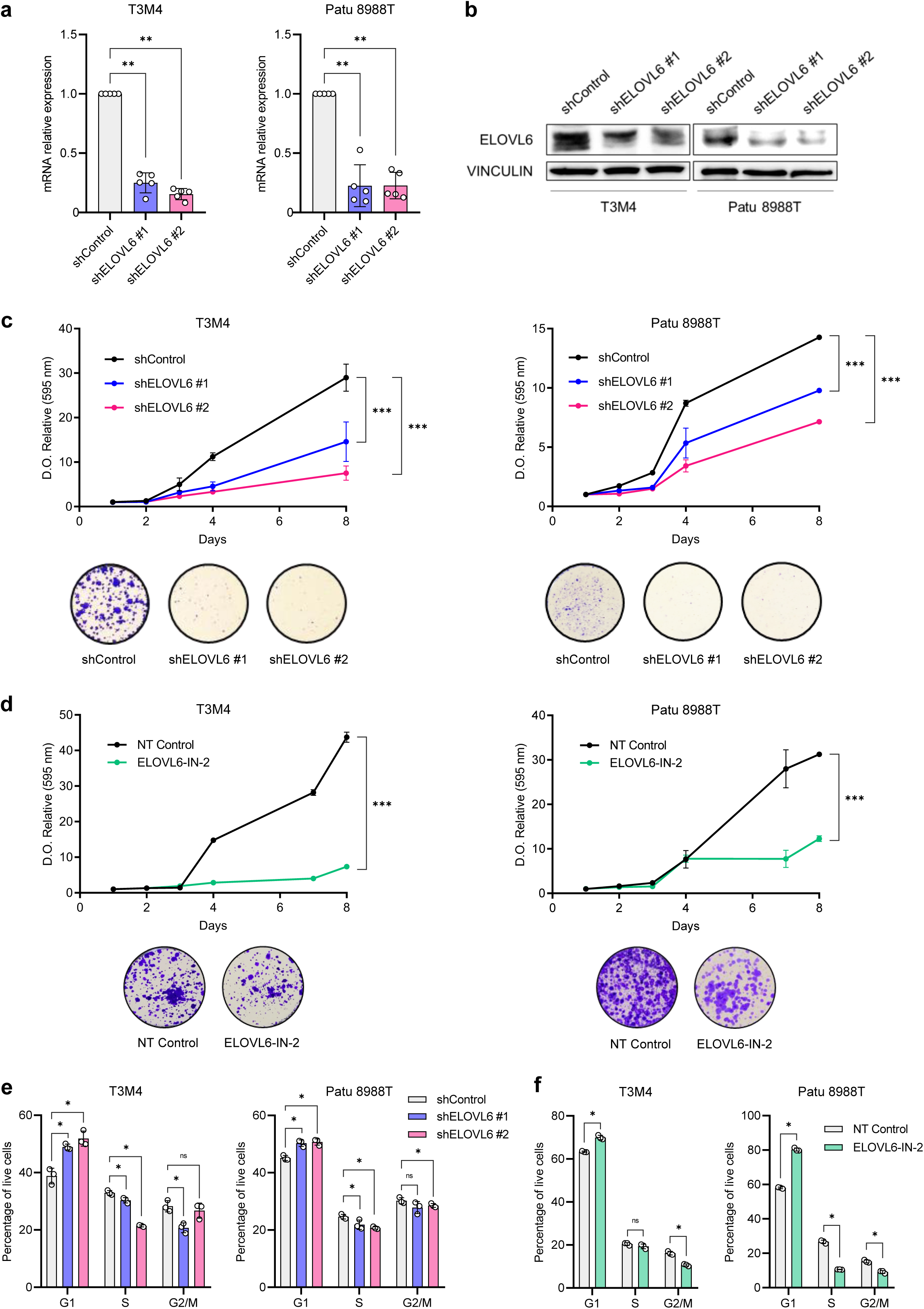
ELOVL6 interference decreases cell proliferation *in vitro*. **a** mRNA expression showing *ELOVL6* silencing in PDAC cell lines; gene expression is normalized to *HPRT* and shControl; n = 5 per genotype and cell line. **b** Western blot showing *ELOVL6* silencing in PDAC cell lines. **c** Proliferation and colony assays of *ELOVL6*-silenced cells compared to non-targeted control in PDAC cell lines; n = 3. **d** Proliferation and colony assays of ELOVL6-inhibited cells compared to non-treated control in PDAC cell lines; n = 3. **e** Cell cycle assay by FACS of *ELOVL6*-silenced cells compared to non-targeted control in PDAC cell lines; n = 3. **f** Cell cycle assay by FACS of ELOVL6-inhibited cells compared to non-treated control in PDAC cell lines; n = 3. All data are presented as mean ± SD; ns: not statistically significant, *p ≤ 0.05, **p ≤ 0.01, ***p ≤ 0.001; (a, e, f) Mann-Whitney test; (c, d) Two-way ANOVA test followed by Dunnett test (c) or Sidak test (d). Source data are provided as a Source Data file.

These results were confirmed at the transcriptomic level. Conducting RNA-seq analysis, we explored the alterations in the transcriptome resulting from the inhibition of ELOVL6 using the chemical inhibitor ELOVL6-IN-2. The analysis revealed 202 differentially expressed genes, illustrated in the volcano plot and heatmap **(Fig. 4a and 4b)**, with 142 exhibiting downregulation in the inhibited samples. To elucidate the functional implications of this inhibition, we performed GSEA. The results indicated that untreated cells displayed a higher enrichment in “myc targets” and “cell cycle” pathways, suggesting that inhibition led to cell cycle arrest **(Fig. 4c and 4d)**. Furthermore, signatures associated with lipid metabolism were examined and, once again, they exhibited enrichment in untreated cells, indicating a downregulation of these pathways due to ELOVL6 inhibition **(Fig. 4e and 4f, Supplementary Fig. 10a, 10c and 10c)**.

**Fig. 4.**
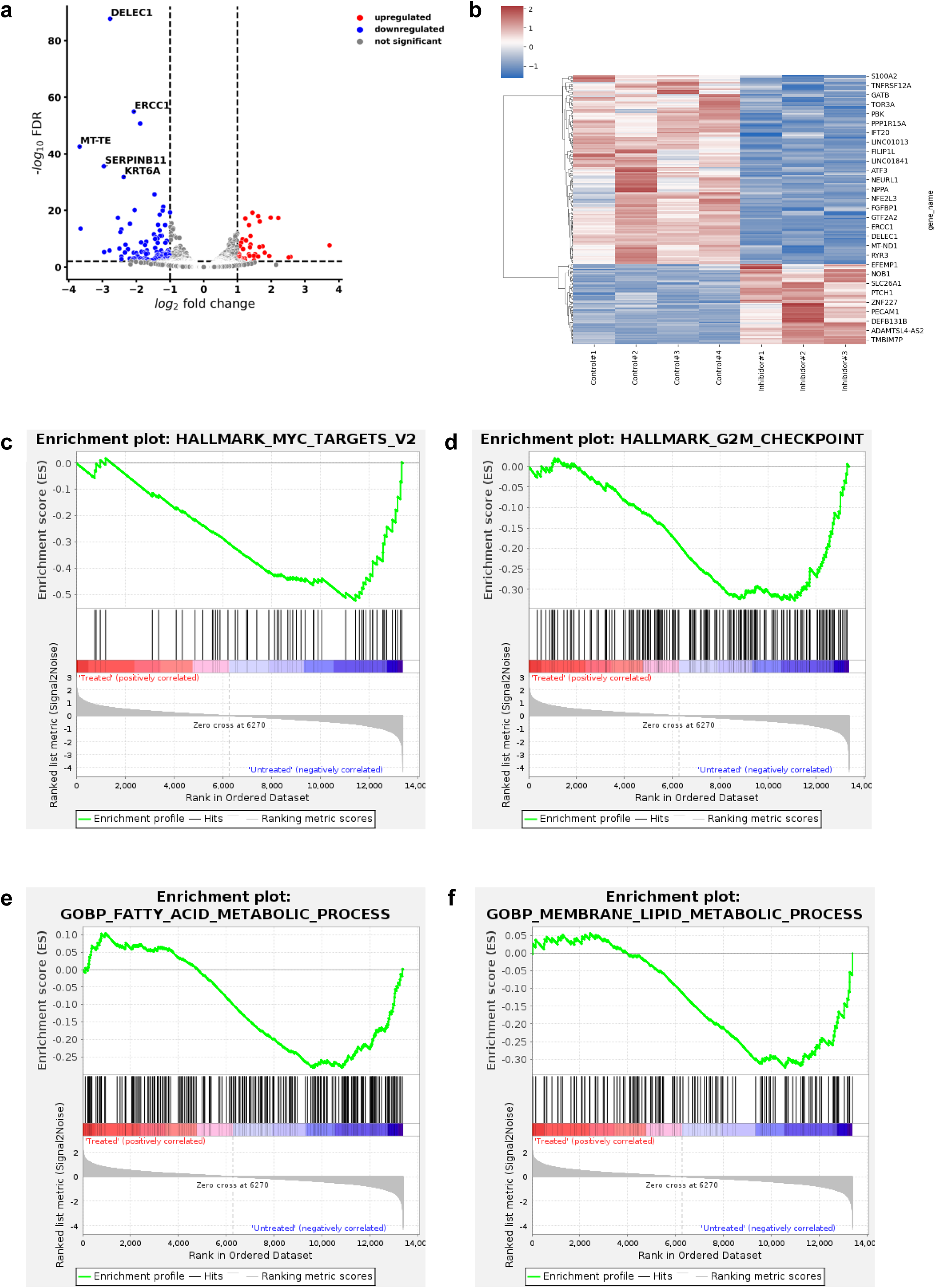
ELOVL6 inhibition causes cell cycle arrest and lipid metabolism downregulation. **a** Volcano plot of DEGs in ELOVL6-inhibited cells compared to non-treated control in T3M4; each dot represents one DEG and colored circles represent DEGs significantly upregulated or downregulated; n = 4. **b** Heatmap of DEGs in (**4a**); scale bar shows DESeq2-normalized and logarithmically transformed gene counts. **c** GSEA of “myc targets hallmarks”. **d** GSEA of “G2M checkpoint hallmarks”. **e** GSEA of “fatty acid metabolic process GOBP”. **f** GSEA of “membrane lipid metabolic process GOBP”.

To validate the RNAseq results functionally and to determine which pathways were affected by ELOVL6 inhibition, we decided to perform a cell signaling phosphor-kinase array. We observed a decrease in the activation of cell proliferation-related pathways such as ERK1/2, Src, WNK1, β-Catenin or RSK1/2/3 **(Supplementary Fig. 11a and 11c)**. To further investigate the pathways affected by ELOVL6 inhibition, we performed a GSEA focusing on cell proliferation and signaling pathways associated with the EGFR **(Supplementary Fig. 11c and 11c)**. This analysis revealed significant downregulation of signatures associated with EGF receptor-related signaling pathways, corroborating our findings from the phospho-kinase array and further supporting the role of ELOVL6 in modulating key signaling mechanisms involved in cell proliferation and growth.

Finally, we investigated the impact of ELOVL6 on cell migration through wound healing and transwell assays. *ELOVL6* downregulation via shRNAs exhibited an impairment in cell migration, as evidenced by delayed wound closure and reduced migration through the transwell **(Supplementary Fig. 12a and 12c)**.

Collectively, these results strongly reinforce the pivotal role of ELOVL6 in both cell proliferation and migration processes, substantiated by transcriptomic analysis and experimental data.

### 2.6. ELOVL6 interference modifies lipid composition

Our hypothesis posited that the observed proliferative and migratory effects were linked to changes in the composition of cell membranes. To investigate this, we conducted a lipidomic analysis focusing on different lipidic species using mass spectrometry in T3M4 genetically or chemically inhibited for ELOVL6. We focused our attention on phosphatidylethanolamine, a phospholipid integral to biological membranes. Remarkably, irrespective of their saturation state, delimited by dashed lines, we noted an accumulation of shorter fatty acids composing phosphatidylethanolamine compared to control counterparts, coupled with a decrease in longer fatty acids. This effect was consistently observed with the interference of ELOVL6 using both shRNAs **(Fig. 5a)** and ELOVL6-IN-2 **(Fig. 5b)**. Additionally, a parallel analysis of phosphatidylcholine, another prevalent phospholipid in biological membranes, yielded similar effects **(Supplementary Fig. 13a and 13c)**. Furthermore, the analysis of total fatty acid composition using isotopically labelled media also confirmed the impact of ELOVL6 interference -either genetically or chemically-on fatty acid metabolism **(Supplementary Fig. 14a and 14c)**. These findings are consistent with the results obtained in the lipidomic analysis.

**Fig. 5.**
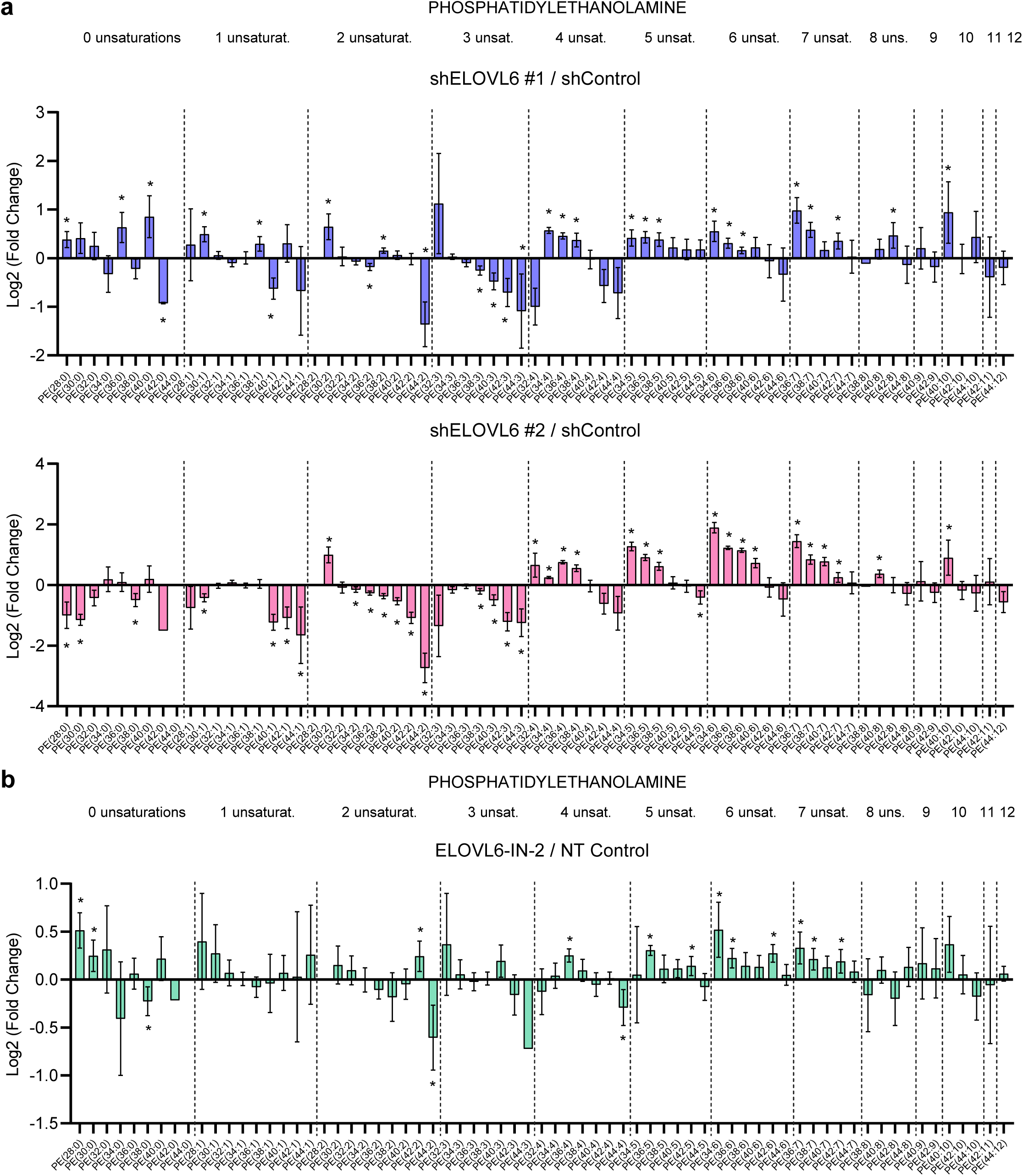
ELOVL6 interference modifies lipid composition. **a** Mass spectrometry lipidomic analysis of phosphatidylethanolamine in *ELOVL6*-silenced cells compared to non-targeted control in T3M4; separated regions represent fatty acids with the same saturation state and carbons’ number increases along each region; n = 3. **b** Mass spectrometry lipidomic analysis of phosphatidylethanolamine in ELOVL6-inhibited cells compared to non-treated control in T3M4; n = 4. Source data are provided as a Source Data file.

To further confirm the results observed upon ELOVL6 interference, we also profiled total fatty acid composition in T3M4 when downregulating its high *c-MYC* expression level. We obtained similar results to ELOVL6 silencing or inhibition **(Supplementary Fig. 14c and 14c)**.

These results leave no doubt that the genetic or chemical inhibition of ELOVL6 exerts a profound influence on the lipidomic landscape within cells. This comprehensive insight underscores the pivotal role of ELOVL6 in orchestrating lipid metabolism and highlights its significance in modulating cell membrane composition.

### 2.7. ELOVL6 interference alters membrane mechanical properties and permeability

We investigated whether the depletion of fatty acids composing phospholipids resulted in a reduction in cell membrane thickness and resistance. To evaluate membrane thickness, we employed transmission electron microscopy (TEM) and observed a notable decrease in membrane thickness in T3M4 and Patu 8988T cells when ELOVL6 was interfered with using shRNAs **(Fig. 6a and 6b).**

**Fig. 6.**
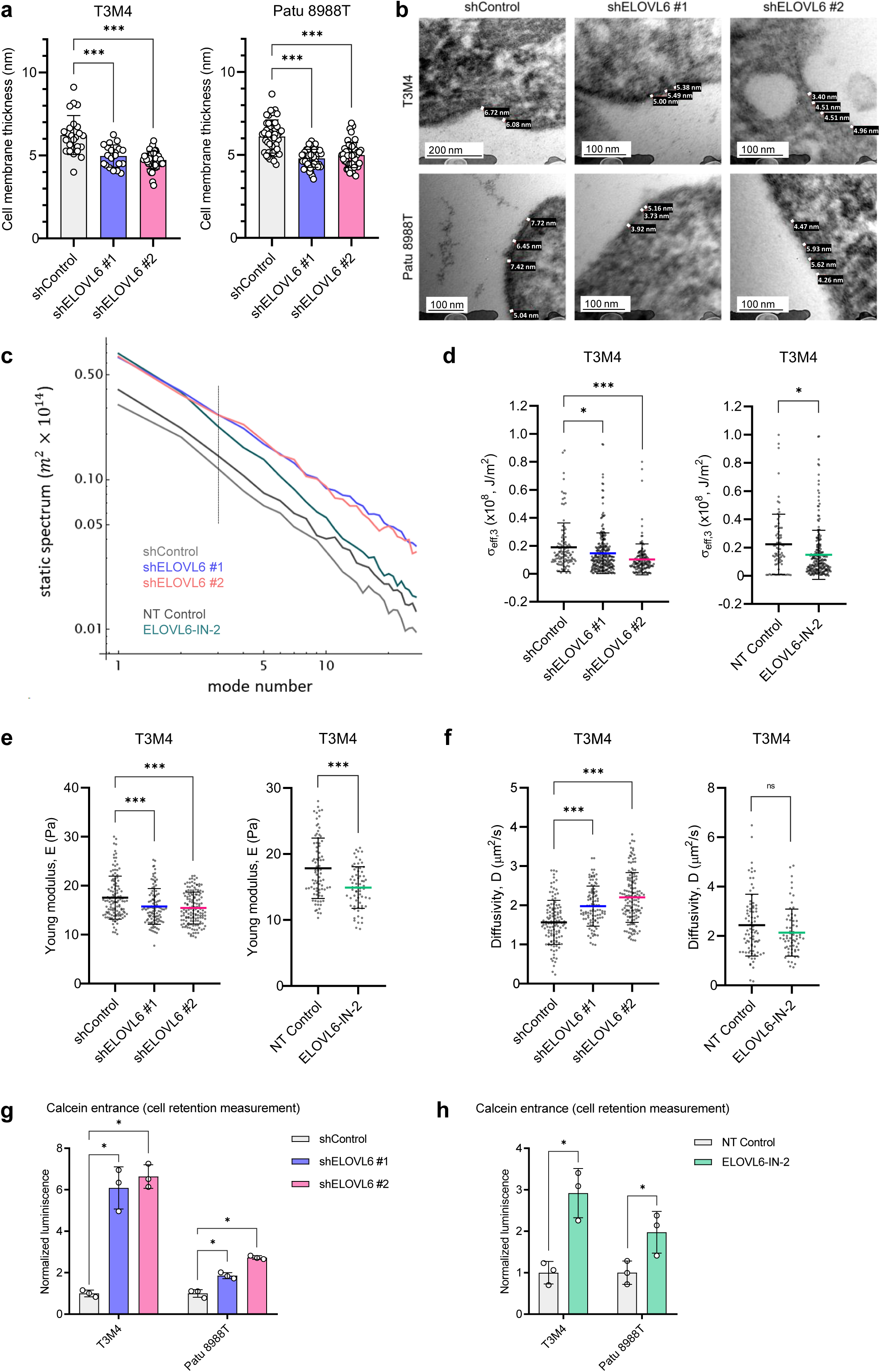
ELOVL6 interference alters membrane mechanical properties and permeability. **a** Cell membrane thickness measurement in transmission electron microscopy images in *ELOVL6*-silenced cells compared to non-targeted control; n = 3 per condition. **b** Representative transmission electron microscopy images of *ELOVL6*-silenced cells compared to non-targeted control. **c** Helfrich-like statics spectra estimated from collective spatial fluctuations for different experimental setups, as a function of the wave vector mode number in log-log scale. **d** Effective surface tension for the wave number value at the tension dominated regime; n = 3. **e** Indentation experiments showing membrane rigidity (E, elasticity, Pa). **f** Indentation experiments showing membrane permeability (D, diffusivity, μm^2^/s). **g** Normalized calcein-AM uptake and processing into calcein in *ELOVL6*-silenced cells compared to non-targeted control in PDAC cell lines; cell retention measurement; n = 3. **h** Normalized calcein-AM uptake and processing into calcein in ELOVL6-inhibited cells compared to non-treated control in PDAC cell lines; cell retention measurement; n = 3. All data are presented as mean ± SD; ns: not statistically significant, *p ≤ 0.05, ***p ≤ 0.001; (a, d, e, f) Ordinary one-way ANOVA test followed by Dunnett test; (d, e, f) Unpaired t test; (g, h) Mann-Whitney test. Source data are provided as a Source Data file.

To estimate cell membrane mechanical modifications, we performed a Fourier decomposition of the equatorial shapes in single-cell experiments and illustrated the resulting static (Helfrich-like) collective spectrum of deformable cellular shapes, representing the dependence of the sample-averaged squared deformations **(Supplementary Fig. 15a)**. The spectra corresponding to ELOVL6 interference, either genetically or chemically, appear above controls, indicating a more flexible behavior and higher variability in cell shape **(Fig. 6c and Supplementary Fig. 15b and 15c)**. By focusing on the third normal mode and applying the previously mentioned relation to each cell, we can extract an effective value for the membrane tension in the experimental groups. Similarly, we observe significant differences between controls and treatments, demonstrating that ELOVL6 interference results in a more flexible behavior **(Fig. 6d)**. Finally, we decided to investigate membrane rigidness against normal stress and permeability under induced cortical deformation using indentation techniques upon ELOVL6 interference either through shRNA downregulation or ELOVL6-IN-2 inhibition. Our findings revealed a significant reduction in membrane rigidity upon ELOVL6 interference, demonstrating a tangible effect on the mechanical properties of the cellular membrane **(Fig. 6e)**. Additionally, we observed a concurrent increase in membrane permeability, indicating a notable shift in the membrane’s functional characteristics although this effect was only evident in the shRNA **(Fig. 6f)**.

To further investigate if all these alterations in membrane composition and mechanical properties correlated with increased membrane permeability, we conducted calcein uptake assays in T3M4 and Patu 8988T. Calcein-acetoxymethyl ester, calcein-AM, is a non-fluorescent compound that enters cell membrane in a passive way, and esterases reduce it to calcein, producing the fluorescent compound **(Supplementary Fig. 16a)**. The results demonstrated a dramatic increase in calcein uptake upon interfering ELOVL6 using shRNAs **(Fig. 6g and Supplementary Fig. 16b)** and ELOVL6-IN-2 **(Fig. 6h and Supplementary Fig. 16c)**. ELOVL6-IN-2 also demonstrated to be ELOVL6-specific when assessing calcein entrance **(Supplementary Fig. 8c)**. Notably, no changes in calcein expulsion were observed **(Supplementary Fig. 16d, 16e and Supplementary Fig. 8d)**, indicating that the increased calcein uptake was not attributed to reduced drug release from the cells but rather to an augmented entrance.

These results suggest a crucial role for ELOVL6 in regulating membrane permeability and mechanical properties, emphasizing its potential implications in modulating cellular processes associated with both rigidity and permeability.

### 2.8. ELOVL6 interference enhances pinocytosis

One of the primary chemotherapeutic drugs for pancreatic ductal adenocarcinoma (PDAC) is paclitaxel. In its formulation as albumin nanoparticles (nab-paclitaxel, marketed under the name Abraxane), paclitaxel enters tumor cells through pinocytosis. Consequently, we examined both micro and macropinocytosis in T3M4 and Patu 8988T using dextran-rhodamine B with molecular weights of 10,000 Da and 70,000 Da, respectively. ELOVL6 interference, either through shRNAs or ELOVL6-IN-2, resulted in an observed increase in micropinocytosis, as indicated by the signal of dextran-rhodamine B 10.000 Da **(Fig. 7a)**. This finding was further supported by immunofluorescence analysis and quantification **(Fig. 7b and 7c)**. Additionally, macropinocytosis exhibited an increase, measured by the signal of dextran-rhodamine B 70,000 Da **(Fig. 7d)**, along with confirmation through immunofluorescence **(Fig. 7e and 7f)**. ELOVL6-IN-2 also demonstrated to be ELOVL6-specific when assessing micro and macropinocytosis **(Supplementary Fig. 8e and 8c)**.

**Fig. 7.**
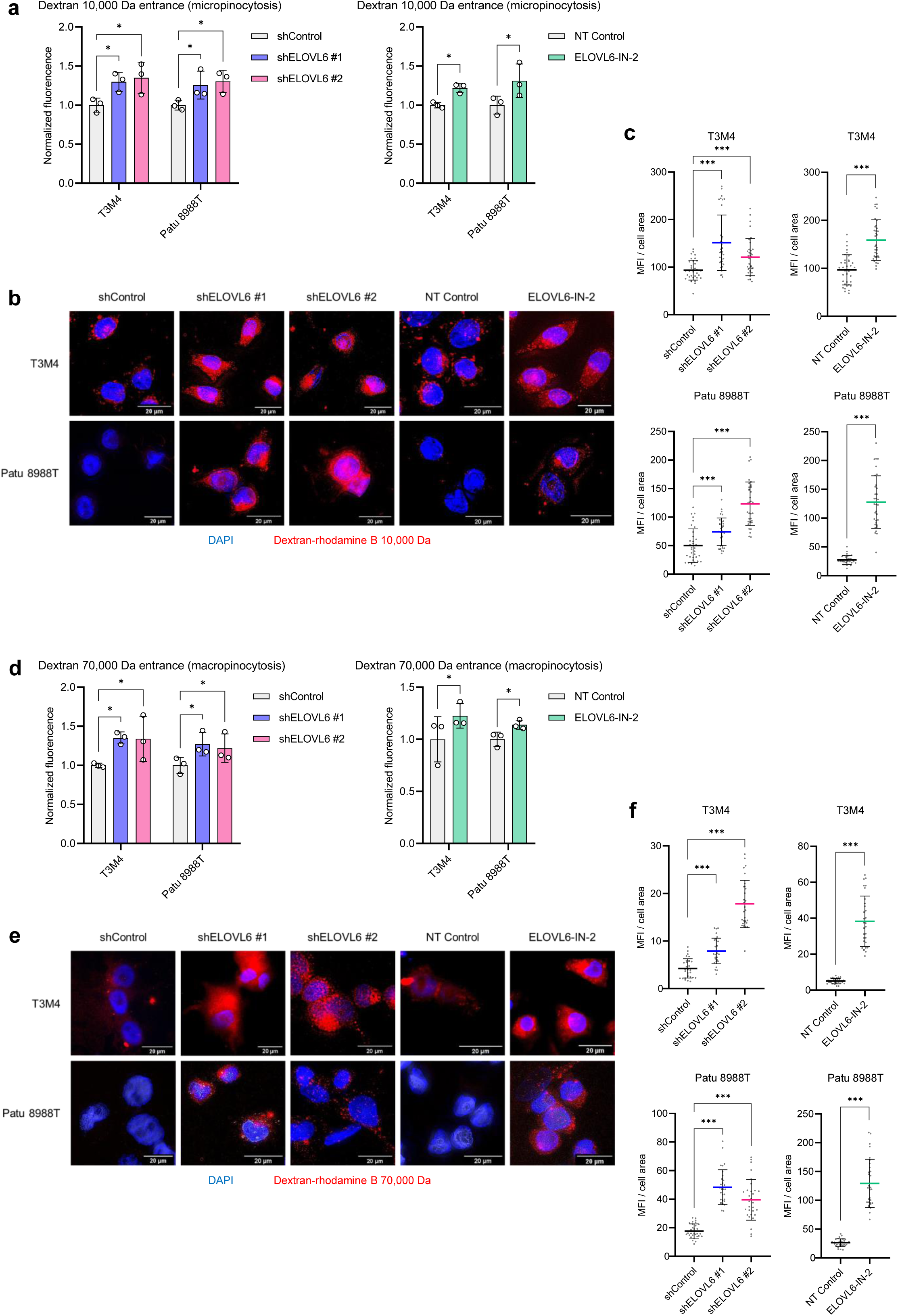
ELOVL6 interference enhances pinocytosis. **a** Micropinocytosis analysis by normalized fluorescence of dextran-rhodamine B 10,000 Da incorporated by *ELOVL6*-silenced cells compared to non-targeted control and ELOVL6-inhibited cells compared to non-treated control in PDAC cell lines; n = 3 per condition. **b** Representative images of immunofluorescence staining for dextran-rhodamine B 10,000 Da (red) and DAPI (blue) on PDAC cell lines. **c** Quantification of micropinocytosis immunofluorescence staining; n = 5 per condition. **d** Macropinocytosis analysis by normalized fluorescence of dextran-rhodamine B 70,000 Da incorporated by *ELOVL6*-silenced cells compared to non-targeted control and ELOVL6-inhibited cells compared to non-treated control in PDAC cell lines; n = 3 per condition. **e** Representative images of immunofluorescence staining for dextran-rhodamine B 70,000 Da (red) and DAPI (blue) on PDAC cell lines. **f** Quantification of macropinocytosis immunofluorescence staining; n = 5 per condition. All data are presented as mean ± SD; *p ≤ 0.05, ***p ≤ 0.001; (a, d) Mann-Whitney test; (c, f) Ordinary one-way ANOVA test followed by Dunnett test; (c, f) Unpaired t test. Source data are provided as a Source Data file.

Hence, these results strongly suggest that ELOVL6 interference exerts a profound impact on fatty acid elongation, ultimately leading to an increased permeability of the cell membrane.

### 2.9. ELOVL6 interference sensitizes to chemotherapy *in vitro*

To evaluate the impact of enhanced permeability, pinocytosis, and changes in membranes properties on Abraxane treatment in PDAC cell lines, we measured the entrance of Flutax-2, a green, fluorescent taxol derivative that binds to microtubules. Strong labelling was observed in T3M4 and Patu 8988T cells when ELOVL6 was interfered with using shRNAs **(Fig. 8a)** and ELOVL6-IN-2 **(Fig. 8b)**. This suggests that ELOVL6 interference enhances the uptake of Abraxane in these PDAC cell lines. Additionally, ELOVL6-IN-2 also demonstrated to be ELOVL6-specific when assessing Flutax-2 entrance **(Supplementary Fig. 8g)**. Next, to explore if ELOVL6 interference could synergize with Abraxane treatment, we calculated the IC50 of Abraxane for T3M4 and Patu 8988T. Both *ELOVL6* silencing by shRNAs and ELOVL6 inhibition by ELOVL6-IN-2 significantly reduced Abraxane IC50s **(Fig. 8c and 8d)**. In aggregate, these findings unveil that ELOVL6 interference synergizes with Abraxane treatment, offering a new therapeutic perspective through the combination of chemotherapy and ELOVL6 inhibition. This synergy holds the potential for increased effectiveness or the utilization of lower Abraxane doses, thereby minimizing off-target effects.

**Fig. 8.**
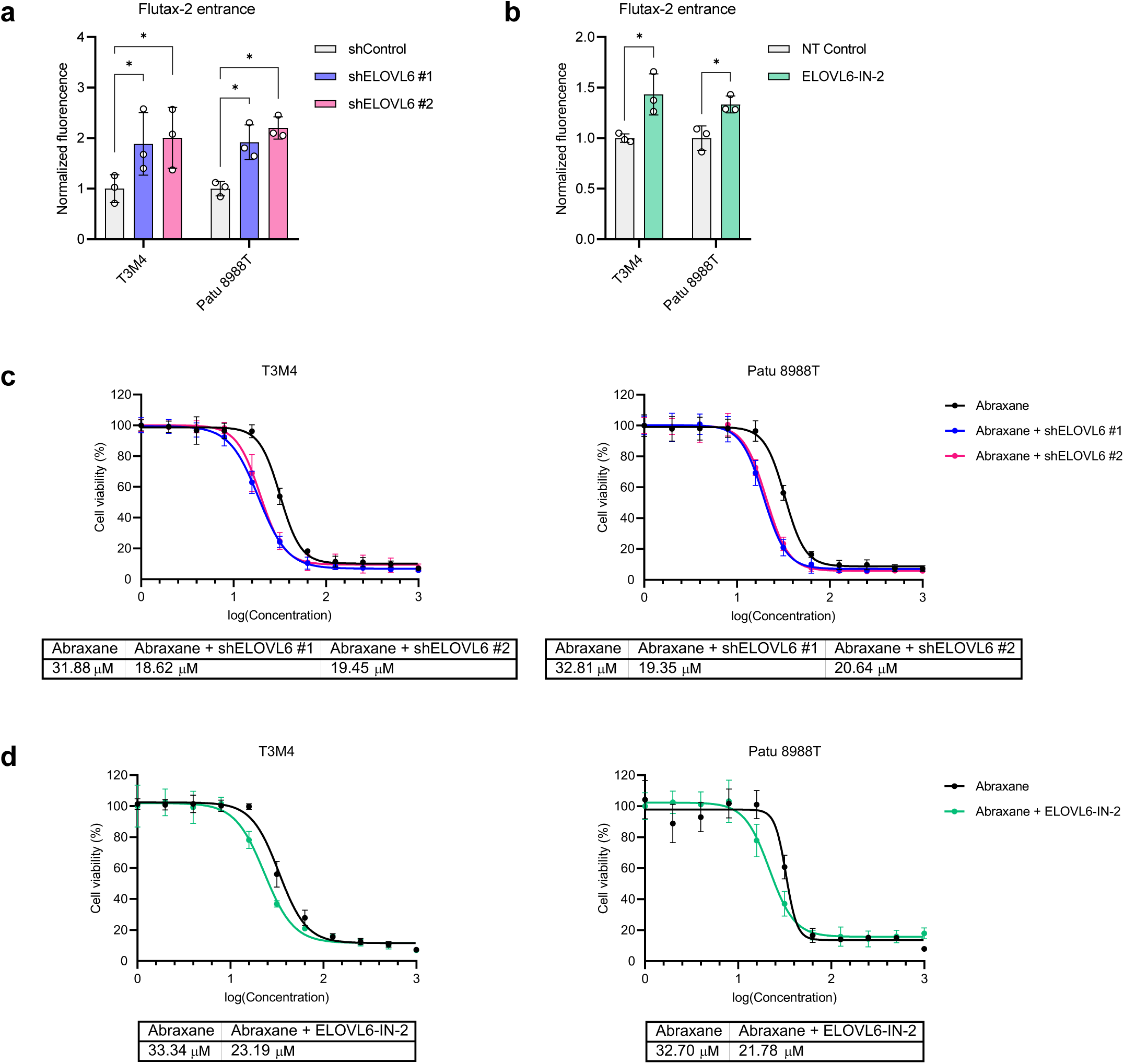
ELOVL6 interference sensitizes to chemotherapy *in vitro.* **a** Normalized Flutax-2 uptake in *ELOVL6*-silenced cells compared to non-targeted control in PDAC cell lines; n = 3. **b** Normalized Flutax-2 uptake in ELOVL6-inhibited cells compared to non-treated control in PDAC cell lines; n = 3. **c** Abraxane IC50 analysis and values using ATPlite in *ELOVL6*-silenced cells compared to non-targeted control in PDAC cell lines; n = 6. **d** Abraxane IC50 analysis and values using ATPlite in ELOVL6-inhibited cells compared to non-treated control in PDAC cell lines; n = 6. All data are presented as mean ± SD; *p ≤ 0.05; (a, b) Mann-Whitney test. Source data are provided as a Source Data file.

### 2.10. ELOVL6 interference efficacy is dependent on *c-MYC* expression levels

When we silenced *ELOVL6* in Panc1 cells, we anticipated several outcomes, including a decrease in both mRNA and protein levels **(Supplementary Fig. 17a and 17c)**, reduced proliferation and colony formation **(Supplementary Fig. 17c)**, an accumulation of cells in the G1 cell cycle phase **(Supplementary Fig. 17d)**, diminished transwell migration **(Supplementary Fig. 17e)**, and impaired wound healing **(Supplementary Fig. 17f)**. However, our assessments of compound uptake revealed that ELOVL6 interference did not enhance cell permeability in a low *c-MYC* expressing context.

To investigate whether this lack of effect was attributed to low c-MYC levels in Panc1, the PDAC cell line with the lowest c-MYC expression, we proceeded to overexpress c-MYC in Panc1 **(Fig. 2g and 2h)** and conducted proliferative analyses. Interestingly, we observed that Panc1-pBH-MYC cells exhibited heightened proliferation and colony formation compared to Panc1 cells. Furthermore, upon ELOVL6 interference using shRNAs **(Supplementary Fig. 18a)** or ELOVL6-IN-2, we observed that Panc1-pBH-MYC ELOVL6-interfered cells experienced a more pronounced proliferative arrest and a complete inability to form colonies compared to their controls **(Fig. 9a and 9b)**.

**Fig. 9.**
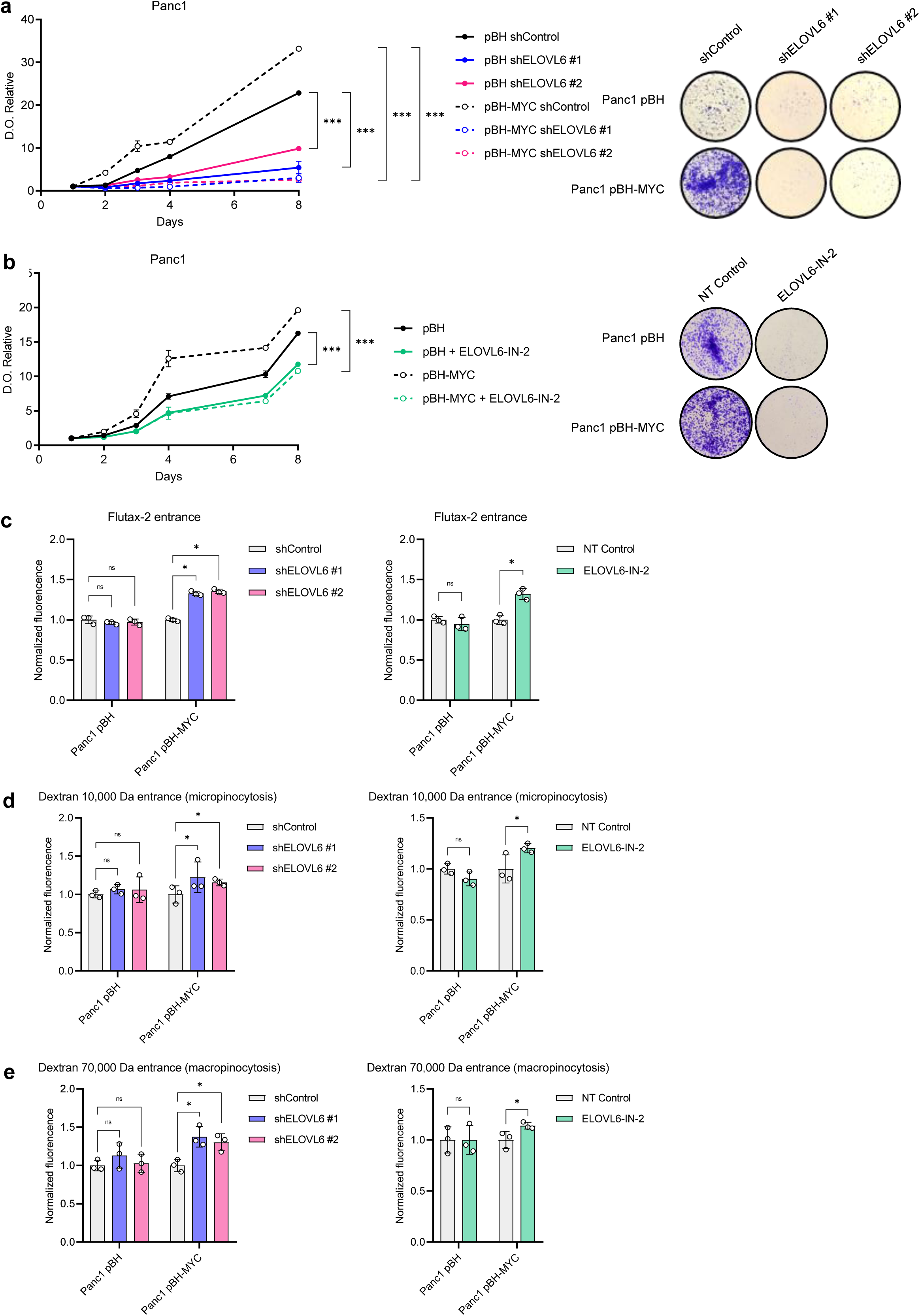
ELOVL6 interference efficacy is dependent on *c-MYC* expression levels. **a** Proliferation and colony assays of *ELOVL6*-silenced cells compared to non-targeted control in Panc1 (pBH and pBH-MYC); n = 3. **b** Proliferation and colony assays of ELOVL6-inhibited cells compared to non-treated control in Panc1 (pBH and pBH-MYC); n = 3. **c** Normalized Flutax-2 uptake in *ELOVL6*-silenced cells compared to non-targeted control in PDAC cell lines and ELOVL6-inhibited cells compared to non-treated control in Panc1 (pBH and pBH-MYC); n = 3. **d** Micropinocytosis analysis by normalized fluorescence of dextran 10,000 Da incorporated by *ELOVL6*-silenced cells compared to non-targeted control and ELOVL6-inhibited cells compared to non-treated control in Panc1 (pBH and pBH-MYC); n = 3 per condition. **e** Macropinocytosis analysis by normalized fluorescence of dextran 70,000 Da incorporated by *ELOVL6*-silenced cells compared to non-targeted control and ELOVL6-inhibited cells compared to non-treated control in Panc1 (pBH and pBH-MYC); n = 3 per condition. All data are presented as mean ± SD; ns: not statistically significant, *p ≤ 0.05, ***p ≤ 0.001; (a, b) Two-way ANOVA test followed by Dunnett test (a) or Sidak test (b); (c-e) Mann-Whitney test. Source data are provided as a Source Data file.

Furthermore, the analysis of total fatty acid composition using isotopically labelled media also confirmed an increase in fatty acid metabolism upon *c-MYC* overexpression **(Supplementary Fig. 14c and 14c)**.

Significantly, the impact of *c-MYC* overexpression was even more evident when assessing cell permeability. In Panc1-pBH-Myc cells, we detected that ELOVL6 interference using shRNAs or ELOVL6-IN-2 caused an increase in permeability, measured by calcein **(Supplementary Fig. 18b and 18c)** and Flutax-2 **(Fig. 9c)** uptake specifically. It also elevated micro and macropinocytosis **(Fig. 9d and 9e)**. Notably, these effects were not observed when interfering ELOVL6 in Panc1-pBH cells.

Collectively, these results underscore that the effects of ELOVL6 interference on cell membrane permeability are intricately tied to the levels of c-MYC and ELOVL6 in the cell, highlighting the tumor’s dependency on the c-MYC/ELOVL6 axis.

### 2.11. ELOVL6 interference reduces tumor growth and synergizes with Abraxane *in vivo*

In our *in vivo* validation, we implanted T3M4 cells subcutaneously in nude mice, distinguishing between those subjected to *ELOVL6* silencing through shRNAs and non-target controls. To confirm the observed *in vitro* synergistic effect, we administered Abraxane at a dose of 40 mg/kg or PBS (vehicle) twice per week at specified intervals, closely monitoring tumor growth. We observed a diminished tumor growth upon *ELOVL6* silencing and this effect was significantly amplified with Abraxane treatment, impairing tumor relapse **(Fig. 10a)**. This underscores the pivotal role of ELOVL6 in the proliferation and growth of human PDAC cells and emphasizes how silencing *ELOVL6* sensitizes tumor cells to Abraxane. Hence, mice implanted with *ELOVL6*-silenced cells exhibited prolonged survival, further accentuated with Abraxane treatment **(Fig. 10b)**. For further validation of the effect of ELOVL6 inhibition through ELOVL6-IN-2 treatment *in vivo* and its synergy with Abraxane, we implanted T3M4 cells in nude mice and administered ELOVL6-IN-2 at a dose of 10 mg/kg or methylcellulose (vehicle) twice per week at specified intervals. In tandem, we administered Abraxane at a dose of 40 mg/kg or PBS (vehicle). These interventions led to reduced tumor growth upon ELOVL6 inhibition, markedly intensified with Abraxane treatment **(Fig. 10c)**. Particularly noteworthy was the significant impact on survival resulting from the combined ELOVL6-IN-2 and Abraxane treatments, that impaired tumor relapse **(Fig. 10d)**.

**Fig. 10.**
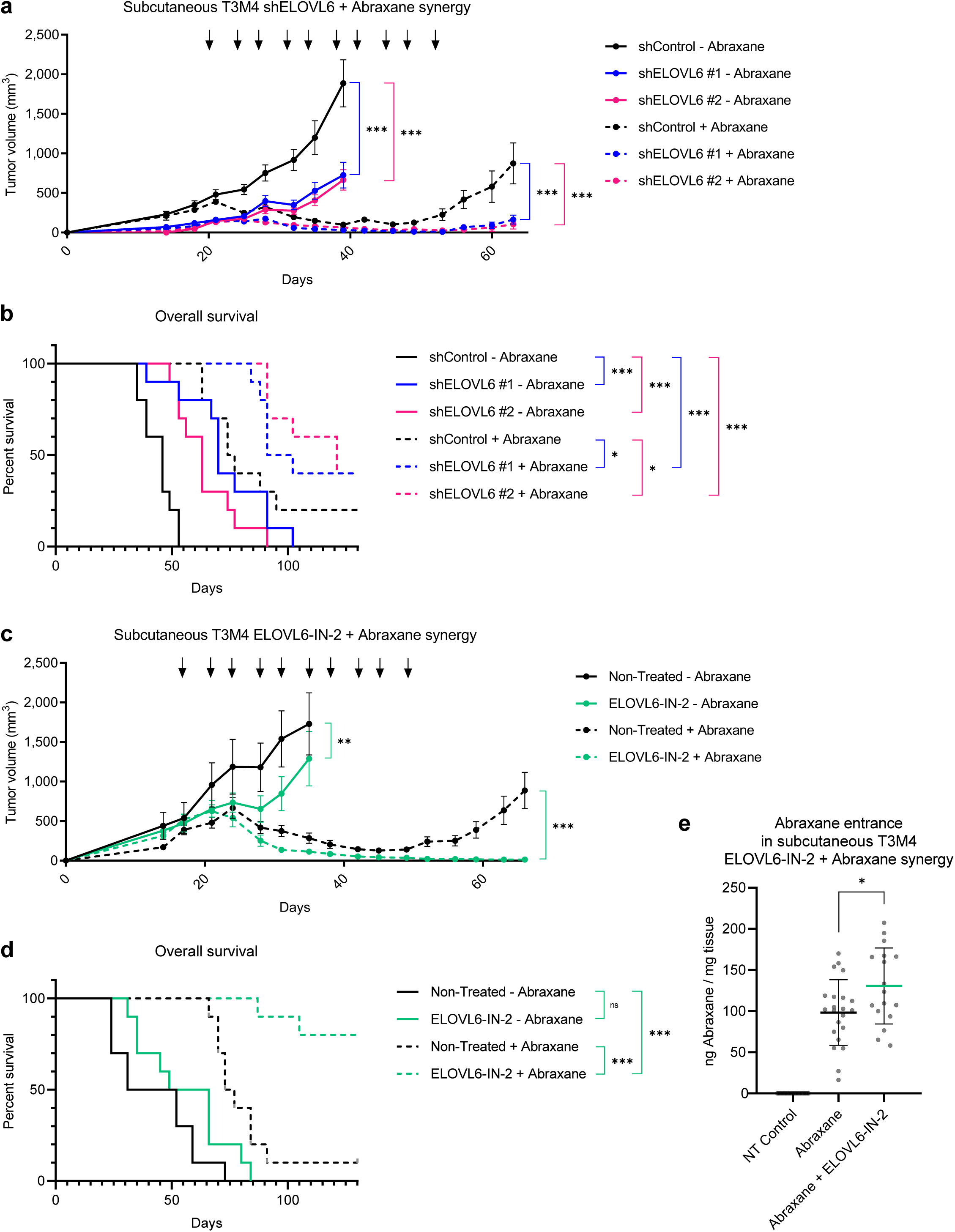
ELOVL6 interference reduces tumor growth and synergizes with Abraxane *in vivo.* **a** Tumor growth in nude mice subcutaneously implanted with T3M4 either *ELOVL6*-silenced or non-targeted control cells; Abraxane or PBS was administered i.p. at a dose of 40 mg/kg twice per week when tumor volume reached 500 mm^3^; n = 10. **b** Survival analysis of the mice in (**10a**). **c** Tumor growth in nude mice subcutaneously implanted with T3M4 either ELOVL6-inhibited or non-treated control cells; Abraxane or PBS was administered i.p. at a dose of 40 mg/kg twice per week when tumor volume reached 500 mm^3^; ELOVL6-IN-2 or methylcellulose was administered by oral gavage at a dose of 10 mg/kg in combination with chemotherapy; n = 10. **d** Survival analysis of the mice in (**10c**). **e** Accumulation of Abraxane inside the tumors determined by mass spectrometry. All data are presented as mean ± SD; ns: not statistically significant, *p ≤ 0.05, **p ≤ 0.01, ***p ≤ 0.001; (a, c) Two-way ANOVA test followed by Dunnett test (a) or Sidak test (c); (b, d) Log-rank (Mantel-Cox) test; (e) Unpaired T test. Source data are provided as a Source Data file.

Furthermore, to determine if this synergistic effect was linked to an increased entry of Abraxane into the tumor due to ELOVL6 inhibition, the intratumoral levels of Abraxane were quantified using a targeted liquid chromatography coupled to tandem mass spectrometry method in mice treated with chemotherapy in the presence or absence of the ELOVL6-IN-2 inhibitor. This experiment revealed an increase in Abraxane uptake by tumor cells upon ELOVL6 inhibition, which correlates with the synergistic effect observed **(Fig. 10e)**.

Finally, we designed an inducible CRISPRi system targeting ELOVL6 and selected the sgRNAs which presented a stronger ELOVL6 downregulation both at RNA and protein levels **(Supplementary Fig. 19a and 19c)**, being #6 and #8 the chosen ones. The implantation of these T3M4 modified cell lines subcutaneously in nude mice and the activation of the system through doxycycline diet successfully slowed down tumor growth **(Supplementary Fig. 19c)** and increased survival rates **(Supplementary Fig. 19d)**, thereby validating our previous findings.

This comprehensive set of experiments conclusively demonstrates, for the first time, that ELOVL6 interference -either by shRNAs or the chemical inhibitor ELOVL6-IN-2-synergizes with Abraxane *in vitro* and *in vivo*, culminating in a robust and promising antitumoral response.

## 3. DISCUSSION

The present study establishes ELOVL6 as a promising therapeutic target in pancreatic ductal adenocarcinoma (PDAC) downstream of c-MYC. Our investigation initially demonstrates a meaningful correlation between *c-MYC* and *ELOVL6* expression. ELOVL6 interference showcases impressive outcomes, impairing tumor growth both *in vitro* and *in vivo*. This intervention induces G1 phase arrest, leading to a notable increase in overall survival in mice. These positive effects are associated with alterations in membrane lipid composition, resulting in changes in rigidity and permeability, along with a reduction in thickness. These changes in mechanical properties increase Abraxane uptake *in vitro*, enhancing its therapeutic effect *in vitro* and *in vivo*.

Tumor cells undergo membrane reorganization to facilitate proliferation, evade apoptosis, and hinder the entry of chemotherapeutic agents^28^. *Frallicciardi et al.* demonstrated how the length of these fatty acids contributes to reduced fluidity and permeability of cell membranes^29^. Other studies have shown that the length of fatty acids is also pivotal in the formation of lipid rafts^30,31^, critical microdomains in the plasma membrane that actively participate in essential processes, particularly signal transduction. Regarding sphingolipids, the length of fatty acids composing them can influence protein dimerization across the membrane. For instance, the modulation of the dimerization of the ErbB2 receptor is involved in cell signaling through an increase in the length of lipid tails^32^. Furthermore, it has been established that elongases play a crucial role in other tumors, influencing EGFR localization, dimerization, and signaling, as demonstrated in the work of *Gimple et al.*^33^.

Changes in lipid profile may alter the biophysical properties of tumor cells’ membranes, particularly rigidness and permeability. This, in turn, influences the entry of numerous chemotherapeutic drugs that require interaction with or penetration through cell membranes to reach their targets^34^. In this context, recent studies have indicated that within populations comprising cells with both high and low permeability membranes, the highly permeable cells exhibit greater reactivity to drug uptake. This phenomenon exerts a negative influence on the uptake by low permeability cells, as the extracellular space quickly becomes depleted of the drug^35^.

The important role of the elongases during tumor progression is supported by several studies. Investigations into the lipidome alterations in lung and prostate cancers have revealed elongation of acyl chains in phospholipids as a common oncogenic trait. Notably, in lung squamous cell carcinoma (SCC), analysis of 30 human SCCs and the L-Ikkα^KA/KA^ mouse model identified ELOVL6 as the primary enzyme driving this lipid modification, correlating with enhanced tumor growth and colony formation^36^. Similarly, androgen-regulated changes in prostate cancer highlight the upregulation of ELOVL5, which is pivotal for mitochondrial function and metastatic progression^37^. These findings underscore the potential of targeting ELOVL enzymes, particularly ELOVL6 in lung SCC and ELOVL5 in prostate cancer, to disrupt cancer cell lipid metabolism and offer new avenues for therapeutic intervention.

Inhibition of lipogenesis has demonstrated noteworthy benefits in cancer treatment, with certain inhibitors progressing to preclinical stages. Nevertheless, their potential in pancreatic ductal adenocarcinoma (PDAC) remains largely unexplored, necessitating further investigation^38^. Notably, FASN inhibitors have undergone testing in PDAC, revealing a synergistic effect with gemcitabine in both cell and mouse models, leading to apoptosis attributed to endoplasmic reticulum stress^39–41^. Additionally, these FASN inhibitors have demonstrated synergy with paclitaxel^42^, aligning with our *in vitro* and *in vivo* results using the ELOVL6 inhibitor. However, inhibitors targeting ACCA or ACLY, although effective, await testing in PDAC^43^.

Certainly, an evolving field is emerging, centered around the investigation of tumor cell membrane composition and its modification for the development of novel drugs to enhance their efficacy^44–46^. Targeting ELOVL6 to selectively alter the cell membranes of tumor cells could potentially enhance chemotherapeutic drug delivery and could serve as a valuable strategy to combat drug resistance and overcome alterations in membrane composition. This synergy with chemotherapeutic agents offers a promising avenue to enhance efficacy while reducing dosages and mitigating off-target effects. Importantly, targeting ELOVL6 may provide therapeutic benefits to patients characterized by elevated c-MYC levels linked to KRAS activation, as well as copy number gains and amplifications. However, it is important to approach this strategy with caution, as inhibitors could also affect ELOVL6 activity in other organs, such as the liver where *ELOVL6* is highly expressed.

In summary, the current therapeutic options for treating pancreatic ductal adenocarcinoma (PDAC) are severely limited, underscoring the imminent need to identify new targets. Lipid synthesis stands as a crucial aspect of tumor progression, presenting an unexplored field in cancer treatment. Our findings position ELOVL6 as a novel therapeutic target, showcasing the effectiveness of its chemical inhibition in PDAC. This synergy with Abraxane treatment not only highlights the potential for enhanced efficacy but also unveils a spectrum of possibilities for advancing PDAC treatment strategies.

## 4. MATERIALS AND METHODS

### 4.1. Reference datasets

#### Sanchez-Arevalo et al. dataset

Pancreas from *Ela1-Myc* and Control mice were used to assess the Myc-mediated transcriptomic profile through RNA-seq and its binding sites through ChIP-seq. We downloaded FASTQ files through GEO Series accession number GSE77411 and GSE77410.

#### Walz et al. dataset

Pancreatic tumors isolated from KPC (*Pdx1-cre; LSL-Kras^G12D^; LSL-Trp53^R172H^)* pancreas mice were subjected to ChIP-seq against c-Myc. We downloaded the FASTQ files through GEO Series accession number GSE44672.

#### Sabo et al. dataset

3T9 mouse fibroblast cells were infected with MYCER, cells were treated with OHT, and samples collected at different timepoints 2, 4, 8 and 16h for RNA-seq. B-cells obtained from the *Eµ-Myc* mouse model were isolated at different stages (Control, Pre-tumoral and Tumoral stages) for RNA-seq and ChIP-seq. We downloaded the count expression matrix and the binding enrichment through GEO Series accession number GSE51011.

#### Schlesinger et al. dataset

To explore cell heterogeneity following the expression of constitutively active *Kras* in acinar cells, which causes acinar metaplasia and pancreatic dysplasia, we carried out analyses of scRNA-seq experiments of pancreatic tissues. Tamoxifen was injected into six- to eight-week-old *Ptf1a-CreER, LSL-Kras-G12D, LSL-tdTomato (PRT)* mice, and the pancreas was collected for single-cell isolation at six different timepoints post-tamoxifen injection (PTI). We downloaded the dataset through GEO Series accession number GSE141017.

#### Peng et al. dataset

Single-cell RNA-seq profiles from 24 PDAC tumor samples and 11 control pancreases without any treatment. Fresh specimens of control pancreases were harvested from 3 cases of non-pancreatic tumor patients (e.g. bile duct tumors or duodenal tumors) and 8 cases of non-malignant pancreatic tumor patients (e.g. pancreatic cyst) receiving pancreatoduodenectomy or distal pancreatectomy. We downloaded the dataset through GSA accession number CRA001160.

### 4.2. RNA-seq

Data quality and Trimming: it was controlled using FastQC (v.0.11.9). After trimming the adaptor and removing low-quality reads with Trim Galore (v.0.6.10) we proceeded to the mapping. Mapping: alignment of trimmed fastq files was performed with STAR aligner (v.2.7.10b)^47^ using hg38 as the reference genome. Gene-level expression data in terms of expected counts were obtained using FeatureCounts from the Subread R package (v.2.01)^48^. Differential expression: differential analyses of the expression profiles between different conditions were performed using the DESeq2 package (v.1.38.0) based on negative binomial distribution^49^. P-values were FDR-adjusted with Benjamini-Hochberg method.

### 4.3. RNA-Seq GSEA analysis

We carried out a functional analysis to determine the altered cell processes in treated vs untreated T3M4 cells using GSEA^50^. The molecular signatures files tested were HALLMARK_G2M_CHECKPOINT.gmt, HALLMARK_MYC_TARGETS_V2.gmt, GOBP_FATTY_ACID_METABOLIC_PROCESS.gmt, GOBP_MEMBRANE_LIPID_METABOLIC_PROCESS.gmt, HALLMARK_MYC_TARGETS_V1.gmt, GOBP_CELL_CYLCE_PROCESS.gmt, GOBP_FATTY_ACYL_COA_METABOLIC_PROCESS.gmt, GOBP_LONG_CHAIN_FATTY_ACYL_COA_METABOLIC_PROCESS.gmt, GOBP_FATTY_ACID_DERIVATIVE_METABOLIC_PROCESS.gmt, GOBP_ERBB2_EGFR_SIGNALING_PATHWAY.gmt and KOBAYASHI_EGFR_SIGNALING_6HR_DN. Significant gene sets enriched by treatment of control cells were identified using a false discovery rate FDR-q-value < 0.25 (Benjamini and Hochberg correction) and a nominal P value < 0.05. All analyses were performed using GSEA v2.1 software (www.broadinstitute.org/gsea) with 1000 data permutations.

### 4.4. ChIP-seq

Data quality and Trimming: it was controlled using FastQC (v.0.11.9). After trimming the adaptor and removing low-quality reads we proceeded to the mapping. Mapping: the obtained clean data was aligned to human genome hg38 by the Burrows–Wheeler Aligner tool BWA (v.0.7.17)^51^. ChIP quality controls: read counts were analyzed by Deeptools (v.3.13.3)^52^; as quality controls of the experiments we used the functions plotFingerprint and plotCoverage. The function computeMatrix was used to generate the plotHeatmap and plotProfile. Bigwig files were generated for peaks visualization using the function bamCompare. Track visualization: integrative Genomics Viewer (IGV) (v.2.4.9) was used to track visualization. Peak calling: a peak calling analysis was conducted using MACS2 (v.2.2.71)^53^ with the following parameters: macs2callpeak -t Bam_file_sort_nondup.bam -c Bamfile_file_control_sort_nondup.bam -f BAM -g 1.87e9 -n File_name -B -q 0.05. Motif analysis: the TSSs were annotated to genomic feature using the annotatePeaks.pl of HOMER (v.4.11) software^54^. Transcription factor binding motifs were identified with HOMER findMotif-sGenome.pl tool in the chromatin-accessible region; those with P<0.05 were considered significant.

### 4.5. Functional enrichment analysis

Over Representation Analysis (ORA) was performed using the GSEApy python package^55^, which implements GSEA and over-representation analysis (ORA). We validated the results for gene ontology (GO) term enrichment and the Kyoto Encyclopedia of Genes and Genomes (KEGG). Enriched terms with FDR-adjusted P<0.05 were deemed as statistically significant.

### 4.6. scRNA-seq

Data preprocessing: filtered raw counts data were imported into Scanpy (v.1.7.2) for further processing and analysis^56^. Raw transcript counts of gene-cell matrices were filtered to remove cells with fewer than 200 transcripts and cells with higher than 20% mitochondrial contents were excluded from the analysis. Features expressed in less than three cells were also excluded. We excluded the doublets in the datasets using Solo from scVI tools (v.0.19.0)^57,58^. Dimension reduction and clustering analysis: we used Scanpy for data analysis and visualization with the following steps performed in order: data normalization, log-transformation, highly variable genes (HVG) selection and principal component analysis (PCA). Genes with the highest variance were used to perform linear dimensional reduction (principal component analysis), and the number of principal components used in downstream analyses was chosen considering Elbowplot. Expression levels were normalized to counts per 10,000 (NC). The normalized counts were then transformed by log2 (NC + 1) and used for further analysis. Highly variable genes were selected based on the log-transformed data. The principal component analysis was run with the selected high variable genes. If needed, the HVG and PCA were re-calculated for subsets of cells. Uniform Manifold Approximation and Projection (UMAP) and Leiden clustering were performed using Scanpy. Firstly, the neighborhoods were calculated with ‘scanpy.pp.neighbors’. The connectivities were computed using the ‘umap’ method with Euclidean distance. Then UMAP embedding was calculated using ‘scanpy.tl.umap’. The Leiden clustering was calculated using ‘sc.tl.leiden’ on the neighborhood calculated in the previous steps, with the resolution of 1 for *Schlesinger et al.* dataset^26^ and 0.7 for *Peng et al.* dataset^27^. Scanpy and seaborn were used for data visualization. Integration: to integrate data from different samples we used scVI tools (v.0.19.0). Differentially expressed genes (DEGs): Wilconxon test implemented in the function ‘sc.ranked_genes_group()’ was used to obtain DEGs between cell types. Genes with p-adj < 0.05 and |Log2foldchange| > 1 were selected as DEGs.

### 4.7. Cell culture

Hek293T, and human PDAC cells -Patu 8988T, Panc1 and T3M4-were cultured in Dulbecco’s modified Eagle’s medium (DMEM; LONZA, Ref 12-604F) supplemented with 10% fetal bovine serum (FBS; TICO EU, Ref A3FBSEU500)) and 1% penicillin/streptomycin (Life Technologies). For ELOVL6-inhibited cells, this medium was supplemented with the chemical inhibitor ELOVL6-IN-2 10 nM (MedChemExpress, Ref HY-12146) for 48 hours. For the inducible CRISPRi system experiments, cells were supplemented with doxycycline 1 µg/mL for 72 hours. All cell lines were kindly provided by Dr. Francisco X Real (Centro Nacional de Investigaciones Oncologicas, CNIO).

### 4.8. Retroviral and lentiviral vector constructs, and virus production

c-MYC downregulation was performed using the plasmid pRS-shMYC and compared to an empty vector (pRS, shControl). c-MYC overexpression was performed using the plasmid pBH2-c-MYC and compared to an empty vector (pBH2). pBH2, pBH2-c-MYC and shRNAs were kindly provided by Dr. Bruno Amati (European Institute of Oncology, EIO, Milan, Italy). For retroviral production, infectious retroviruses were produced in Hek293T cells by calcium chloride transfection of the retroviral construct together with the packaging plasmid pCL-Ampho.

Mission shRNAs (Sigma-Aldrich) were used for RNA-interference. Two ELOVL6-targeting shRNAs (shELOVL6 #1 -clone TRCN0000163912- and shELOVL6 #2 -clone TRCN0000161318-) were used and compared to a control non-targeting shRNA (pLKO-shControl). shRNAs were kindly provided by Dr. Francisco X Real (CNIO). For lentiviral production, infectious lentiviruses were produced in Hek293T cells by calcium chloride transfection of the lentiviral construct together with the packaging plasmids psPAX2 and pCMV-VSV-G.

Mediums were harvested twice (48 h and 72 h post-transfection). Viral supernatants were filtered and applied on target cells in the presence of 5 µg/ml polybrene. Cells were used after 72 h puromycin selection or one-week hygromycin selection.

To generate a knockout (KO) of the *ELOVL6* gene, two CRISPR-Cas9 guide RNAs (sgRNAs) were designed to target the first and second exons of the gene. The sgRNAs were designed using Benchling CRISPR design tool (www.benchling.com) following established protocols^59^. The sequences of the sgRNAs were sgELOVL6 #1: ACGAGAATGAAGCCATCCAA and sgELOVL6 #2: AACCTACCTGAAGACTGCAA. Custom DNA oligos for these sgRNAs were synthesized by SIGMA. The sgRNAs were annealed and subsequently ligated into the BsmBI-digested lentiCRISPRv2 plasmid (Addgene, #52961). The integrity and accuracy of the cloned sgRNA sequences were confirmed by Sanger sequencing. LentiCRISPRv2 viral vectors containing either the ELOVL6 sgRNAs or a control sgRNA (sgAAVS1: CCTCTAAGGTTTGCTTACGA) were used. Cellular clones with complete KO of *ELOVL6* expression were selected for subsequent experiments.

For *ELOVL6* knock-down (KD) experiments, a CRISPR inhibition (CRISPRi) strategy was employed, using a lentiviral and inducible vector, pLVXTet on dCas9-KRAB, previously developed in the laboratory. The T3M4 cell line was first transduced with recombinant lentiviral vectors generated from this plasmid at a multiplicity of infection (MOI) of 2. Cell clones with no detectable leaky expression were selected for further use in this study. sgRNAs were designed to target sequences at the end of the promoter region or within the initial bases of the transcriptional start site, following established guidelines^60^. In line with these criteria, five distinct sgRNAs were generated: sgELOVL6 #3: GACGACCGCTGGAGACCGAG, sgELOVL6 #4: CAGCCCTGGATGTAGCTGAG, sgELOVL6 #5: GCGTCCGCATCCACCGTAGG, sgELOVL6 #6: AGGGTGATGGACAAACGTGC and shELOVL6 #8: CAGCCTCTCAGCTACATCCA. These sgRNAs were cloned into the lentiGuide-hygro-eGFP vector (Addgene, #99375) using the methodology described above, and tested to be selected.

### 4.9. Quantitative real-time PCR

Total RNA was isolated from cultured cells using the NucleoSpin RNA kit for RNA purification (MACHEREY-NAGEL, Ref 22740955.250) according to manufacturer’s instructions. Samples were treated with DNase I before reverse transcription. cDNA was generated from 1 μg RNA using random hexamers and reverse transcriptase (TaqMan Reverse Transcription Reagents; Thermo Fisher Scientific™, Ref N8080234). qPCR amplification and analysis were conducted using the 7500HT Real-Time PCR System (ABiosystems) using GoTaq^®^ qPCR Master Mix (Promega, Ref A6002). RNA levels were normalized to *HPRT* expression using the DDCt method. Primer sequences are provided in **Supplementary Table 1**.

### 4.10. Western blot

Total protein was isolated from cultured cells by cell lysis and sonication. Equal amounts of protein were loaded in polyacrylamide gels and SDS-PAGEs were performed. Proteins were transferred to PVDF membranes and blocked with 5% powdered skimmed milk in TTBS. Membranes were incubated with the antibodies provided in **Supplementary Table 2**. Membranes were imaged using Clarity Western ECL Substrate (BIO-RAD, Ref. 1705061) and a ImageQuant LAS 4000 CCD camera.

### 4.11. Chromatin immunoprecipitation (ChIP)

10^8^ cells (T3M4) were fixed with 1% formaldehyde for 10 min at room temperature. Fixation was stopped by adding glycine (to 125 mM) with an additional incubation of 5 min. Cells were collected by scraping, pelleted and then lysed in ice for 10 min in 5 ml of buffer with 50 mM HEPES (pH 7.5), 140 mM NaCl, 1 mM EDTA, 0.5% NP-40, 0.25% Triton X-100 and 10% glycerol. After centrifugation, pelleted nuclei were resuspended in 5 ml of buffer with 10 mM Tris-HCl (pH 8.0), 200 mM NaCl, 0.5 mM EGTA and 1 mM EDTA, and incubated at room temperature for 10 min. Pelleted nuclei were resuspended in 1 ml of buffer with 100 mM NaCl, 50 mM Tris-HCl (pH 8), 5 mM EDTA (pH 8), 0.5% SDS and protease inhibitors (Qiagen, Valencia, CA, USA). We sonicated for 3 min in a Covaris sonicator, yelding DNA fragments of 200–500 bp. 2 mg protein were precleared and incubated overnight with c-MYC antibody (Cell Signaling, Ref 13987). Samples were incubated with beads (Cell Signaling, Ref 9007) for 2 hours at 4°C and then washed with Triton buffer (100 mM Tris-HCl (pH 8), 100 mM NaCl, 5 mM EDTA (pH 8) and 5% Triton X-100), mixed micelle wash buffer (150 mM NaCl, 20 mM Tris-HCl (pH 8), 5 mM EDTA (pH 8), 5% sucrose, 1% Triton X-100 and 0.2% SDS), Buffer 500 (0.1% sodium deoxycholate, 1 mM EDTA (pH 8), 50 mM HEPES (pH 7.5), 1% Triton X-100 and 500 mM NaCl), LiCl buffer (0.5% sodium deoxycholate, 1 mM EDTA (pH 8), 250 mM LiCl, 10 mM Tris-HCl (pH 8) and 0.5% NP-40) and Tris-EDTA (TE). DNA was eluted in buffer with 1% SDS and 1% sodium bicarbonate. RNA and protein were digested using proteinase K and crosslinks were reversed by incubation overnight at 65°C. DNA was purified and target DNA abundance was assayed by RT-qPCR with primer pairs designed to achieve products of 50–200 bp (**Supplementary Table 3**).

### 4.12. Growth and colony assays

To determine proliferation, 2.5×10^3^ cells per well were seeded in 96-well plates. After 1, 2, 3, 4, 7 (in inhibitor-treated cells) and 8 days, cells were washed with PBS 1X, fixed with 0.5% glutaraldehyde and incubated with 0.5% crystal violet in 25% methanol; crystal violet was eluted with 10% acetic acid and the OD590 nm was determined. To assess colony formation, 8×10^2^ cells were seeded in 6-well plates and after 14 days, cells were processed as described earlier, fixing with ice-cold methanol instead.

### 4.13. Cell cycle assay

For cell cycle profiling, cells were fixed with ice-cold 70% ethanol for 1 h, treated with 0.25% Triton X-100 for 15 minutes, and stained at room temperature for 30 minutes with 20 µg/ml propidium iodide (Life Technologies, Ref P3566) containing 10 µg/ml RNAse A. Cells were processed by flow cytometry (FACSCalibur Becton Dickinson) and the results were analyzed by FlowJo v10.

### 4.14. Apoptosis assay

Cells were resuspended in 200 μl of Binding Buffer (10 mM HEPES/NaOH pH = 7.4; 140 mM NaCl; 2.5 mM CaCl_2_) with 0.5 μg/ml of Annexin V - FITC Conjugate (Biotium, Ref 29001) and incubated 15 minutes at RT. 2.5 μg/ml of propidium iodide (Life Technologies, Ref P3566) was added and after 30 minutes cells were processed by flow cytometry (FACSCalibur Becton Dickinson). The results were analyzed by FlowJo v10.

### 4.15. Migration assays

For transwell assays 2×10^5^ cells were seeded in each upper chamber in 0.1% FBS DMEM and placed in a well with 10% FBS DMEM. Next day, the chambers were fixed with 0.5% glutaraldehyde, washed twice with PBS 1X, and incubated with 0.5% crystal violet in 25% methanol. Each chamber was photographed using optical microscopy and analyzed using ImageJ/Fiji. For wound healing assays, cells were growth until reaching confluence in 10% FBS DMEM. A wound was performed using a tip to scratch the plate longitudinally, the media was replaced by 0.1% FBS DMEM and wound healing was monitored during 4, 6 and 24 h. Wound healing was quantified using ImageJ/Fiji software.

### 4.16. Cell signaling analysis

Cell signaling was determined using an array for the parallel determination of the relative phosphorylation levels of human protein kinases, following the manufacturer’s instructions (Biotechne, Ref ARY003C). The array membranes were developed using an ImageQuant LAS 4000 CCD camera.

### 4.17. Lipidic profile analysis

#### Lipid extraction

An amount of cells containing 10 μg of DNA was homogenized in 700 μL of water with a handheld sonicator and was mixed with 800 μl HCl(1M):CH_3_OH 1:8 (v/v), 900 μl CHCl_3_, 200 μg/ml of the antioxidant 2,6-di-tert-butyl-4-methylphenol (BHT; Sigma-Aldrich, Ref 8.22021) and 3 μl of UltimateSPLASH™ ONE internal standard mix (Avanti Polar Lipids, Ref 330820). After vortex and centrifugation, the lower organic fraction was collected and evaporated using a Savant SpeedVac SPD111V (Thermo Fisher Scientific™) at room temperature, and the remaining lipid pellet was stored at −20°C under argon.

#### Mass spectrometry

Just before mass spectrometry analysis, lipid pellets were reconstituted in 100% ethanol. Lipid species were analyzed by liquid chromatography electrospray ionization tandem mass spectrometry (LC-ESI/MS/MS) on a Nexera X2 UHPLC system (Shimadzu) coupled with hybrid triple quadrupole/linear ion trap mass spectrometer (6500+ QTRAP system; AB SCIEX). Chromatographic separation was performed on a XBridge amide column (150 mm × 4.6 mm, 3.5 μm; Waters) maintained at 35°C using mobile phase A [1 mM ammonium acetate in water-acetonitrile 5:95 (v/v)] and mobile phase B [1 mM ammonium acetate in water-acetonitrile 50:50 (v/v)] in the following gradient: (0-6 min: 0% B → 6% B; 6-10 min: 6% B → 25% B; 10-11 min: 25% B → 98% B; 11-13 min: 98% B → 100% B; 13-19 min: 100% B; 19-24 min: 0% B) at a flow rate of 0.7 mL/min which was increased to 1.5 mL/min from 13 minutes onwards. SM, CE, CER, DCER, HCER, LCER were measured in positive ion mode with a product ion of 184.1, 369.4, 264.4, 266.4, 264.4 and 264.4 respectively. TAG, DAG and MAG were measured in positive ion mode with a neutral loss for one of the fatty acyl moieties. PC, LPC, PE, LPE, PG, PI and PS were measured in negative ion mode by fatty acyl fragment ions. Lipid quantification was performed by scheduled multiple reactions monitoring (MRM), the transitions being based on the neutral losses or the typical product ions as described above. The instrument parameters were as follows: Curtain Gas = 35 psi; Collision Gas = 8 a.u. (medium); IonSpray Voltage = 5500 V and −4,500 V; Temperature = 550°C; Ion Source Gas 1 = 50 psi; Ion Source Gas 2 = 60 psi; Declustering Potential = 60 V and −80 V; Entrance Potential = 10 V and −10 V; Collision Cell Exit Potential = 15 V and −15 V.

The following fatty acyl moieties were taken into account for the lipidomic analysis: 14:0, 14:1, 16:0, 16:1, 16:2, 18:0, 18:1, 18:2, 18:3, 20:0, 20:1, 20:2, 20:3, 20:4, 20:5, 22:0, 22:1, 22:2, 22:4, 22:5 and 22:6 except for TGs which considered: 16:0, 16:1, 18:0, 18:1, 18:2, 18:3, 20:3, 20:4, 20:5, 22:2, 22:3, 22:4, 22:5, 22:6. P- vs O-ether lipid distinction relies on the biological assumption that the ether bonded acyl chains are predominantly saturated and that unsaturations are only present in the ester bonded acyl chains.

#### Data Analysis

Peak integration was performed with the MultiQuant™ software version 3.0.3. Lipid species signals were corrected for isotopic contributions (calculated with Python Molmass 2019.1.1) and were quantified based on internal standard signals and adheres to the guidelines of the Lipidomics Standards Initiative (LSI) (level 2 type quantification as defined by the LSI). Unpaired T test p-values and FDR corrected p-values (using the Benjamini/Hochberg procedure) were calculated in Python StatsModels version 0.10.1.

*Only relevant if you use the univariate statistics from the html report.

### 4.18. Analysis of FA metabolism

Isotopically-labelled media were prepared from DMEM without glucose, glutamine, sodium pyruvate (D9802-01, US-Biological). U-^13^C-glucose (CLM-1396, Cambridge Isotope Laboratories) and the rest of components were added at the normal concentration found in DMEM media. The media was supplemented with 10% fetal bovine serum and 1% penicillin/streptomycin.

Human PDAC cells T3M4 were seeded at 5×10^5^ cells/well in 6-well plates in DMEM. After 24h, media was changed to DMEM containing U-^13^C-glucose. Cells were incubated with U-^13^C-glucose for 96 h, media was renewed every 24 h. At the end of the incubation, media were removed, cells were washed once with cold PBS 1X, scraped with 500 μl of cold PBS 1X, transferred to 1.5 ml Eppendorf tubes and stored at −80°C^61^.

Fatty acid (FA) saponification and LC-MS-based analysis was performed as previously described^61^. To analyze the total FAs, 450 μl of cell suspension were transferred to a glass vial, and 1,000 μl of a 9:1 MeOH:KOH (3M in H_2_O) solution containing PC(16:0/16:0)D62 at 3 ppm were added. Saponification was performed for 1 h at 80 °C in a water bath. After saponification, samples were cooled on ice and acidified by adding 100 μl of formic acid. FAs were extracted with 2 ml of heptane:isooctane (1:1) (2x), dried in a nitrogen flow, resuspended in 200 μl of mobile phase A containing myristic acid D27 at 1 ppm and transferred to a glass HPLC vial. FAs were analyzed in a quadrupole– orbitrap mass spectrometer (Q Exactive, Thermo-Fisher Scientific) coupled to reverse phase chromatography via electrospray ionization. Liquid chromatography separation was performed in a Cortecs C18 column (2.1 mm × 150 mm, 1.6 μm particle size; Waters). Solvent A was 2.5 mM ammonium acetate in 60:40 water:methanol. Solvent B was 2.5 mM ammonium acetate in 95:5 acetonitrile:isopropanol. The flow rate was 300 μl/min, the column temperature was 45°C, the autosampler temperature was 5 °C and the injection volume was 5 μl. The liquid chromatography gradient was: 0 min, 45% B; 0.5 min, 45% B; 19 min, 55% B; 23 min, 99% B; 34 min, 99% B. Between injections, the column was washed for 2 min with 50:50 acetonitrile:isopropanol before being equilibrated to the initial conditions. The mass spectrometer operated in the negative-ion mode to scan from m/z 100 to 450 at a resolving power of 140000. Data were acquired in the centroid mode.

Data preprocessing was performed using LipidMS3.0^62^. The analysis of FA metabolism was performed using FAMetA^61^.

### 4.19. Abraxane tumoral accumulation measurement

Quantification of Abraxane in tumor samples was performed by using a modified version of a previously described protocol^63^. Frozen tumor samples (5–100 mg) were placed in 2 ml tubes containing CK14 ceramic beads (Precellys Lysing kit, soft tissue homogenizing CK14, Bertin Technologies, France). For each 100 mg of tissue, 1 mL PBS was added. Then, tumors were homogenized at 4°C in a Precellys Evolution system (2 x 30 s at 6,000 rpm, 30 s rest) equipped with a Criolys cooler (Bertin Technologies, France).

Tumor homogenates were transferred to 1.5mL Eppendorf tubes and extracted with 4 volumes of acetonitrile (20 μL sample + 80 μL acetonitrile). Samples were centrifuged twice at 13000 g for 15 min at 4°C, and the clean supernatants were transferred to 96-well plates for their LC-MS/MS-based analysis.

A standard calibration curve of Abraxane was performed in the 20.000 ng/mL to 39 ng/mL range by serial half dilutions in PBS. Then, paclitaxel was extracted using 4 volumes of acetonitrile (40 μL standard + 160 μL acetonitrile). Samples were centrifuged twice at 13000g g for 15 min at 4°C, and the clean supernatants were transferred to 96-well plates for their LC-MS/MS-based analysis.

UPLC separation was performed in an Acquity UPLC system (Waters, UK) equipped with an Acquity UPLC BEH C18 (1.7 μm, 2.1 × 100 mm; Waters) column. The temperatures of the column and the autosampler were set at 40°C and 4°C, respectively. The sample injection volume was 4 μL. Eluents consisted in 10 mM ammonium formate (pH 4.0) (eluent A) and acetonitrile (eluent B). The flow rate was set at 0.4 mL/min. A 5-min elution gradient was performed as follows: initial eluent composition was set at 75% A and 25% B, which was linearly changed to 25% A and 75% B in 2.5 min; then the proportion of B was increased to 100% in the next 1 min. Finally, the initial conditions were recovered and maintained for 1 min for column conditioning.

The MS analysis was performed using a Waters Xevo TQ-S mass spectrometer (Waters) equipped with an ESI source working in positive-ion mode multiple reaction monitoring (MRM) mode. A capillary voltage of 3 kV, a source temperature of 120°C and a desolvation temperature of 500 °C were used. Desolvation and cone gas flows were set as 800 L/h and 150 L/h, respectively, and the collision gas was 0.25 mL/min. The cone voltage was set at 30V and the following transitions were detected: 854>569.3 (collision energy 10eV, employed for quantification), 854>509.3 (collision energy 20eV, employed for confirmation). The data station operating software used was MassLynx 4.1 (Waters).

### 4.20. Calcein-AM assay

5×10^3^ cells were seeded in black 96-well plates. After 24 h, Calcein-AM was added to a final concentration of 5 µM and its uptake was determined using Invitrogen™ Calcein-AM assay kit according to the manufacturer instructions (Life Technologies, Ref C1430). Fluorescence was determined at 494/520 nm using a luminometer reader (EnSpire^®^ Multimode Plate Reader). We also treated trypsinized cells with Calcein-AM and used a THUNDER DMi8 Leica fluorescence microscope to take images. Green channel signal was quantified using ImageJ/Fiji software.

### 4.21. Drug synergy assay

IC50 for albumin-bound paclitaxel nanoparticles (Abraxane^®^; Bristol Myers Squibb, Ref 660458) was determined in each cell line by drug/response curves. 5×10^3^ cells were seeded into 96-well plates and treated with different dilutions of Abraxane^®^ the next day. 72 h after treatment, cell viability was determined using ATPlite 1step Luminescence Assay System (PekinElmer, Ref 6016731) in a luminometer reader (EnSpire^®^ Multimode Plate Reader).

### 4.22. Flutax-2 assay

Flutax-2 (Tocris Bioscience, Ref. 6254) entrance was measured according to manufacturer’s instructions. The dose used was 2 μM in HBSS for 5 minutes. Fluorescence was determined at 496/526 nm in a luminometer reader (EnSpire^®^ Multimode Plate Reader).

### 4.23. Pinocytosis assays

Dextran-Rhodamine B entrance was measured according to manufacturer’s instructions. Dextran-Rhodamine B 10,000 Da (Invitrogen™, Ref. D1824) was used to quantify micropinocytosis and Dextran-Rhodamine B 70,000 Da (Invitrogen™, Ref. D1841) was used to determine macropinocytosis. The dose used for each molecule was 0.2 mg/ml in DMEM without FBS. After 30 minutes, fluorescence was determined at 570/590 nm in a luminometer reader (EnSpire^®^ Multimode Plate Reader).

### 4.24. Immunofluorescence

We followed the protocol established by Le at al.^64^. The day before, we seeded 5×10^4^ cells on coverslips. On the day of the experiment, we put the cells on ice to stop endocytosis and synchronize the cells to enhance dextran entrance. After 30 minutes of treatment with the correspondent dextran (Invitrogen™, Ref. D1824 and D1841) at 0.2 mg/ml in DMEM without FBS, we washed the coverslips three times with cold PBS to remove dextran and stop endocytosis. We fixed the cells 15 minutes with 4% PFA and stained with DAPI 1:5000 for 15 minutes. We mounted the coverslips in slides and used a THUNDER DMi8 Leica fluorescence microscope to take the images. They were processed and red channel signal was quantified using ImageJ/Fiji software.

### 4.25. Electron microscopy

Cells were embedded in TAE 1% agarose and thin slices of the cell blocks were fixed in 2.5% glutaraldehyde, washed in 0.1 M cacodylate buffer, and then fixed in 1% aqueous osmium tetroxide. They were then dehydrated and embedded in epoxy resin. Semithin sections were performed at 1 μm thickness and stained with toluidine blue; they were evaluated using light microscopy to select the most cellular areas. Thin sections were cut from these areas, placed in copper grids and stained with uranyl acetate. They were evaluated using a transmission electron microscope (JEM 1011; JEOL Ltd, Japan).

### 4.26. Membrane tension measurement experiments

We charged trypsinized cells with Calcein-AM (5 µM) according to the manufacturer instructions (Life Technologies, Ref C1430) and used a fluorescence super-resolution microscope to take images for cytometric fields (THUNDER DMi8 Leica with a 63x/ 1.40 Oil objective; HC PL APO. Emission wavelength 525nm). Images were acquired with a Leica KS-14401089 camera (16 bits, 2048×2048) with a resolution of 0.24 µm in the xy-plane. A total of 150 cytometric fields per sample were recorded with an average amount of 20 cells per field imaged. The effective membrane tension was estimated from the static structure factor of cell contours at the equatorial plane, interpreted in terms of the membrane Helfrich Hamiltonian^65–67^. Image processing, cell contour detection, and data analysis were carried out using Mathematica 13.3 software (Wolfram Research, Inc., Mathematica, Version 13.3, Champaign, IL (2023)). In brief, we performed noise reduction through a Gaussian filtering of radius 2-px, being subsequently binarized using a global threshold determined by Otsu’s clustering-based variance maximization method. The morphological components were selectively filtered based on thresholding for both cell radius (greater than 7.5 μm according to the mean size), and a circularity (c > 0.9). We restricted image analysis to suspended cells displaying minimal morphological alterations. Further contour refinement was performed to mitigate the effects of pixelation on the description of normal mode fluctuations. Hence, the raw contours were transformed into polar coordinates, being interpolated by using a 12th-degree BSpline function, and eventually resampled from 0 to 2π at 720 fixed intervals. Finally, each single contour was transformed to complex Fourier space via FFT. For physical analysis, we assumed that the cell culture is biologically homogeneous and represents an ergodic process in such a way that the time average of the cellular membrane fluctuations (static factor) is equivalent to the statistical average of the cell ensemble which shows morphological variability at any given time. This allows for an efficient estimation of the mean membrane rigidness of the cell ensemble. Due to the largest apparent size of the detected components we focused on the Helfrich modes of the longest wavelength, thereby the membrane tension dominates, dealing with an estimator for cell rigidness as:

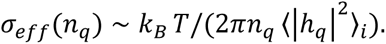

### 4.27. Indentation experiments by optical tweezers (OTs)

Pristine cells resuspended in DMEM buffer were incubated subconfluent for 1.5 hours in a microscope glass coverslip (0.17mm thickness) (aliquot of 45 μL from culture 10^5^ cells/ml). Polystyrene beads with a mean particle size of 5.7 μm were added (Sigma-Aldrich) at final concentration 0.01% (w/v). The optical tweezers platform (Impetux Optics S.L., Spain) is equipped with a direct force measurement device capable of detecting the change in optical momentum from the optical traps (SensoCell). The OT device is mounted on an inverted microscope (Eclipse Ti, Nikon, Japan), using a water immersion objective (Plan Apo VC 60XA/1.20 WI, Nikon) employed to focus the laser trap on the sample. The sample is placed in a custom-made glass chamber closed by a microscope slide. This chamber was mounted on the microscope and measurements were performed at 25 °C. Sample bright-field imaging was captured by a CMOS camera (ORCA-spark, Hamamatsu, Japan). The optical traps were operated with LightAce software developed using LabWiev (licenced by Impetux Optics S.L.). The cells got adhered to the bottom surface of the glass chamber by unspecific interactions. Only adhered cells with no morphological alterations were selected to perform the indentation routine. To constitute an indentation anvil, one indenter bead (active) and three supporting beads (passive) were placed in opposite quadrants next to the plasma membrane at axial positions approximately 2 μm above the glass surface. The exact diameter of the indenter bead used for probing was measured by image analysis using the ImageJ software. The indentation process consisted of pushing the cell laterally by generating an oscillation of the indenting bead. The fixed parameters of the oscillation are the shape (squared), frequency (0.5 Hz, which is enough to permit a complete relaxation between consecutive cycles) and offset (100%). The amplitude was varied in the linear regime, and the routine was set to sweep the 0.6-1.6 μm range with steps of 0.05 or 0.10 μm. Data were acquired during 45 s for each amplitude step. For each indentation experiment, the elasticity constant of the trap was calculated by using a particle scan routine included in the LightAce software. A Matlab (MathWorks, USA) script was written to analyze indentation data from the files obtained by the custom-made LabView indentation routine. Each force-time series is fitted to a creep-relaxation schema for every indentation cycle. The elasticity and permeability parameters were calculated from conventional rheological analysis. The elasticity parameter corresponds to the cortical stiffness (measured as a Young modulus), and the permeability parameter corresponds to the cytoplasm diffusivity (measured as a diffusion coefficient D).

### 4.28. *In vivo* xenograft tumorigenic assay

T3M4 ELOVL6-silenced (shELOVL6 #1 and shELOVL6 #2) and their non-target counterparts (shControl) cells were grown in 6- to 8-week-old female nude mice (Rj:ATHYM-Foxn1^nu/nu^, Janvier Laboratories). 5×10^5^ cells (100 μl: 50% PBS 1X, 50% matrigel) were subcutaneously implanted and growth was monitored using an electronic caliper. Tumor volume was calculated using the formula length x width x width. Albumin nanoparticles-bound paclitaxel (Abraxane^®^; Bristol Myers Squibb, Ref 660458) was administered intraperitoneally at a dose of 40 mg/kg twice per week at the indicated dates. In ELOVL6-IN-2 *in vivo* experiments, non-treated T3M4 cells were implanted in nude mice following the same protocol. The inhibitor was administered by oral gavage at a dose of 10 mg/kg at the indicated dates in combination with chemotherapy. Tumor volume was evaluated every 3-4 days. In the inducible CRISPRi system experiments, T3M4 cells carrying the construct were implanted in nude mice as mentioned. The system was activated using doxycycline diet in treatments of one week and resting another week in between. For all *in vivo* experiments, mice were housed according to institutional guidelines and all experimental procedures were performed in compliance with the institutional guidelines for the welfare of experimental animals approved by the Hospital 12 de Octubre Ethics Committee (CEI 20/377) and La Comunidad de Madrid (PROEX 312.8/21), and in accordance with the guidelines for Ethical Conduct in the Care and Use of Animals as stated in The International Guiding Principles for Biomedical Research involving Animals, developed by the Council for International Organizations of Medical Sciences (CIOMS).

### 4.29. Statistical analysis

All quantitative data are presented as mean ± SD (Standard Deviation) from ≥3 different biological replicates (n). Each biological replicate consists of three technical replicates.

Comparison of data that did not follow a normal distribution was performed using Mann-Whitney test. Comparison of data that followed a normal distribution was performed using ordinary one-way ANOVA test or t test. To analyze time-progressing experiments we performed a two-way ANOVA test. Significance was considered for *p ≤ 0.05, **p ≤ 0.01 and ***p ≤ 0.001. Software Prism 8.0 was used.

## 6. ACKNOWLEDGMENTS

The authors express gratitude to Francisco X. Real and Bruno Amati for providing essential reagents for the study. Maite Iglesias Badiola, Cruz Santos Tejedor and Alberto Lopez Rosado are acknowledged for their unwavering support to the group. Direna Alonso-Curbelo, Juan A. Recio and Ricardo Sanchez-Prieto are thanked for their valuable comments and for engaging in invaluable scientific discussions.

## Author Contributions

A.G.G. performed the *in vitro* experiments, aided by M.F.A., R.C.-G. and M.C.C. A.G.G. performed the *in vivo* experiments, aided by M.F.A., R.M.V. and R.C.-G. V.J.S.-A.L., G.V.P., A.G.T. and A.M.P. performed the bioinformatic analyses. A.G.G. and R.C.-G. were responsible for the FACS analysis. A.G.G. performed the immunofluorescence experiments, aided by D.C.L. and E.S.B. A.G.G., S.M.Q., C.P.R. and D.H.-A. performed the membrane rigidity experiments. C.L.R. and F.M.M. performed the indentation experiments. M.A.R. performed the electron microscopy experiments. A.G.G. was responsible for the statistical analysis. J.D. and J.V.S. performed the lipidomic analysis. JCG-C and A.L performed the characterization of FA metabolism and the detection of Abraxane intratumorally. R.T.-R. and S.R.-P. designed and produced the inducible CRISPRi and KO systems. J.L.R.P provided important reagents and infrastructures for the study. V.J.S.-A.L. designed and supervised the overall conduct of the study and obtained financial support. All authors have read and agreed to the published version of the manuscript.

## Institutional Review Board Statement

Animal studies were performed following protocols approved by the fully authorized animal facility of our institutions; approved by the Consejería de Medio Ambiente, Vivienda y Agricultura, Comunidad de Madrid (PROEX 312.8/21); and in accordance with EU Directive 2010/63.

## Data Availability Statement

The data presented in this study are available on GEO database with the BioProject ID PRJNA1063608.

---

**Supplementary Fig. 1.**
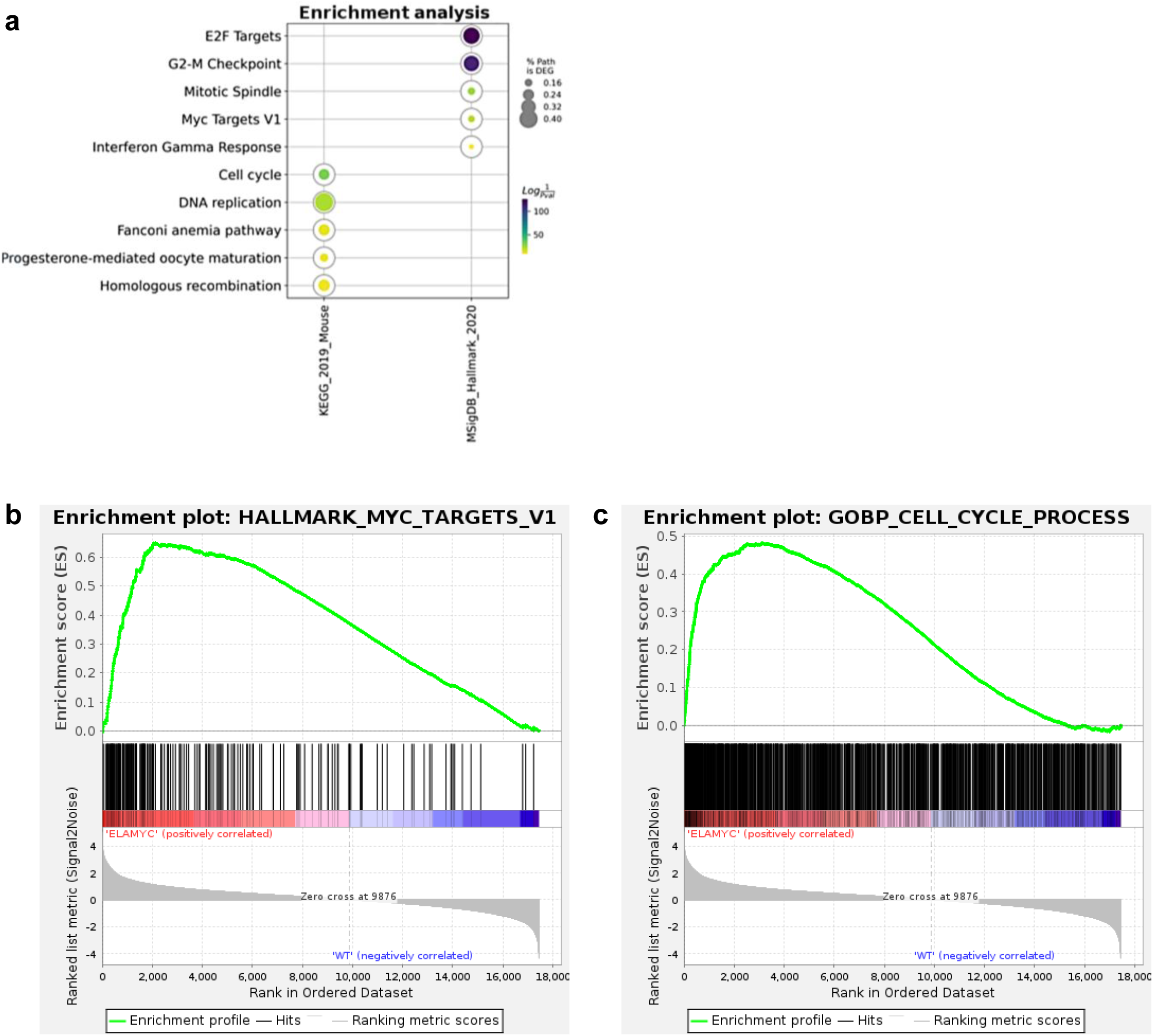
Pathways enriched in *Ela1-Myc* mouse model. **a** Functional enrichment analysis from upregulated genes using MisigDB Hallmark 2020 and KEGG 2019 signatures; RNA-seq data. **b** GSEA showing an enrichment in “Myc targets” pathway. **c** GSEA showing an enrichment in “cell cycle” pathway. NES Normalized enrichment score; FDR false-discovery rate.

**Supplementary Fig. 2.**
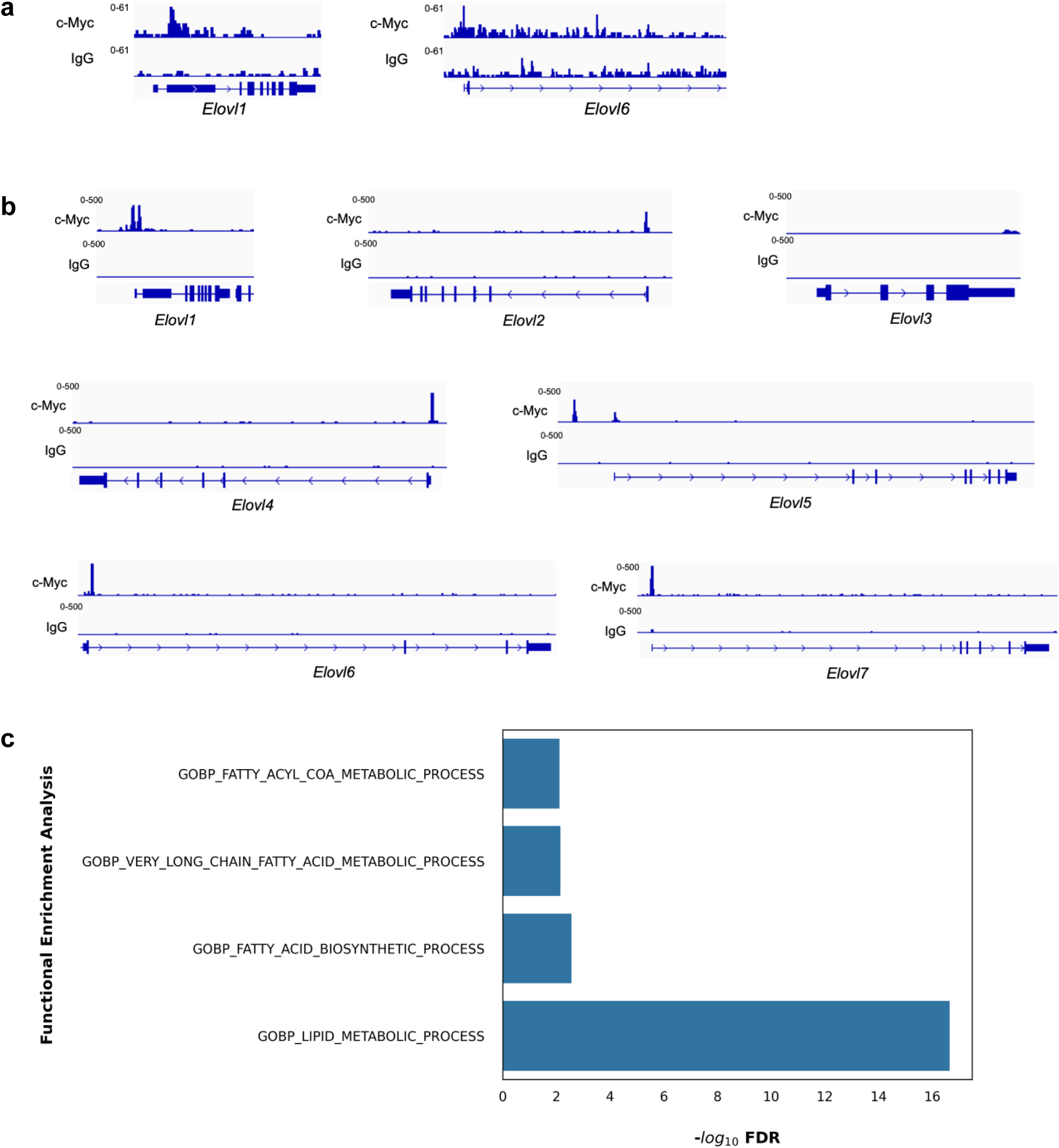
c-Myc ChIP-seq analyses in *Elovls* promoters. **a** ChIP-seq analysis showing c-Myc binding to the promoters of *Elovl1* and *Elovl6* in *Ela1-Myc* mouse model; IGV visualization of *Elovls* genes. **b** ChIP-seq analysis showing c-Myc binding to the promoters of the different *Elovls* in KPC mouse model; IGV visualization of *Elovls* genes. **c** Functional enrichment analysis of genes ascribed to peaks in (**S. 2b**).

**Supplementary Fig. 3.**
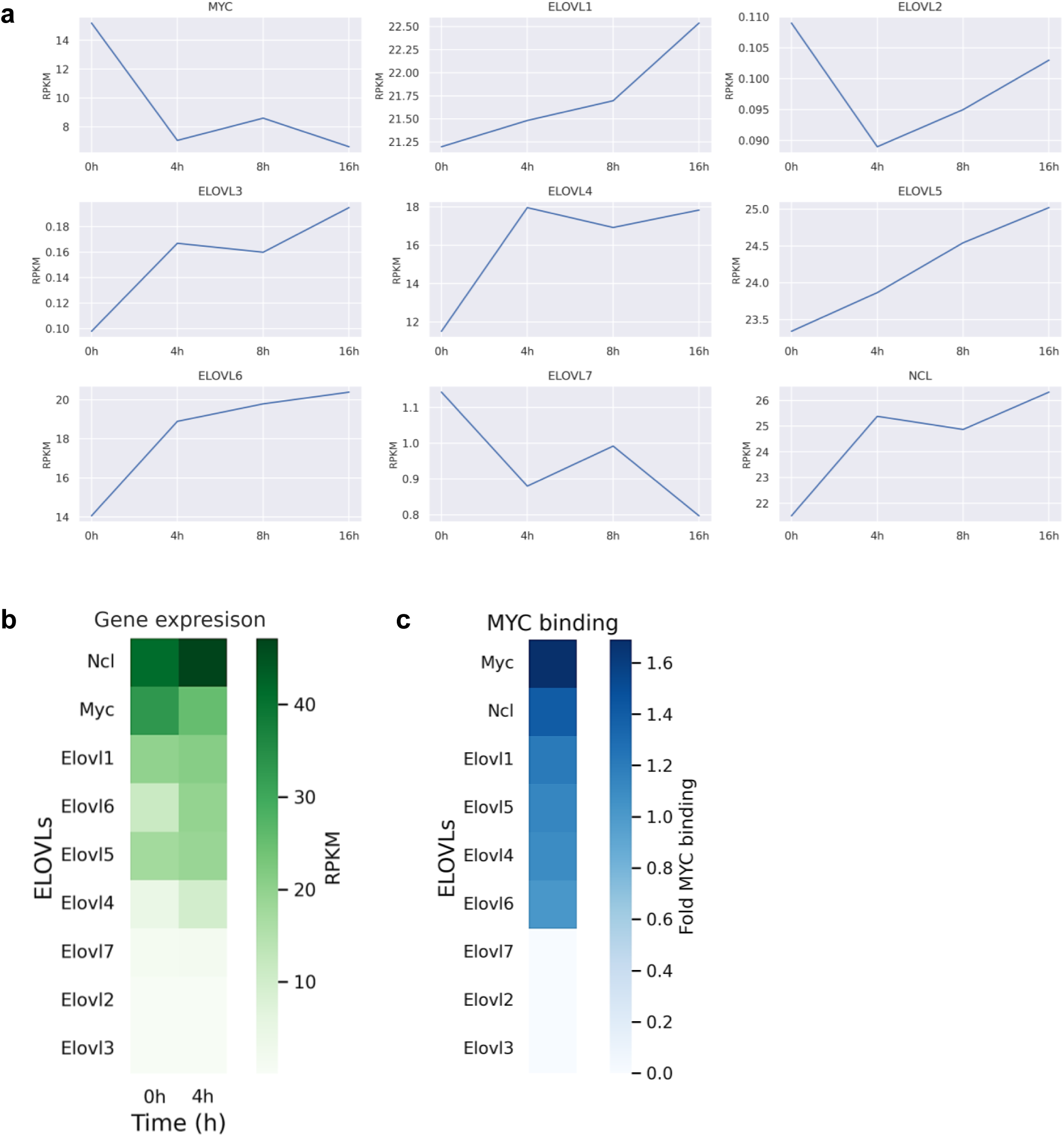
ELOVLs expression upon MYC-ER induction. **a** Mouse fibroblast 3T9 transduced with the inducible system Myc-ER were treated with 4-OHT during 2, 4, 8 and 16h; *Elovls*, *Myc* and *nucleolin* expression were evaluated by RNA-seq (RPKM values are shown). **b** Heatmap represents *Elovls*, *Myc* and *nucleolin* expression (RPKMs) at different timepoints. **c** Heatmap representing Myc binding to *Elovls, Myc* and *nucleolin* promoters at tumoral stage; the values correspond to Log_2_(ChIP-Input).

**Supplementary Fig. 4.**
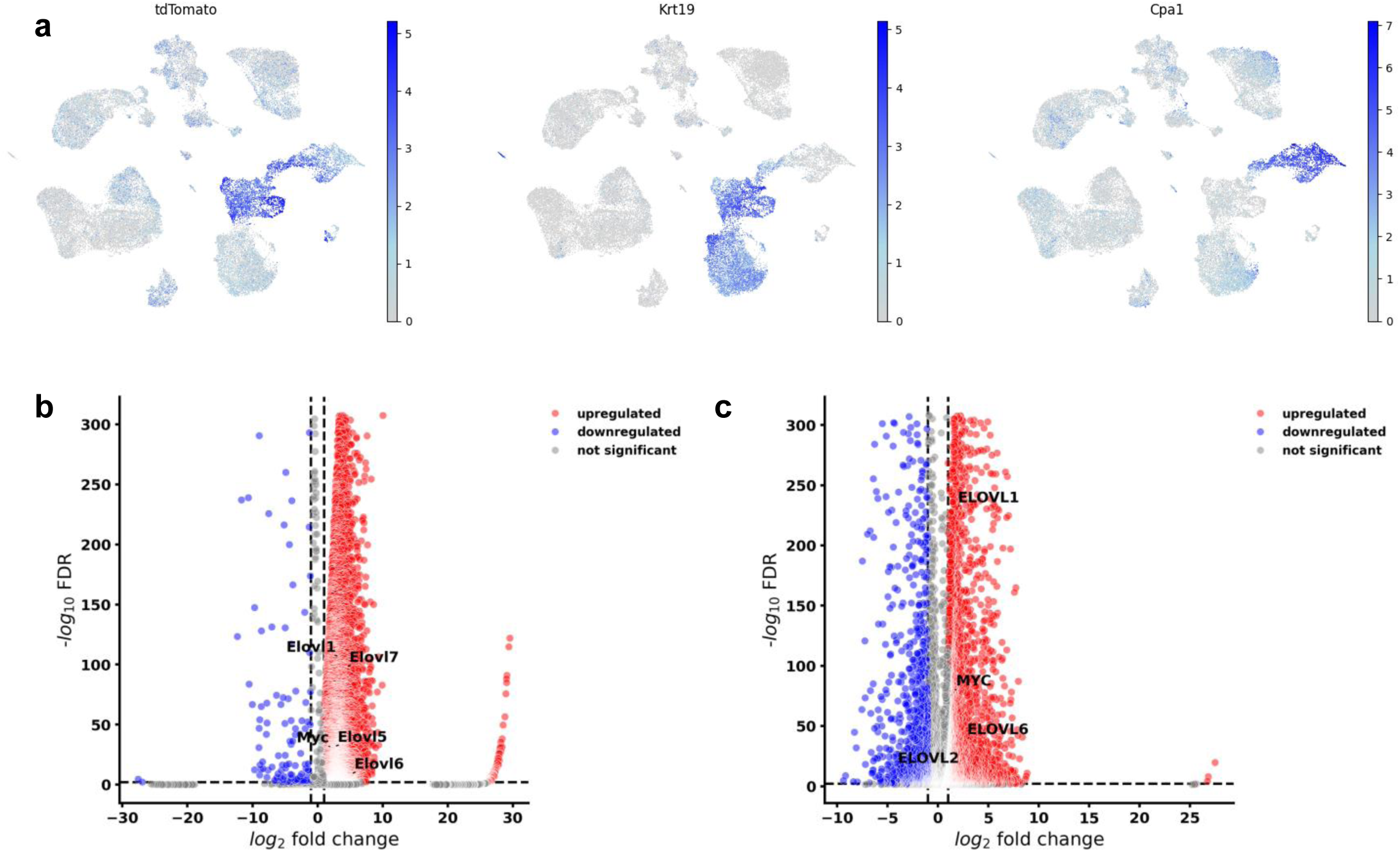
scRNA-Seq gene expression analysis. **a** UMAPs representing the expression per cell of *tdTomato*, *Krt19* and *Cpa1* on ductal, acinar early metaplastic and metaplastic cells; UMAP topology and cell types match the ones shown in (**1c**); scale bars show normalized (10^4^ counts per cell) and log-transformed counts. **b** Volcano plot showing DEGs between metaplastic and acinar cells in the tumoral progression mouse model shown in (**1c-e**). **c** Volcano plot showing DEGs between malign ductal and ductal cells in the PDAC human model shown in (**1f-h**).

**Supplementary Fig. 5.**
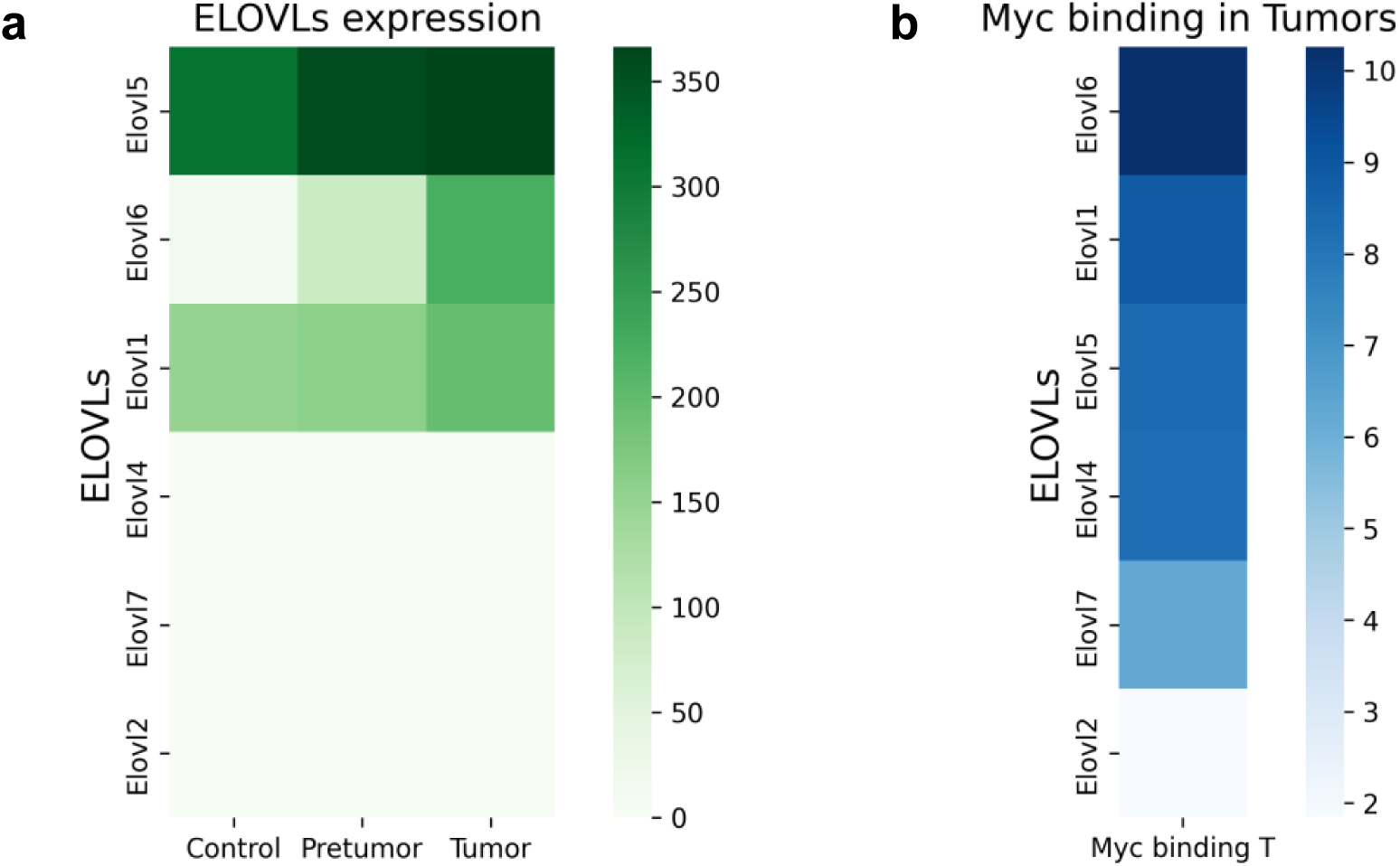
Myc genomic distribution and *Elovls* expression during B-cell lymphomagenesis *in vivo.* **a** B-cells from *Eμ-Myc* mice were collected at different stages (Control, Pre-tumoral and Tumor); heatmap represents *Elovls*, *Myc* and *nucleolin* expression (RPKMs) in Control, Pre-tumoral and Tumoral stages. **b** Heatmap representing Myc binding to *Elovls, Myc* and *nucleolin* promoters at tumoral stage; the values correspond to Log_2_(ChIP-Input).

**Supplementary Fig. 6.**
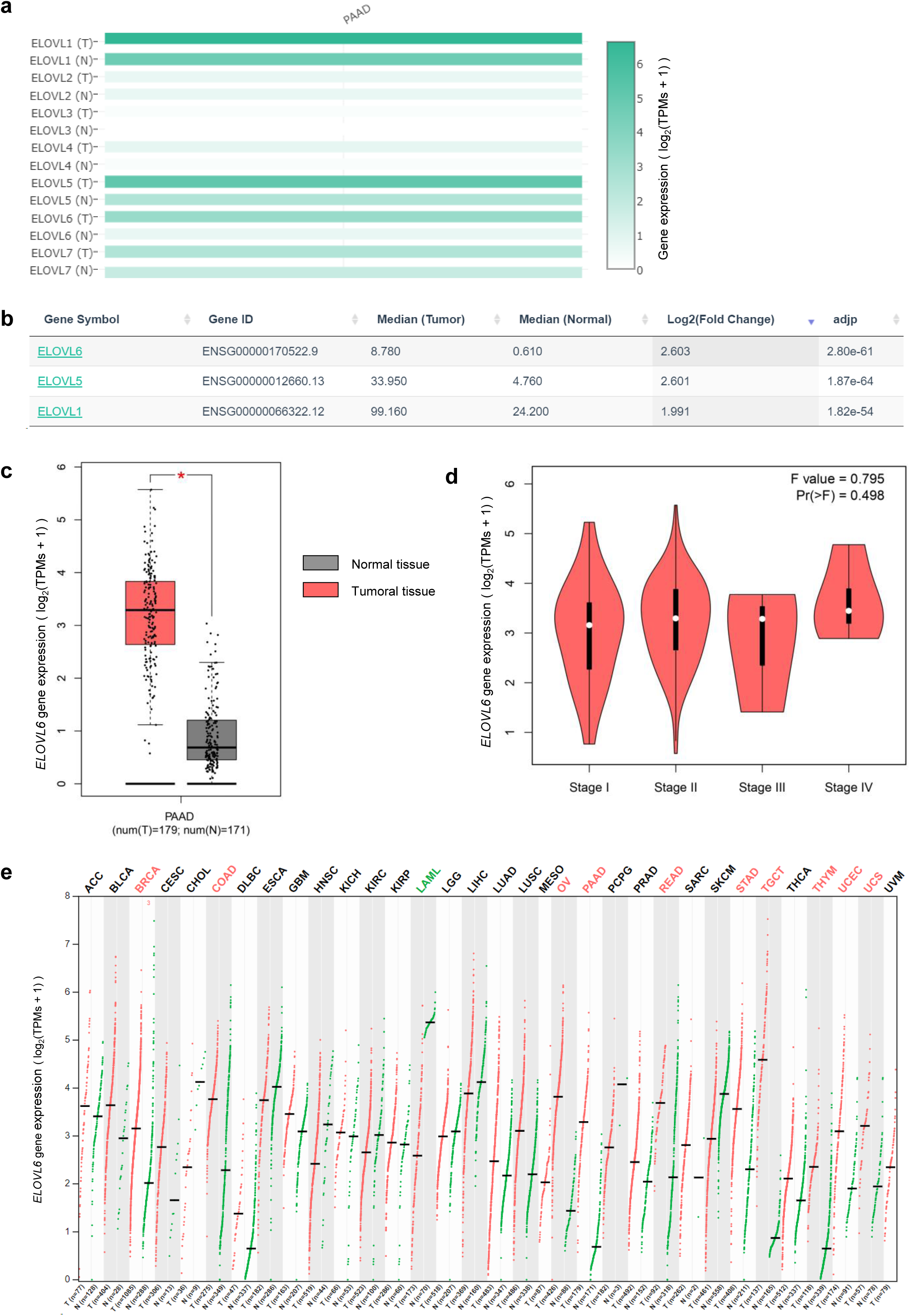
*ELOVLs* expression and alterations analysis in PDAC. **a** Gene expression of *ELOVLs* in pancreatic normal (N) and tumoral (T; PDAC) tissue; TCGA dataset; GEPIA browser; TPM: transcripts per million. **b** Significant differentially expressed *ELOVLs* in pancreatic normal and tumoral (PDAC) tissue; TCGA dataset; GEPIA browser; median expression is showed in transcripts per million. **c** *ELOVL6* gene expression in normal (N) and tumoral (T; PDAC) tissue; TCGA dataset; GEPIA browser; TPM: transcripts per million; logarithmic axis. **d** *ELOVL6* gene expression in different stages of PDAC progression; TCGA dataset; GEPIA browser; TPM: transcripts per million; logarithmic axis. **e** *ELOVL6* gene expression in normal (N) and tumoral (T) tissue; TCGA dataset; GEPIA browser; ACC: adrenocortical carcinoma; BLCA: bladder urothelial carcinoma; BRCA: breast invasive carcinoma; CESC: cervical squamous cell carcinoma and endocervical adenocarcinoma; CHOL: cholangio carcinoma; COAD: colon adenocarcinoma; DLBC: lymphoid neoplasm diffuse large B-cell lymphoma; ESCA: esophageal carcinoma; GBM: glioblastoma multiforme; HNSC: head and neck squamous cell carcinoma; KICH: kidney chromophobe; KIRC: kidney renal clear cell carcinoma; KIRP: kidney renal papillary cell carcinoma; LAML: acute myeloid leukemia; LGG: brain lower grade glioma; LIHC: liver hepatocellular carcinoma; LUAD: lung adenocarcinoma; LUSC: lung squamous cell carcinoma; MESO: mesothelioma; OV: ovarian serous cystadenocarcinoma; PAAD: pancreatic adenocarcinoma; PCPG: pheochromocytoma and paraganglioma; PRAD: prostate adenocarcinoma; READ: rectum adenocarcinoma; SARC: sarcoma; SKCM: skin cutaneous melanoma; STAD: stomach adenocarcinoma; TGCT: testicular germ cell tumors; THCA: thyroid carcinoma; THYM: thymoma; UCEC: uterine corpus endometrial carcinoma; UCS: uterine carcinosarcoma; UVM: uveal melanoma; TPM: transcripts per million; logarithmic axis.

**Supplementary Fig. 7.**
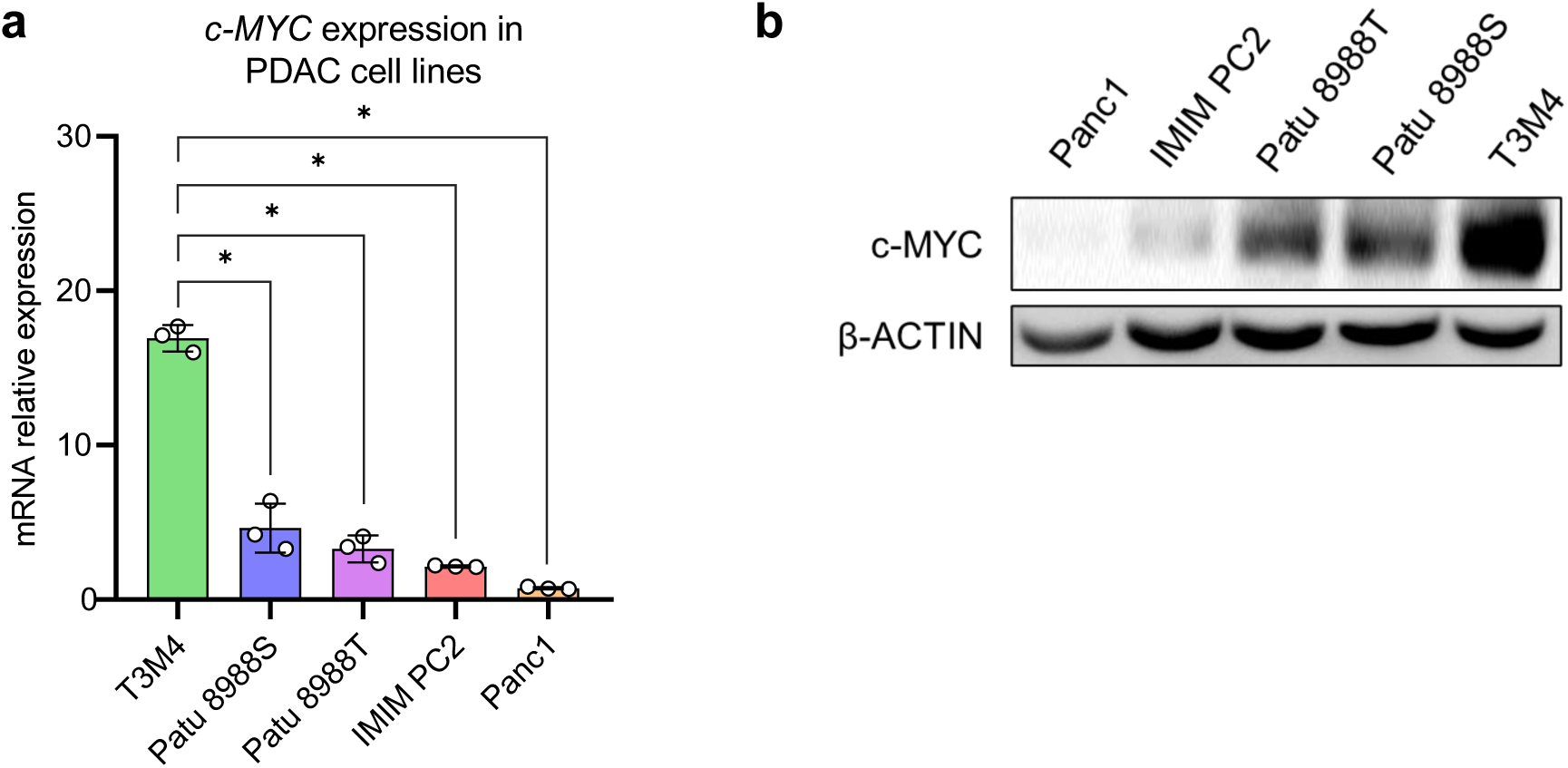
*c-MYC* expression in PDAC cell lines. **a** mRNA expression of *c-MYC* in PDAC cell lines; gene expression is normalized to *HPRT*; n = 3 per cell line. **b** Western blot of c-MYC in PDAC cell lines. All data are presented as mean ± SD; *p ≤ 0.05; (a) Mann-Whitney test. Source data are provided as a Source Data file.

**Supplementary Fig. 8.**
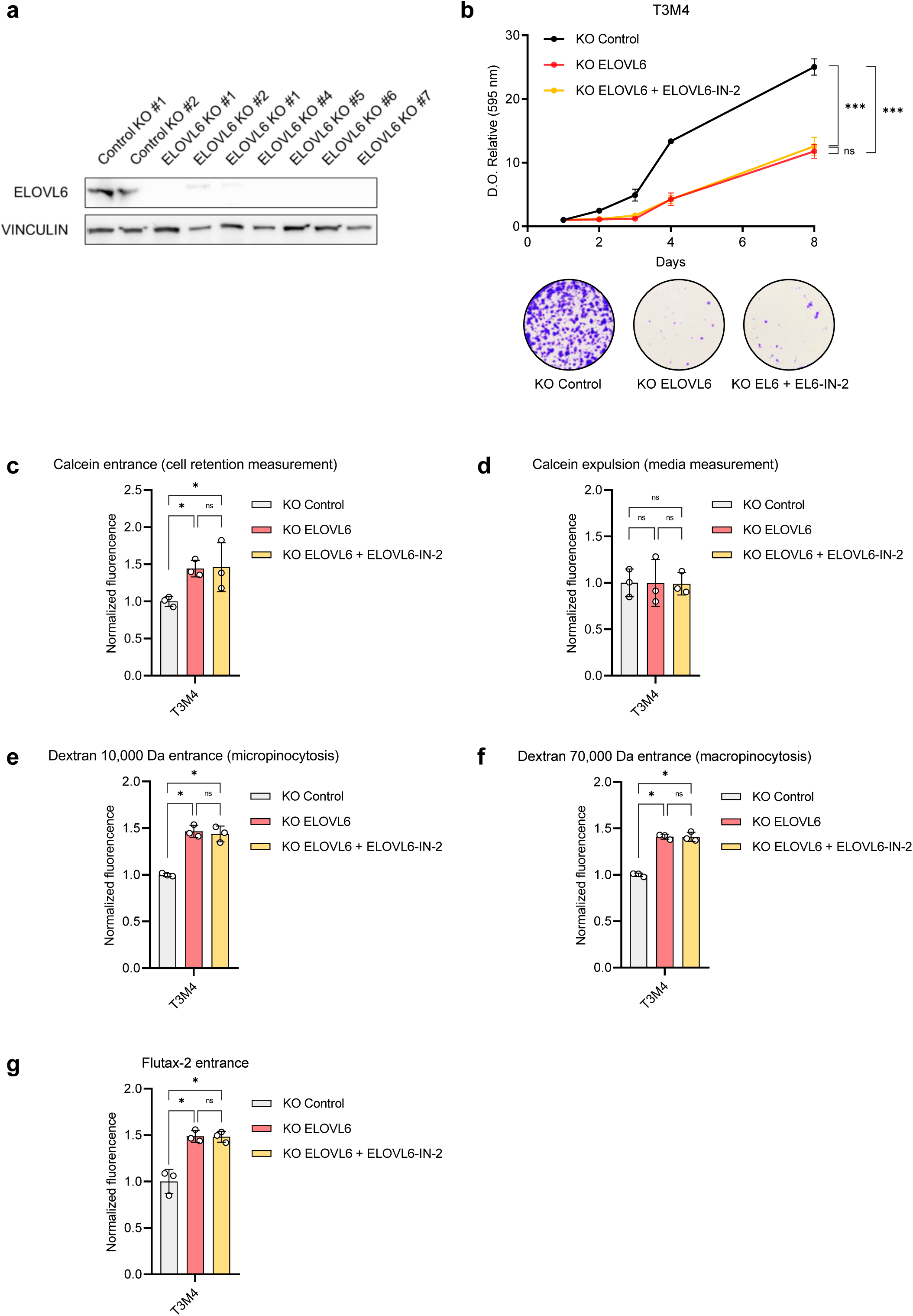
ELOVL6 chemical inhibition is specific. **a** Western blot showing ELOVL6 knockout (KO) in T3M4**. b** Proliferation and colony assays of ELOVL6 KO cells compared to KO control in T3M4; n = 3. **c** Normalized calcein-AM uptake and processing into calcein in ELOVL6 KO cells compared to KO control in T3M4; cell retention measurement; n = 3. **d** Normalized calcein expulsion in ELOVL6 KO cells compared to KO control in T3M4; media measurement; n = 3. **e** Micropinocytosis analysis by normalized fluorescence of dextran-rhodamine B 10,000 Da incorporated by ELOVL6 KO cells compared to KO control in T3M4; n = 3 per condition. **f** Macropinocytosis analysis by normalized fluorescence of dextran-rhodamine B 10,000 Da incorporated by ELOVL6 KO cells compared to KO control in T3M4; n = 3 per condition. **g** Normalized Flutax-2 uptake in ELOVL6 KO cells compared to KO control in T3M4; n = 3. All data are presented as mean ± SD; ns: not statistically significant, *p ≤ 0.05, ***p ≤ 0.001; (b) Two-way ANOVA test followed by Dunnett test; (c-g) Mann-Whitney test. Source data are provided as a Source Data file.

**Supplementary Fig. 9.**
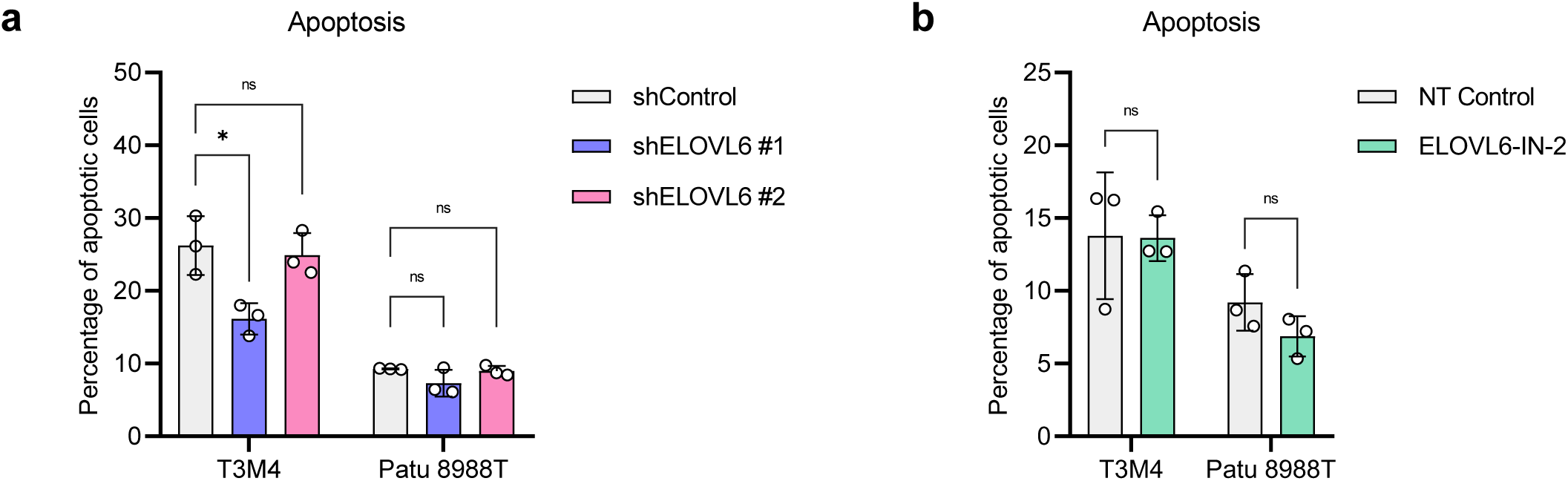
ELOVL6 interference causes no increase in apoptosis. **a** Apoptosis assay by FACS of *ELOVL6*-silenced cells compared to non-targeted control in PDAC cell lines; n = 3. **b** Apoptosis assay by FACS of ELOVL6-inhibited cells compared to non-treated control in PDAC cell lines; n = 3. All data are presented as mean ± SD; ns: not statistically significant, *p ≤ 0.05; (a, b) Mann-Whitney test. Source data are provided as a Source Data file.

**Supplementary Fig. 10.**
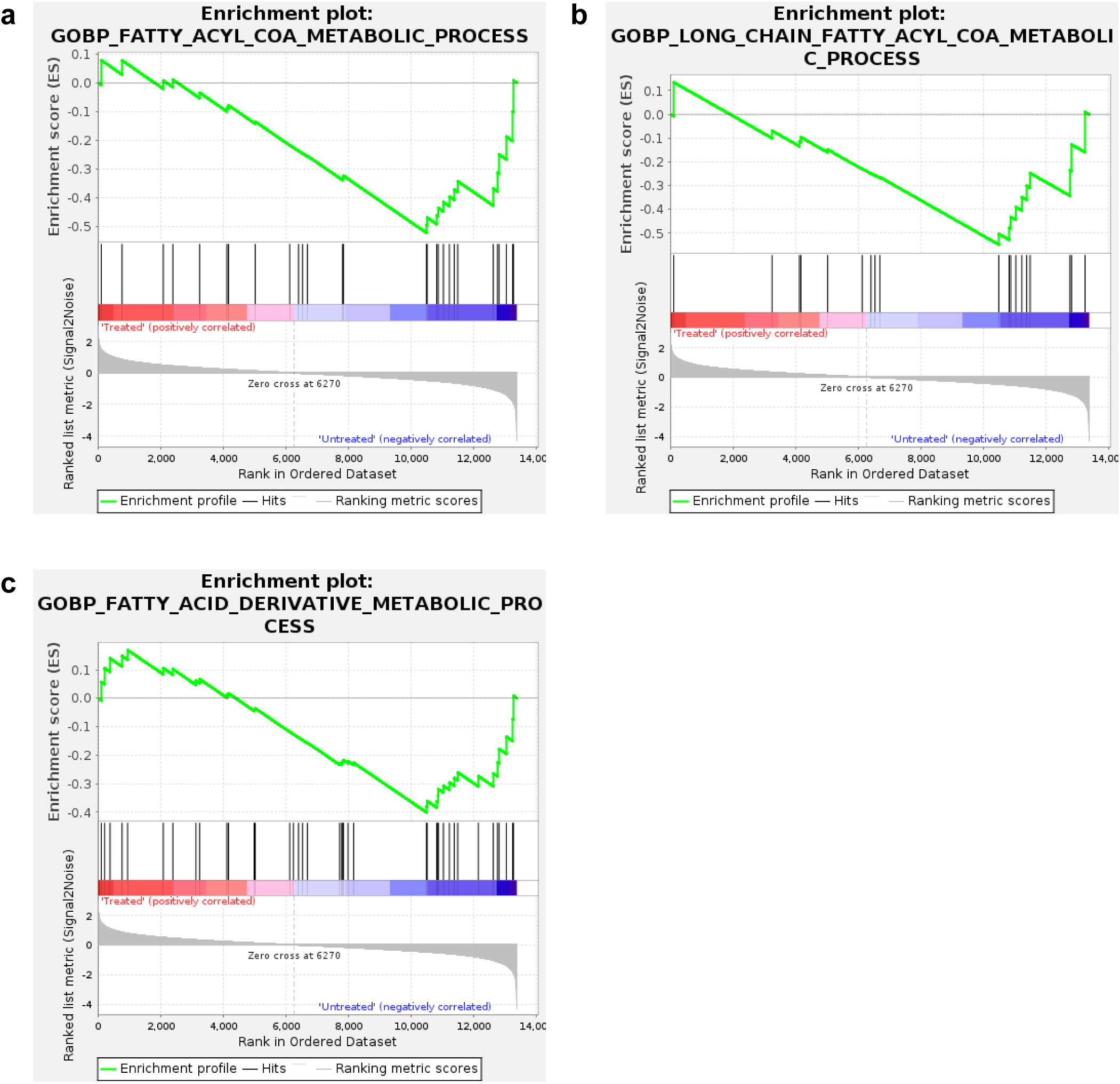
ELOVL6 inhibition causes lipid metabolism downregulation. **a** GSEA of “fatty acyl-CoA metabolic process GOBP”. **b** GSEA of “long-chain fatty acyl-CoA metabolic process GOBP”. **c** GSEA of “fatty acid derivative metabolic process GOBP”.

**Supplementary Fig. 11.**
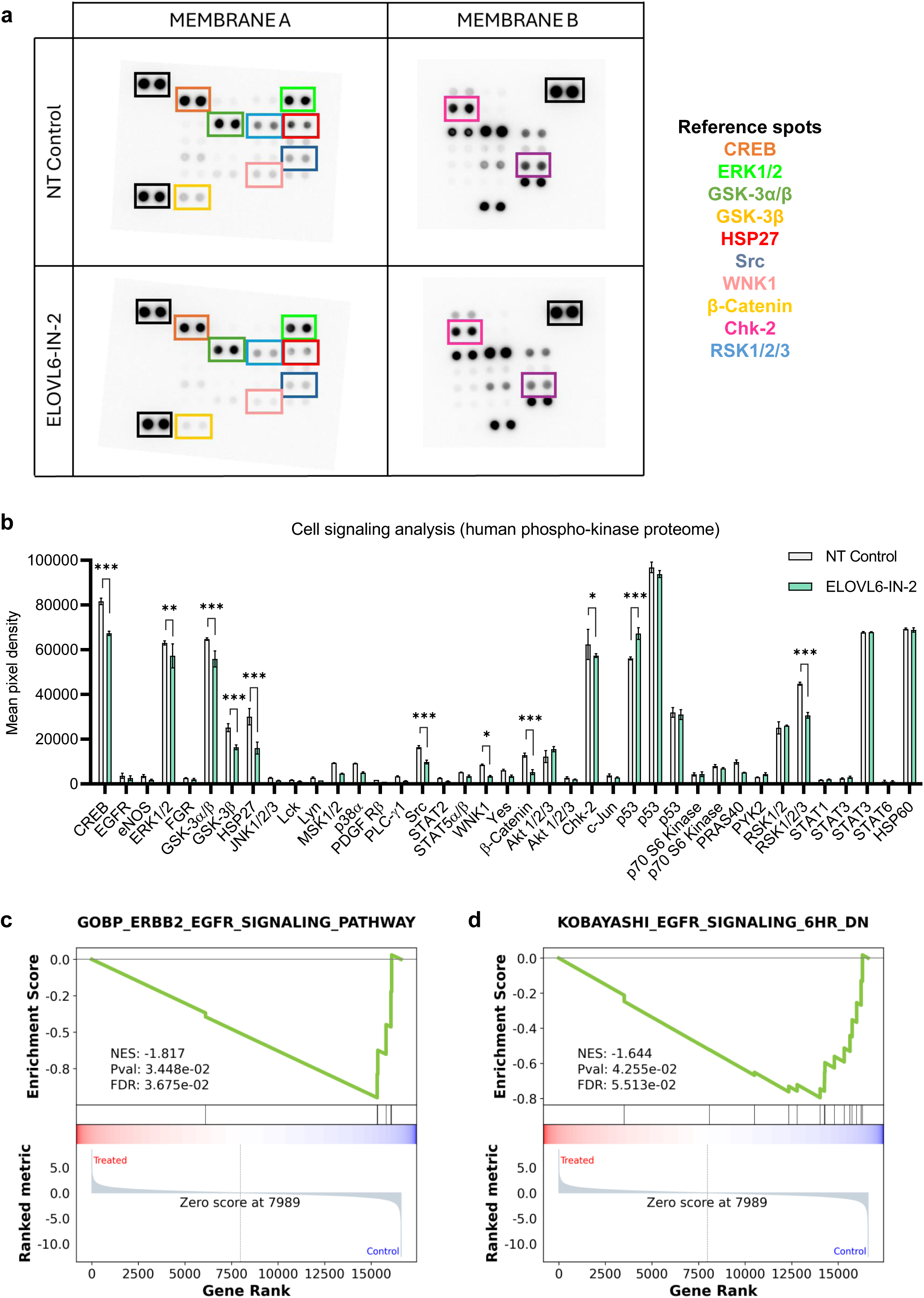
ELOVL6 inhibition decreases cell signaling. **a** Proteome human phospho-kinase array of ELOVL6-inhibited cells compared to non-treated control in T3M4; highlighted proteins present a statistically significant difference. **b** Cell signaling diminishment upon ELOVL6 inhibition. **c** GSEA of “GOBP ERBB2 EGFR signaling pathway”. **d** GSEA of “Kobayashi EGFR signaling 6hr DN”. All data are presented as mean ± SD; *p ≤ 0.05, **p ≤ 0.01, ***p ≤ 0.001; (b) Two-way ANOVA test followed by Sidak test. Source data are provided as a Source Data file.

**Supplementary Fig. 12.**
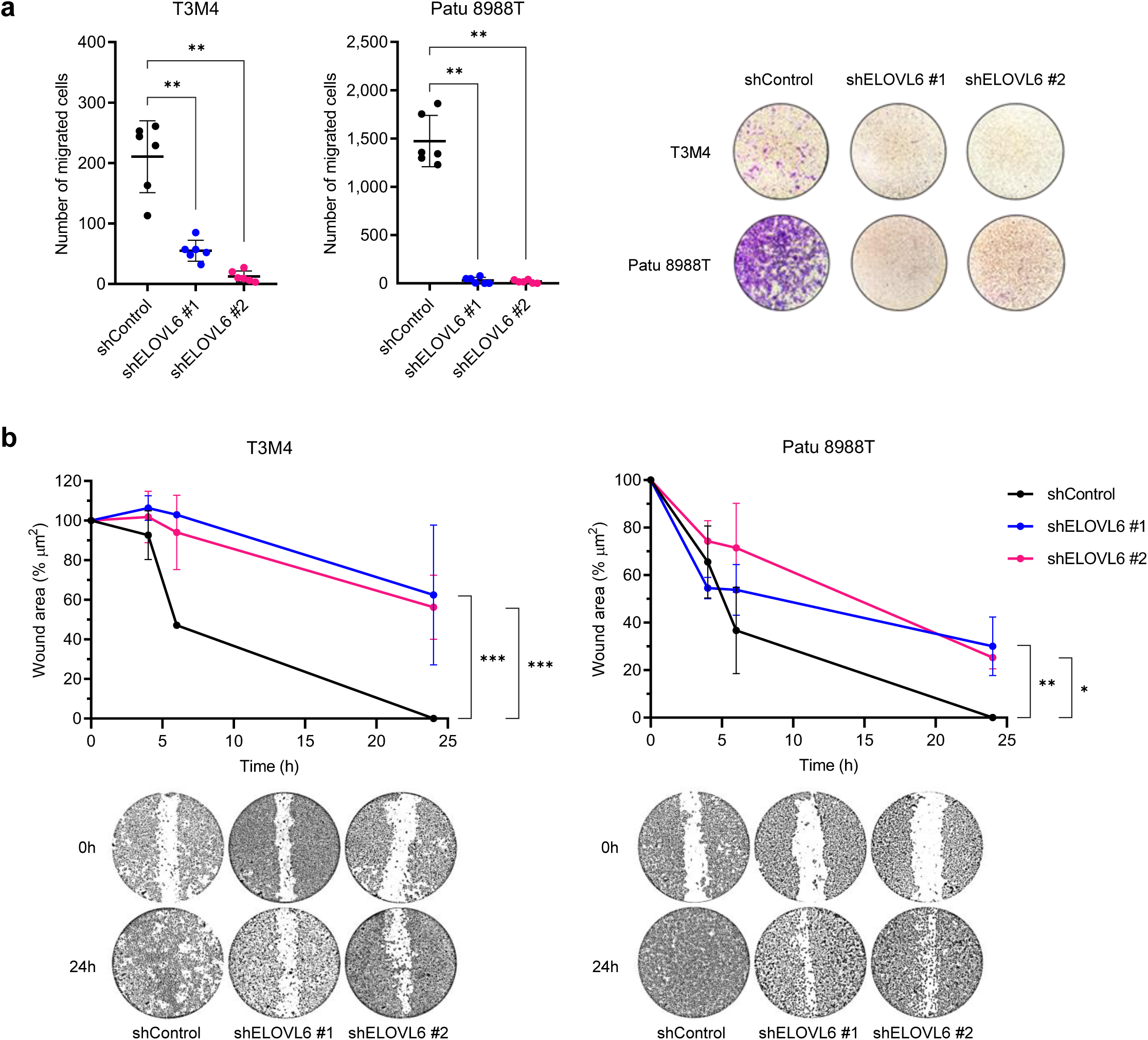
ELOVL6 interference decreases cell migration *in vitro*. **a** Transwell assay of *ELOVL6*-silenced cells compared to non-targeted control in PDAC cell lines; n = 6; representative images of transwell surfaces are shown. **b** Wound healing assay of *ELOVL6*-silenced cells compared to non-targeted control in PDAC cell lines; n = 3; representative images of wound closure are shown. All data are presented as mean ± SD; *p ≤ 0.05, **p ≤ 0.01, ***p ≤ 0.001; (a) Mann-Whitney test; (b) Two-way ANOVA test followed by Dunnett test. Source data are provided as a Source Data file.

**Supplementary Fig. 13.**
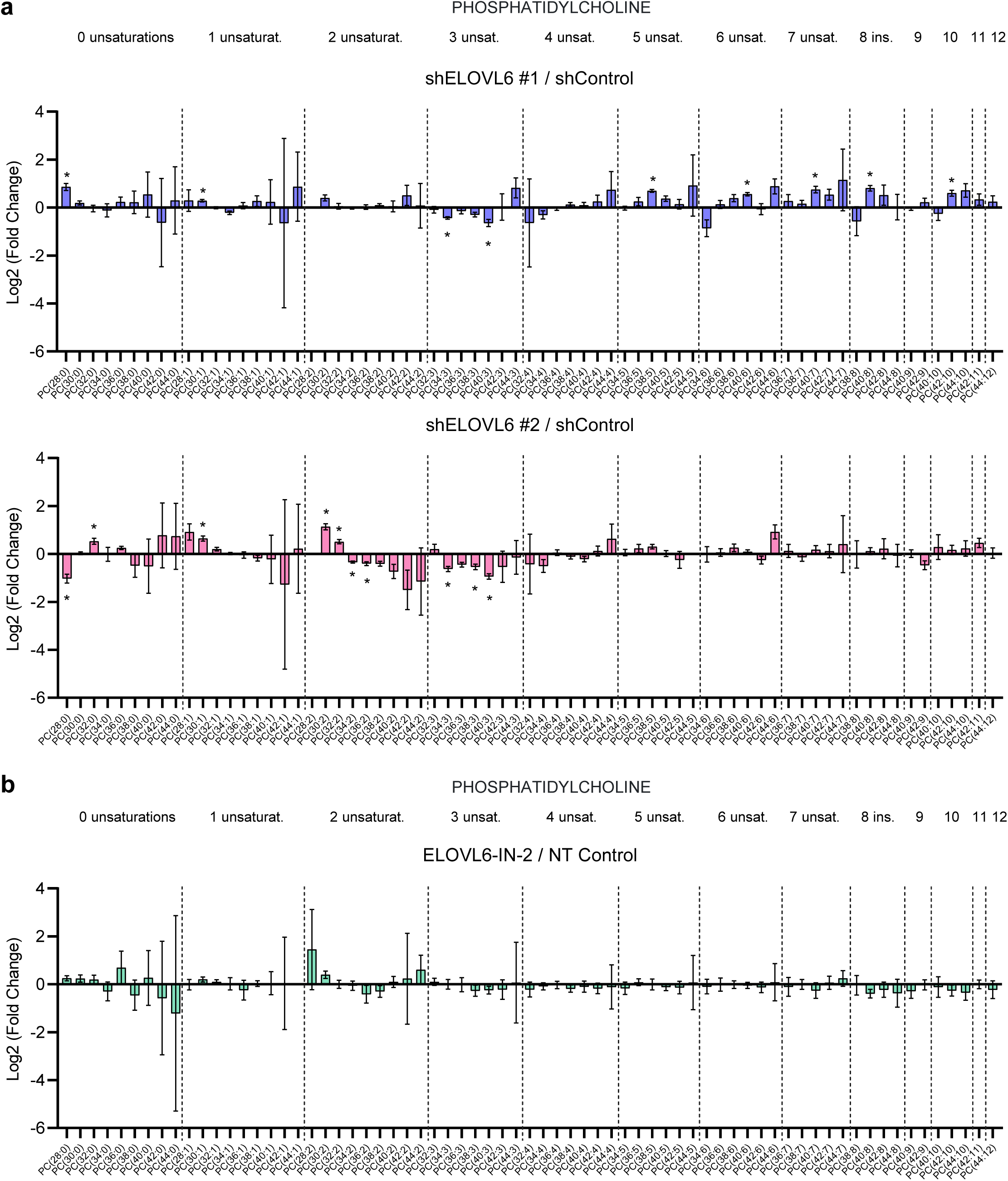
ELOVL6 interference modifies lipid composition. **a** Mass spectrometry lipidomic analysis of phosphatidylcholine in *ELOVL6*-silenced cells compared to non-targeted control in T3M4; separated regions represent fatty acids with the same saturation state and carbons’ number increases along each region; n = 3. **b** Mass spectrometry lipidomic analysis of phosphatidylcholine in ELOVL6-inhibited cells compared to non-treated control in T3M4; n = 4. Source data are provided as a Source Data file.

**Supplementary Fig. 14.**
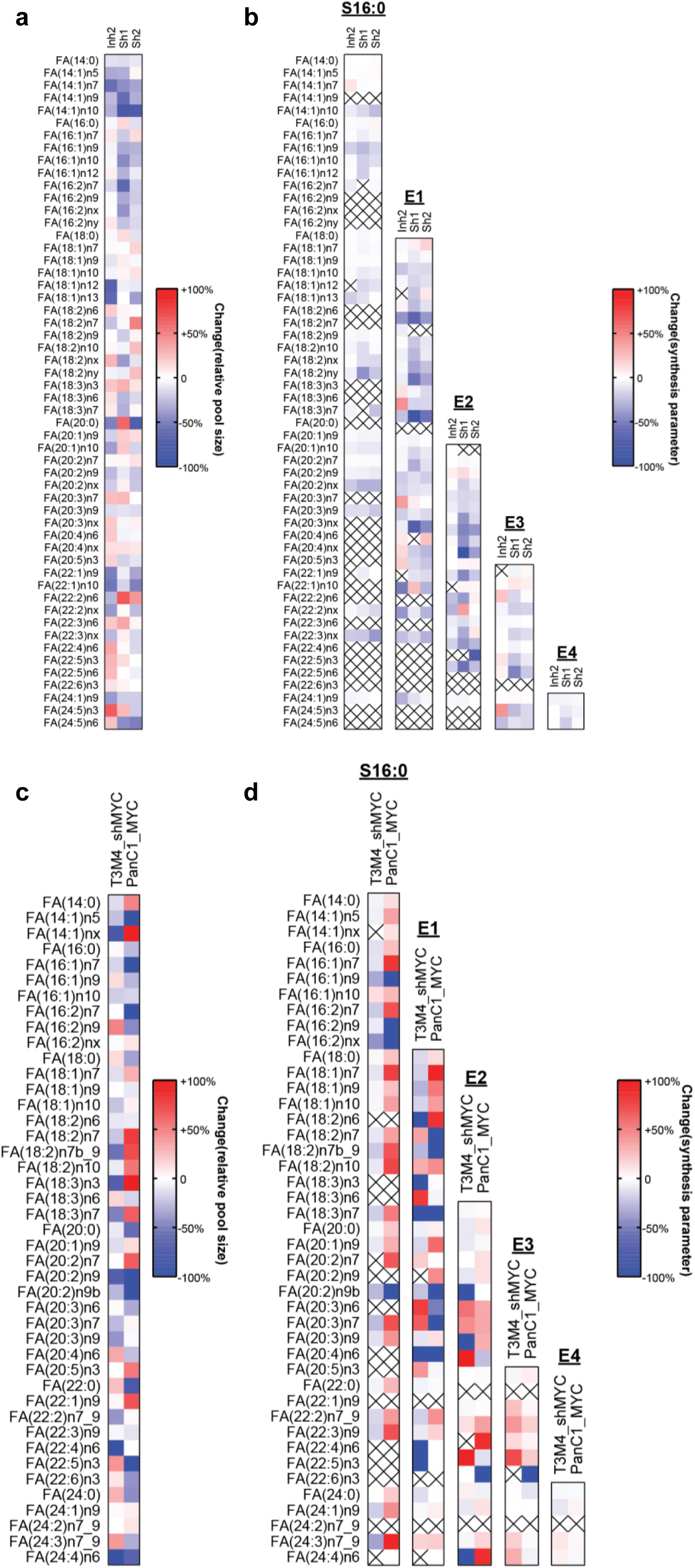
Analysis of fatty acid metabolism. **a** Heatmap showing for each detected fatty acid the mean value of the change (versus NT Control for ELOVL6-IN-2 and vs shControl for shELOVL6 #1 and shELOVL6 #2) in the relative pool size, expressed as percentage. **b** Heatmap showing the mean value the change (versus NT Control for ELOVL6-IN-2 and vs shControl for shELOVL6 #1 and shELOVL6 #2) for each detected fatty acid in the following synthesis parameters: calculated S, E1, E2, E3 and E4, expressed as percentage. **c** Heatmap showing for each detected fatty acid the mean value of the change (versus non-target shRNA for T3M4 shMYC and versus PANC1 empty vector for PANC1 MYC) in the relative pool size, expressed as percentage. **d** Heatmap showing the mean value the change (versus non-target shRNA for T3M4 shMYC and versus PANC1 empty vector for PANC1 MYC) for each detected fatty acid in the following synthesis parameters: calculated S, E1, E2, E3 and E4, expressed as percentage.

**Supplementary Fig. 15.**
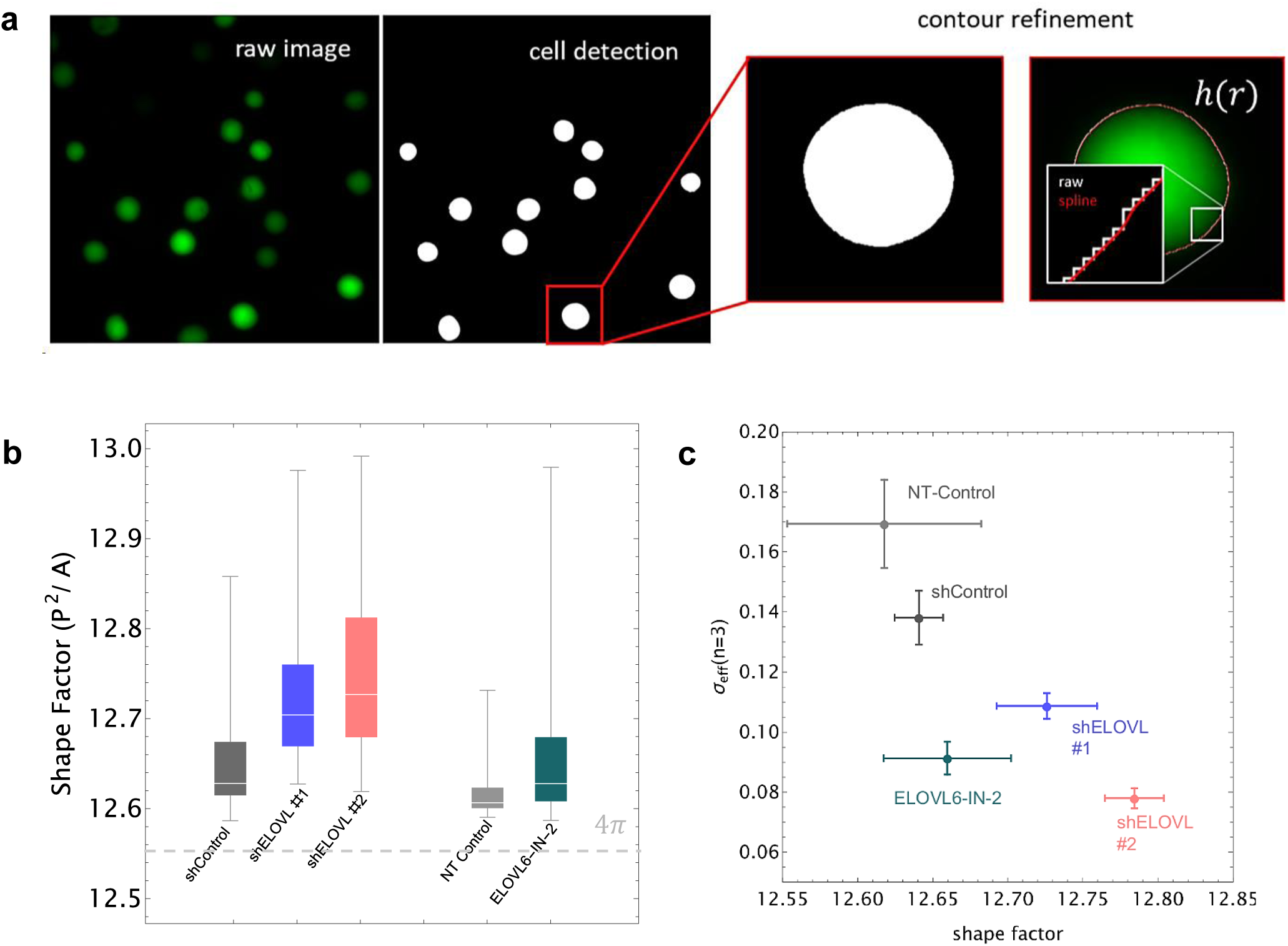
Description of the computational rationale used for membrane rigidity estimation. **a** Raw fluorescence microscopy images are noise-reduced and eventually binarized. The resulting morphological components are selected based on their mean size and circularity to ensure that the analysed images correspond to a 50% homogeneous sample. The raw contours are refined using a B-Spline interpolation and then transformed to polar coordinates. **b** Cell surface shape variability is analyzed in terms of surface excess (shape factor, P^2^/A) and membrane self-deformability, and eventually interpreted in terms of effective surface tension through the evaluation of the Fourier amplitude for the third equatorial mode wavevector (k) in the tension-dominated regime. **c** Panel c shows the averaged values of the shape factor and effective surface tension for the experimental groups.

**Supplementary Fig. 16.**
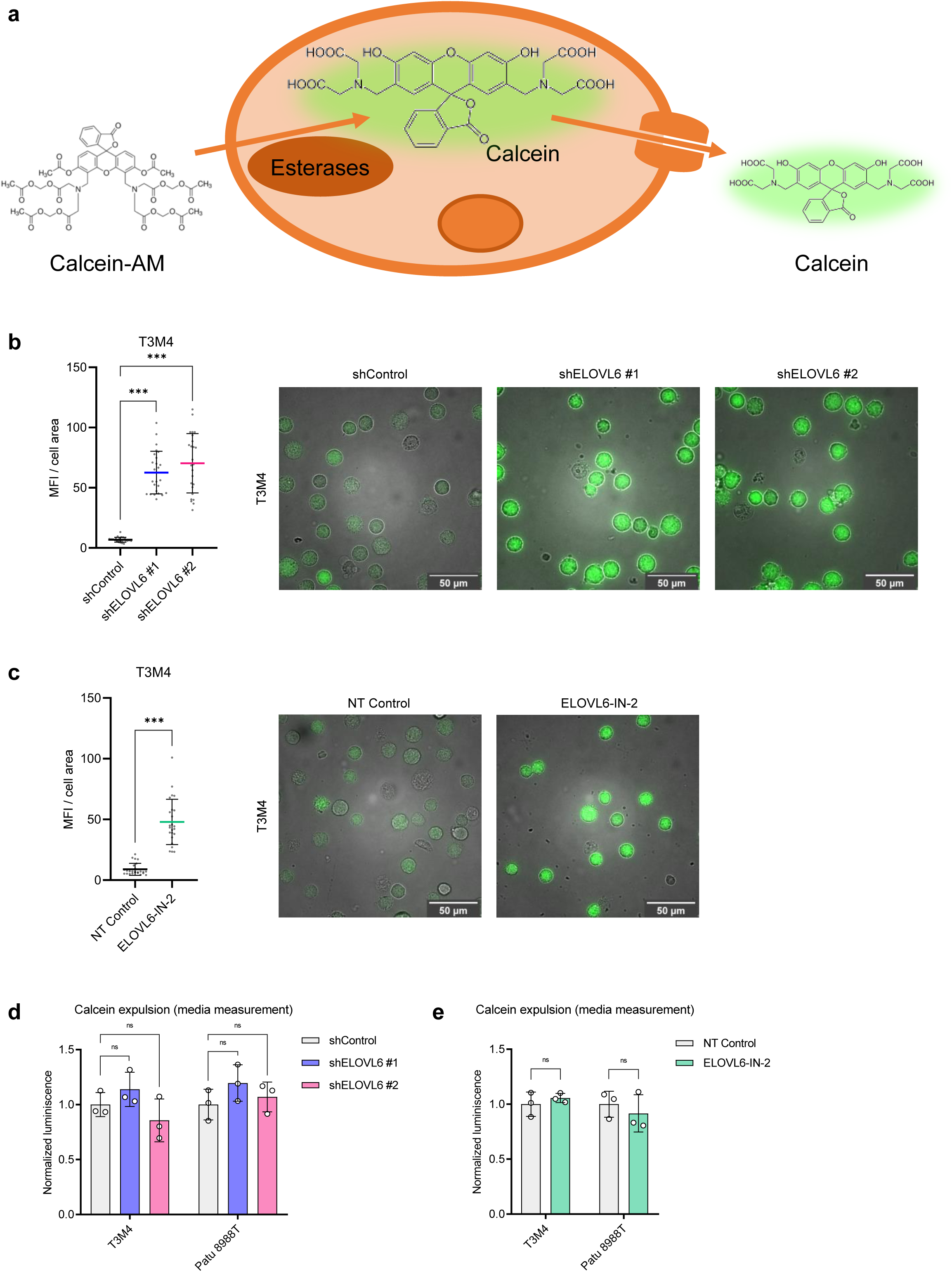
Calcein entrance and expulsion in PDAC cell lines. **a** Scheme of calcein entrance and expulsion mechanisms. **b** Quantification and representative images of immunofluorescence staining for calcein (green) uptake in *ELOVL6*-silenced cells compared to non-targeted control in T3M4; n = 3 per condition. **c** Quantification and representative images of immunofluorescence staining for calcein (green) uptake in ELOVL6-inhibited cells compared to non-treated control in T3M4; n = 3 per condition. **d** Normalized calcein expulsion in *ELOVL6*-silenced cells compared to non-targeted control and ELOVL6-inhibited cells compared to non-treated control in PDAC cell lines; media measurement; n = 3. All data are presented as mean ± SD; ns: not statistically significant, ***p ≤ 0.001; (b) Ordinary one-way ANOVA test followed by Dunnett test; (c) Unpaired t test; (d) Mann-Whitney test. Source data are provided as a Source Data file.

**Supplementary Fig. 17.**
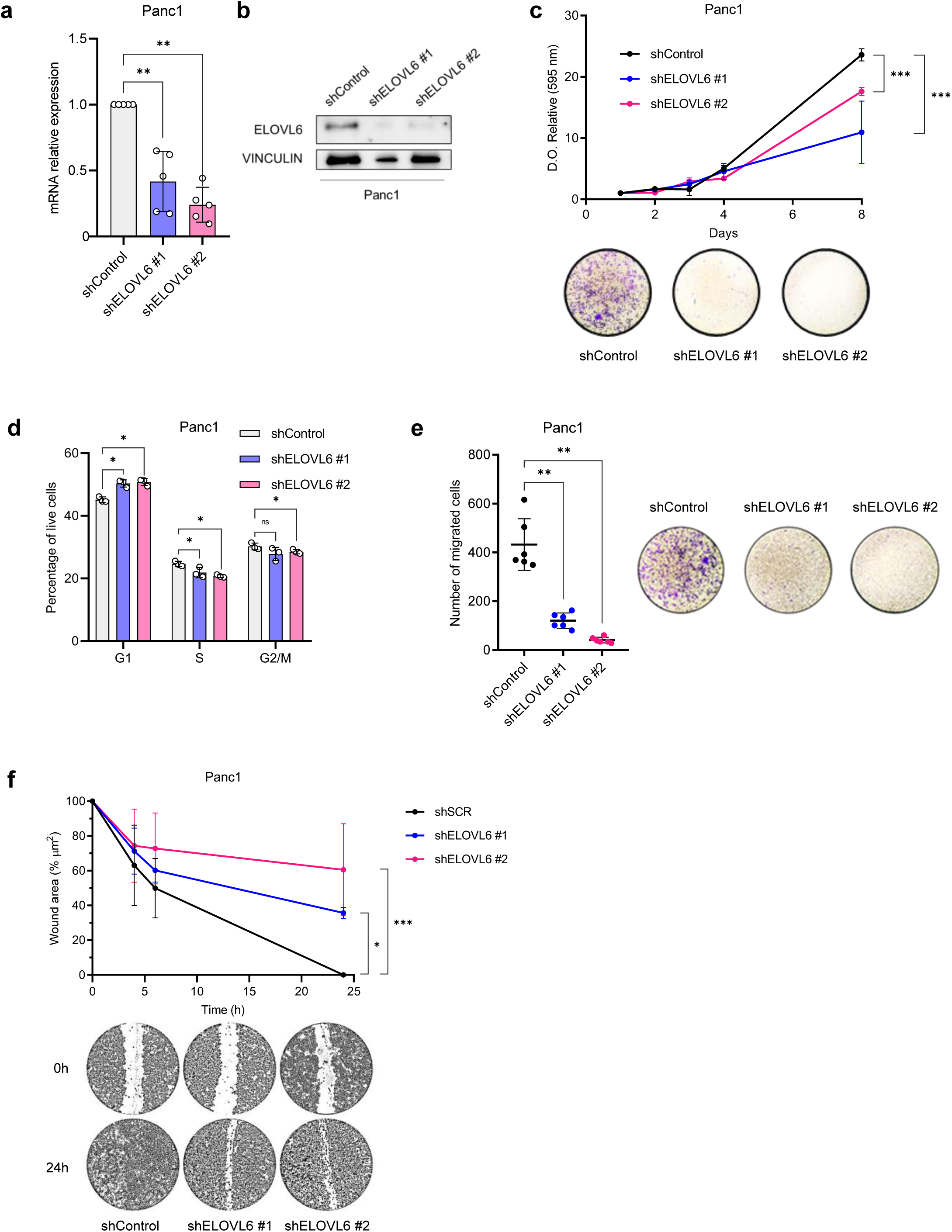
ELOVL6 interference decreases cell proliferation and migration *in vitro* in Panc1 cell line. **a** mRNA expression silencing of *ELOVL6* in Panc1; gene expression is normalized to *HPRT* and shControl; n = 5 per genotype and cell line. **b** Western blot showing ELOVL6 silencing in Panc1. **c** Proliferation and colony assays of *ELOVL6*-silenced cells compared to non-targeted control in Panc1; n = 3. **d** Cell cycle assay by FACS of *ELOVL6*-silenced cells compared to non-targeted control in Panc1; n = 3. **e** Transwell assay of *ELOVL6*-silenced cells compared to non-targeted control in Panc1; n = 6; representative images of transwell surfaces are shown. **f** Wound healing assay of *ELOVL6*-silenced cells compared to non-targeted control in Panc1; n = 3; representative images of wound closure are shown. All data are presented as mean ± SD; ns: not statistically significant, *p ≤ 0.05, **p ≤ 0.01, ***p ≤ 0.001; (a, d, e) Mann-Whitney test; (c, f) Two-way ANOVA test followed by Dunnett test. Source data are provided as a Source Data file.

**Supplementary Fig. 18.**
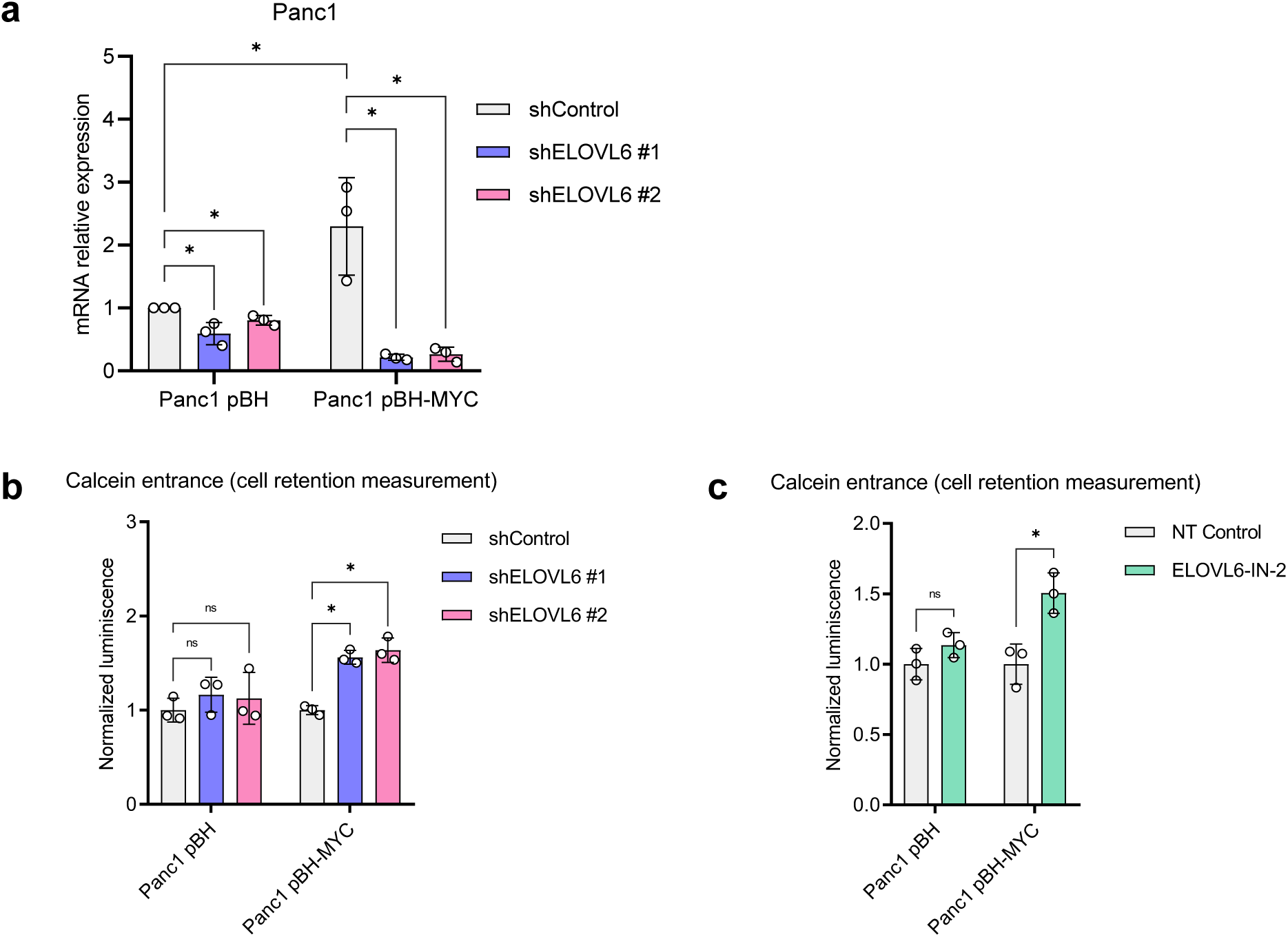
ELOVL6 interference efficacy is dependent on *c-MYC* expression levels. **a** mRNA expression of *ELOVLs* in Panc1 (pBH and pBH-MYC); gene expression is normalized to *HPRT* and shControl in Panc1-pBH; n = 3 per genotype. **b** Normalized calcein-AM uptake and processing into calcein in *ELOVL6*-silenced cells compared to non-targeted control in Panc1 (pBH and pBH-MYC); cell retention measurement; n = 3. **c** Normalized calcein-AM uptake and processing into calcein in ELOVL6-inhibited cells compared to non-treated control in Panc1 (pBH and pBH-MYC); cell retention measurement; n = 3. All data are presented as mean ± SD; ns: not statistically significant, *p ≤ 0.05; (a-c) Mann-Whitney test. Source data are provided as a Source Data file.

**Supplementary Fig. 19.**
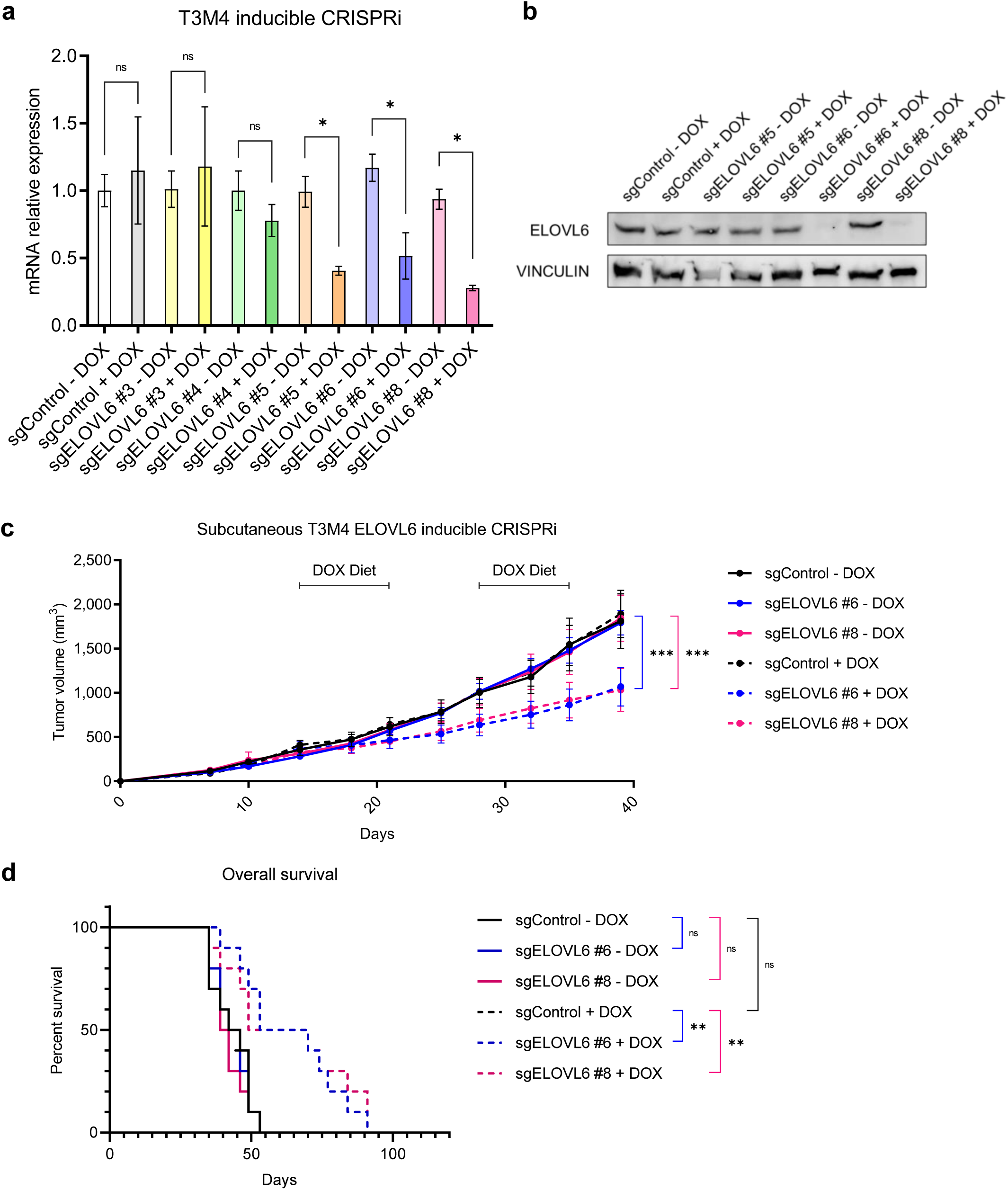
ELOVL6 interference reduces tumor growth *in vivo*. **a** mRNA expression of *ELOVL6* in T3M4 inducible CRISPRi; gene expression is normalized to *HPRT* and sgControl - DOX; n = 3 per genotype. **b** Western blot showing *ELOVL6* downregulation in T3M4 inducible CRISPRi. **c** Tumor growth in nude mice subcutaneously implanted with T3M4 inducible CRISPRi; doxycycline diet was administered for one week periods whit weekly rests when tumor volume reached 500 mm^3^; n = 10. **d** Survival analysis of the mice in (**S. 18c**). All data are presented as mean ± SD; ns: not statistically significant, *p ≤ 0.05, ***p ≤ 0.001; (a) Unpaired T test followed by T-Welch test; (c) Two-way ANOVA test followed by Dunnett test (c); (d) Log-rank (Mantel-Cox) test. Source data are provided as a Source Data file.

**Figure SN1.**
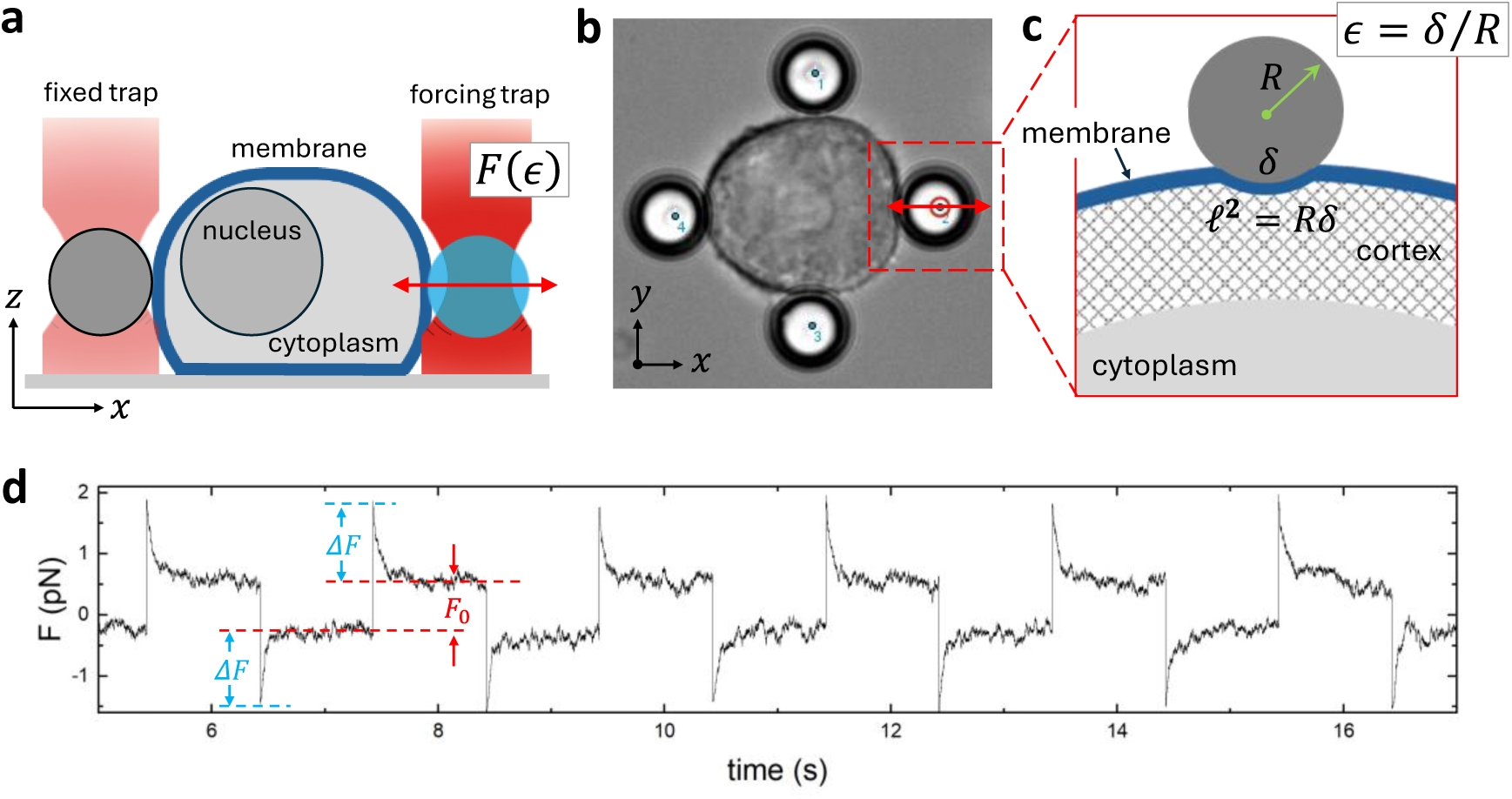
Optical tweezing (OT)-indentation implemented in horizontal geometry. **a** Schematics of the experimental OT setup for lateral cell forcing in a trapping anvil with four fixing beads (horizontal view). The indenting bead acts as a contact probe for the responsive force *F*(*ɛ*; *t*) under indentation strain amplitude *ɛ* ≡ *δ*⁄*R* (right sphere coloured in blue). This deformation stress is measured under slow deformation frequency allowing for poroelastic relaxation in terms of oscillatory indentation strain *ɛ*(*t*) ≅ (*δ*⁄*R*)*cos*(*ωt*) (at *ω* = 0.5 Hz; being *R* the bead radius). **b** Control T3M4 cell trapped in the OT-anvil under lateral indentation forcing (zenithal view). **c** Detail on the cortical contact of a large spherical bead on the plasma membrane and underlying cortex at small indentation strain resulted into the linear contact line ℓ ≅ √*δR* ≈ *Rɛ*^1⁄2^ (since *R* = 5.7*μm*, thus *ɛ* ≪ 1 and ℓ ≪ *R*). **d** Typical force time series for a T3M4 cell undergoing poroelastic relaxation under lateral OT-indentation causing initial permeability overshot of amplitude Δ*F* ≅ Δ*P*ℓ^2^(being Δ*P* a pressure drop), followed by terminal force relaxation into surface rigidness *F*_0_ ≅ *E*_0_ℓ^2^ (being *E*_0_ the Young modulus at the cortical area contact).

**Figure SN2.**
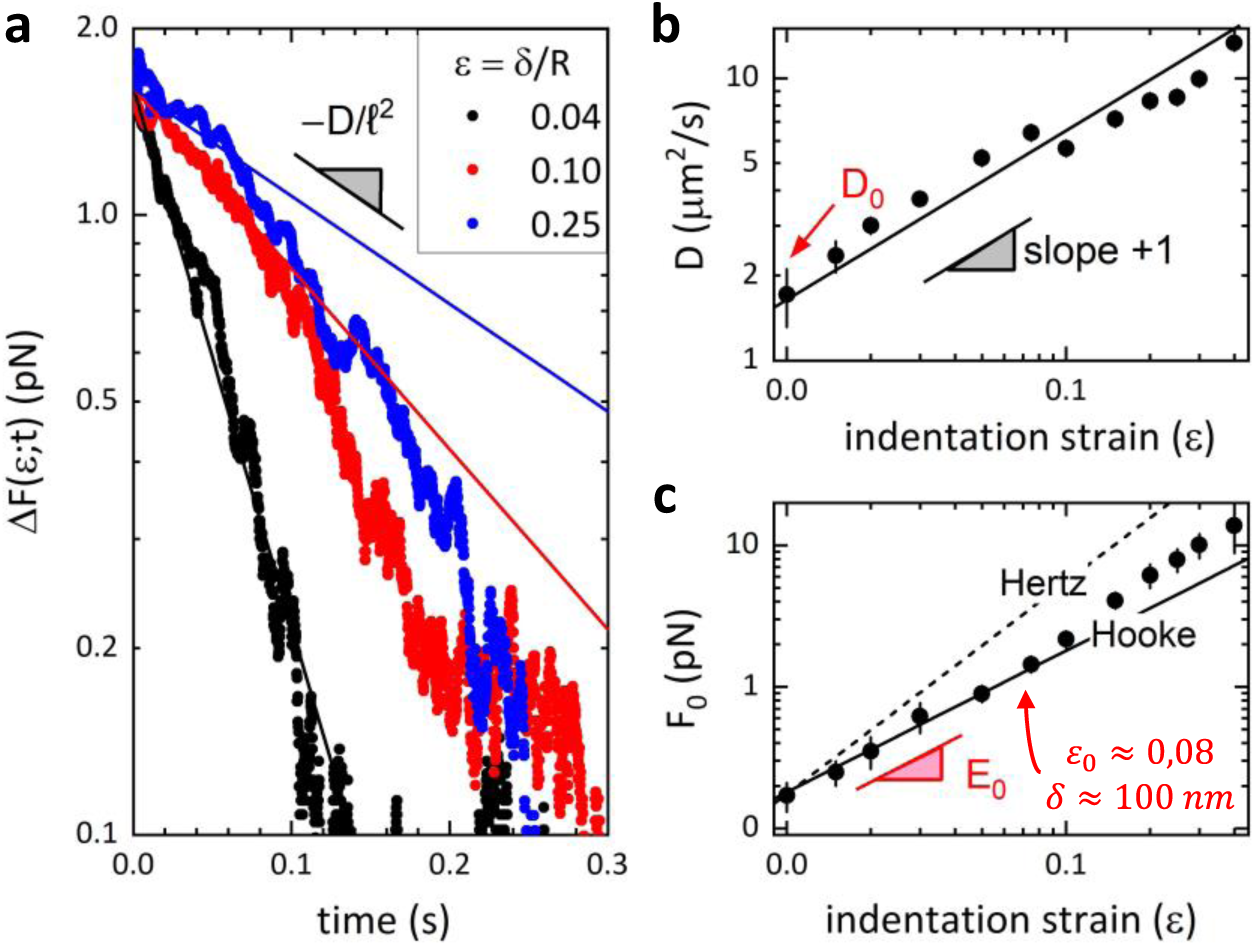
Poroelastic analysis for the lateral OT-indentation of a T3M4 cell. **a** Permeability relaxation plots for different indentation strain (the symbols represent experimental data and the straight lines the best fits to the apparent exponential decay in Eq. 1); **b** Apparent diffusion coefficient as calculated from the initial slope of the relaxation plots for variable indentation strain. The extrapolated zero-strain limiting value represents the bare membrane diffusivity *D*_0_ ≅ (ℓ^2^⁄*τ*_*P*_)_*ɛ*→0_ (see Eq. 2). **c** Cortical stress rigidness calculated as a pure elastic resistance to indentation stress (*F*_0_). At very low deformation below a hardening strain (*ɛ*_0_), we detect the Hookean domain 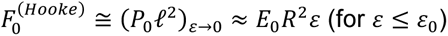, which corresponds to linear shell-like elasticity controlled by the contact pressure describable by an apparent linear Young modulus *E*_0_ → *P*_0_ (see Eq. 3). The Hertz model overestimates the cortical indentation rigidness as calculated for the limiting Young modulus (i.e., 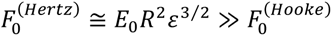.

**Supplementary Table 1:**
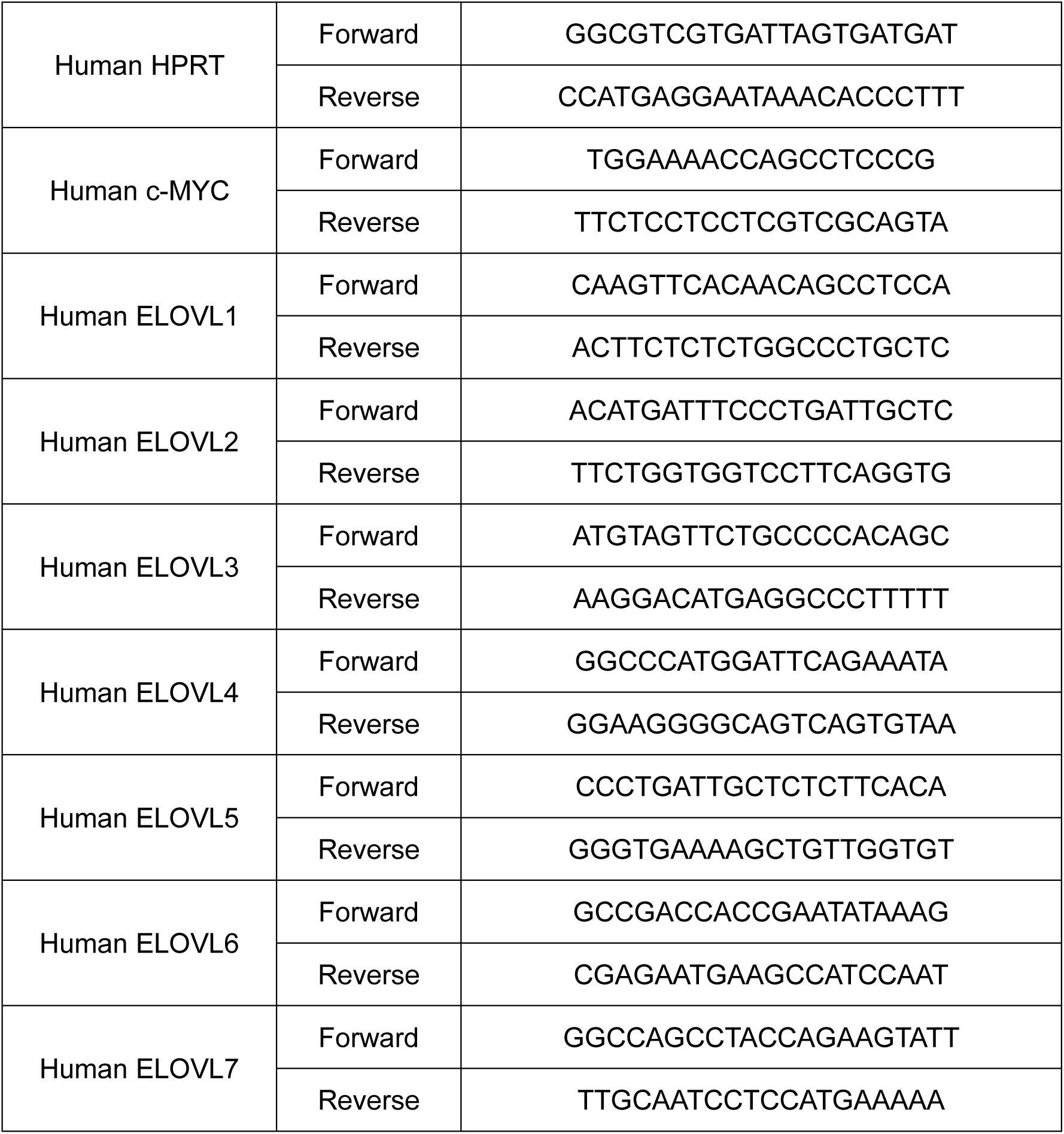
Primers for RT-qPCR.

**Supplementary Table 2:** Antibodies for western blot.

| **Antibody** | **Species reactivity** | **Source** | **Clone** | **Reference** | **Dilution** |
| --- | --- | --- | --- | --- | --- |
| Anti-c-Myc [Y69] | Human, Mouse, Rat | Rabbit | Monoclonal | Abcam, Ref. ab32072 | 1:1000 |
| Anti-ELOVL6 | Human, Mouse, Rat | Rabbit | Polyclonal | Sigma-Aldrich, Ref. PRS4571 | 1:1000 |
| Anti-vinculin | Human, Mouse, Rat, Frog, Canine, Bovine, Chicken, Turkey | Mouse | Monoclonal | Sigma-Aldrich, Ref. V9264 | 1:1000 |
| HRP Anti-beta actin [AC-15] | Human, Mouse, Rat, Hamster, Rabbit, Bovine, Ovine, Canine, African green monkey, Chicken, Guinea pig, Cat, Pig, Carp | Mouse | Monoclonal | Abcam, Ref. ab49900 | 1:25000 |
| Anti-rabbit/HRP | Rabbit | Goat | Polyclonal | Dako, Ref. P0448 | 1:5000 |
| Anti-mouse/HRP | Mouse | Goat | Polyclonal | Dako, Ref. P0447 | 1:5000 |

**Supplementary Table 3:**
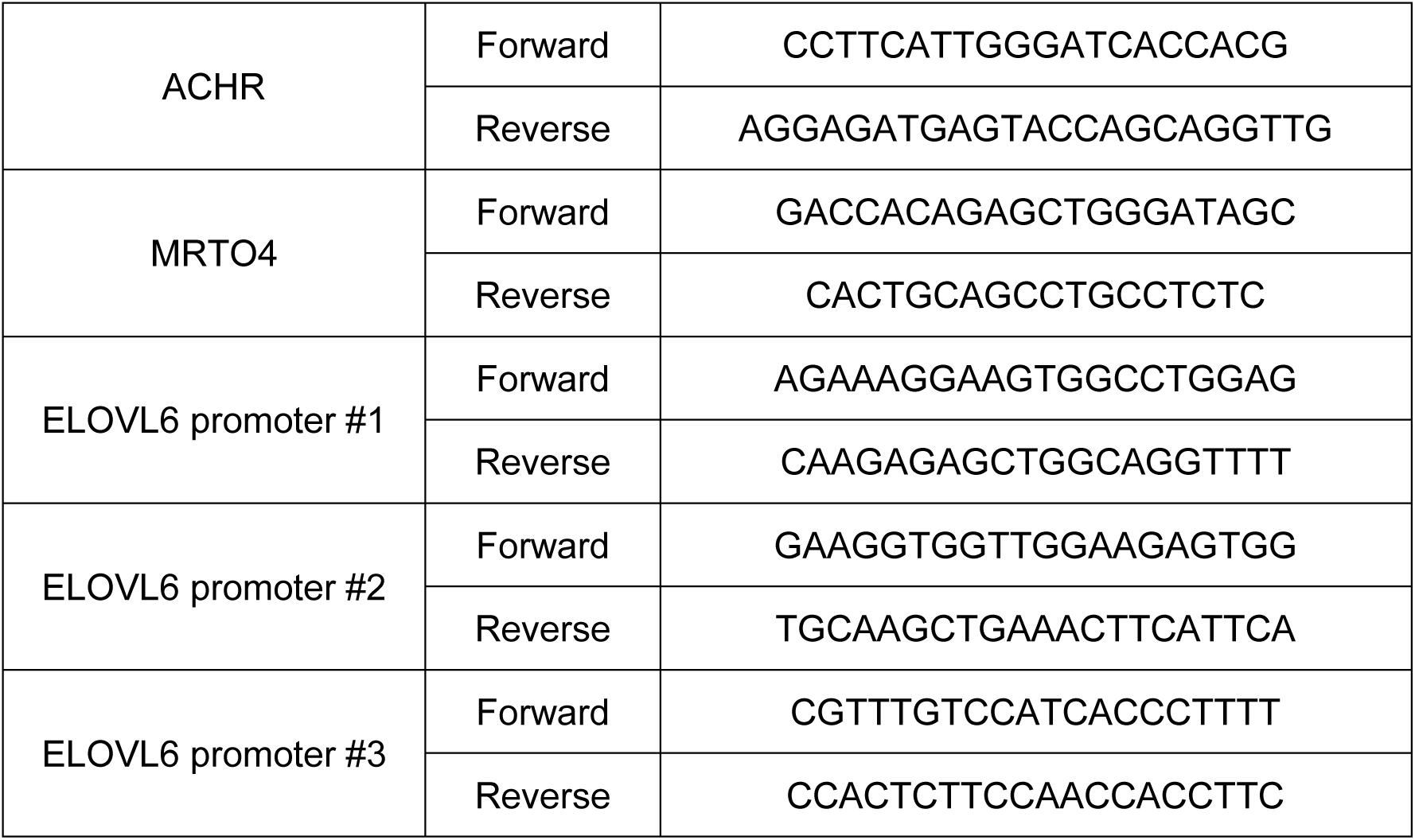
Primers for ChIP.

## Supplementary Note (SN). Lateral indentation using optical tweezers: linear poroelasticity in the shell-like limit

From the seminal 1987’s works by Arthur Ashkin on the trapping of biological objects using infrared radiation^1,2^, optical tweezers (OTs) have become an analytic tool for assessing local stiffness in mechanical indentation contacts with elastic media^3,4^. Advancements in OT-indentation have accomplished the micromechanical analysis of soft materials^5,6^ and living cells^7–10^, even under pathological conditions e.g. in cancer^11^. OT-indentation geometries affordable with spherical microbeads have the advantage to enable stablishing deformation contacts under well characterized strain and minimal invasiveness^5,12^. By analogy to AFM nanoindentation^13–16^, the most usual vertical OT-indentation geometry has been exploited in cell mechanics approaches with adherent cells sticked to the substrate^14,17^. Vertical OT-indentation allows measuring cortical stiffness without employing any retaining element although somewhat affected by the bead-substrate interaction^11,18^. In this work, we have instead adopted the lateral (horizontal) OT-geometry based on a fourfold anvil supported by cell-fixing traps using the forcing one to exert the lateral stress along the horizontal plane (**Fig. SN1**).

### Horizontal (lateral) OT-indentation method and interpretative poroelastic framework

Henceforward, an oscillatory indentation bead is considered to laterally induce cortical deformations along the horizontal direction being the system of interest minimally affected by the rigid substrate (**Fig. SN1a-b**; see caption for experimental details). To perform small indentation deformation in the cellular contact the most flattened as possible (**Fig. SN1a**), we implemented the horizontal OT-geometry with large trapping beads as compared with the inflated cells considered (the bead radius *R* = 5.7 *μm*) (**Fig. SN1b**). Then, the indentation depths *δ* were progressively varied in the reversible regime of relatively small deformations (*δ* ≪ *R*), which define the linear domain at low indentation strain *ɛ* ≡ *δ*⁄*R* ≪ 1 and small contact area ℓ^2^ ≅ *δR* ≈ *R*^2^*ɛ* ≪ *R*^2^ (**Fig. SN1c**). Therefore, the horizontal (lateral) indentation setting allows measurements of plasma membrane and cell cortex properties determined at small deformation strain under minimal or absent mechanical influence from the adhesive substrate^19,20^. For the slow oscillatory indentation performed at variable lateral strain (0 ≤ *ɛ* ≡ *δ*⁄*R* ≪ 1; see also Methods in the main text), the time-force traces exhibited a reversible poroelastic material response sensed into the mechanical surface contact *F*(*ɛ*; *t*) = Δ*F*(*ɛ*; *t*) + *F*_0_(*ɛ*)^21,23^.

**Cell poroelasticity** assumes a mechanical response arising from a porous solid meshwork (plasma membrane, cortex, cytoskeleton, organelles, macromolecules, etc.) bathed by an interstitial fluid (cytosol) considered at osmotic equilibrium with the extracellular medium (or physiological buffer)^23^. Hence, the observable poroelastic response is composed by an initial pressure drop due to solvent permeability Δ*F*(*ɛ*; *t*), then followed by the static resistance force *F*_0_(*∈*) due to the cell rigidity (**Fig. SN1d**). Despite the Hertz model is the most often used to interpret effective indentation rigidness in mechanical surface contacts^4,12^, we have considered a simpler Hookean scheme stablished upon an infinitesimal surface strain defined in a soft medium effectively considered poroelastically homogeneous^13^. In this framework, the viscoelastic properties of the composite membrane-cell cortex system are a manifestation of both the time needed for extracellular fluid intake through of the permeable membrane followed by the redistribution of intracellular fluids in response to applied mechanical stresses and the response of the cell to indenting force application^23^. The poroelastic model depends on two single experimental parameters, the contact pressure *P*_0_(*ɛ*), and the poroelastic diffusion coefficient *D*(*ɛ*)^21,23^. They recapitulate the higher cortical pressures and the faster membrane diffusion expected for the indentation stress relaxations appeared within the increasing larger deformation across the Hookean domain^21,23^. Therefore, the initial fluid overshoot through the porous plasma membrane and the consequent cytosol permeability can be nearly described by an effectively diffusive relaxation function^21,23^:

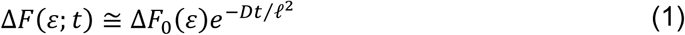

where the strain-dependent poroelastic strength Δ*F*_0_(*ɛ*) ≡ Δ*P*_0_ℓ^2^ ≈ *P*_0_ℓ^2^is determined by the indentation cross section ℓ^2^(*ɛ*) ≈ *πR*^2^*ɛ*, and the contact pressure *P*_0_(*ɛ*) (i.e., the maximal cortical value of the pressure gradient caused under indentation does equilibrate with the osmotic pressure at the cell interior Δ*P*_0_ ≈ *P*_0_ ^21,23^). In poroelastic relaxation terms, the near exponential decay in Eq. 1 is described by an effectively diffusive rate characterized by the apparent diffusion coefficient^23^:

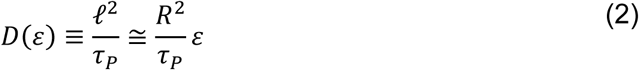

where *τ*_*P*_ is the diffusion time due to fluid permeability in the poroelastic medium^21^. Those poroelasticity parameters can be inferred from lateral indentation experiments (**Fig. SN2**).

**Contact membrane mechanics** frequently assumes the Hertz model for evaluating cortical elasticity in inflated cells under turgor pressure e.g., in plant cells with a rigid wall^16,24^. However, some research groups have indicated that cell wall deformation cannot be described by the pure Hertz model in microindentation experiments^24–27^. An alternative model is a contact model based on elastic shell theory^28,29^, in which the surface cell is assumed to be a thin curved membrane pushed out by inflation pressure^30,31^. We formulate the surface stiffness as an effective Young’s modulus *E*_0_ calculated from the shell-like Hookean theory (corresponding to apparent contact pressure *P*_0_ in the Hertzian theory). Here, we follow the theory of weakly deformed shallow shells under spherical indentation strain (*ɛ* = *δ*⁄*R* ≪ 1), resulted from a small flexural deformation of small amplitude (*δ*), exerted with a spherical bead of large radius (*R* ≫ *δ*) ^28^, the Hookean force results hence determined by the contact pressure (*P*_0_), and the local mean curvature (*R*^−1^) as:

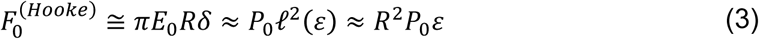

which arises from the effective linear elasticity of the cortical contact with a Young’s modulus *E*_0_ ≡ 12 *B*(1 − *v*^2^)⁄*h*^3^ (including membrane bending rigidity *B*, Poisson ratio *v* and shell thickness *h*)^32^.

Although the shell theory of Hookean spherical indentation infers the apparent membrane stiffness from the flexural bending elasticity and internal pressure^28,29^, the unified formula in Eq. 3 expresses the indentation resistance as a contact pressure equivalent to an effectively linear Young modulus arising from flexural (curvature) rigidness^32^. In our analysis of membrane elasticity based on the elastic shell theory, we further exploited this formula to infer the apparent membrane stiffness observed from the lateral OT measurement extrapolated to zero-limiting strain. Conversely, the Hertzian force in a spherical indentation contact with a bulky elastic medium is predicted with the canonical formula^33^:

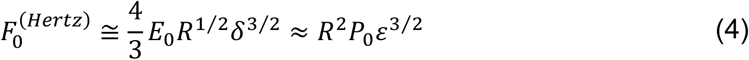

### Experimental data analysis

In **Fig. SN2** we showcase a typical poroelastic analysis as performed in this work. For the sake of example, we present the kind of permeability relaxation plots corresponding to the dynamic indentation forces Δ*F*(*ɛ*; *t*) recorded for three different values of the deformation strain (small *ɛ* = 0.04, medium *ɛ* = 0.10 and large *ɛ* = 0.25; **Fig. SN2a**). The apparently linear values of the Young modulus (*E*_0_) and diffusivity (*D*_0_) can be estimated from the limiting slope of the applied force as measured for variable indentation depths in the Hookean regime^13,21^. On the one hand, the initial diffusive slope is determined from the best fittings to the exponential relaxation decay (see Eq. 1). Hence, the apparent permeability diffusivities *D*(*ɛ*) can be calculated using Eq. 2 (**Fig. SN2b**). The experimental estimations for the diffusion coefficient exhibit the same linear dependence as predicted from the poroelastic model on the indentation strain^21–23^. Hence, the bare diffusivity of the membrane can be extrapolated down to zero-limiting deformation i.e., 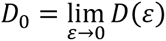 (**Fig. SN2b**). On the other hand, we analyse the resistance of the plasma membrane (and underlying cortex) to spherical indentation using the Hookean model (Eq. 3). Hence, the apparent values of the Hookean Young’s modulus *E*(*ɛ*) of the cell surface can be calculated by reference to the usual Hertzian dependence (Eq. 4), which overestimates the indentation stress as corresponding to the response of a bulk material (**Fig. SN2c**). We detect the linear elastic regime at strain below a stiffening point (*ɛ*_0_ ≈ 0.08), corresponding to a small indention depth (*δ* ≈ 100 *nm*). Hence, the bare Young’s modulus consistent with the flexural rigidity of the membrane can be estimated as 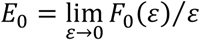 (**Fig. SN2c**).

